# Tracing the vertebrate selenoproteome evolution reveals expansions in ray-finned fishes and convergent depletions in tetrapods

**DOI:** 10.1101/2025.05.29.656587

**Authors:** Max Ticó, Jesus Lozano-Fernandez, Marco Mariotti

## Abstract

Selenoproteins incorporate the rare selenium-containing amino acid selenocysteine (Sec) and play crucial roles for redox homeostasis, stress response, and hormone regulation. Sec is inserted by co-translational recoding of the UGA codon, normally a stop. As a consequence, selenoproteins are often misannotated in public databases and require specialized bioinformatic methods and resources.

Here, we present a refined characterization of the composition and evolution of the vertebrate selenoproteome. Based on analyses of 19 gene families across hundreds of genomes, we show that extant selenoproteomes were shaped by extensive gene duplications (56 selenoproteins), losses (50), and Sec-to-cysteine (Cys) conversions (21). Tetrapods including mammals encode 24-25 selenoproteins, with variations in 6 families. Notably, the same genes underwent convergent evolutionary events in multiple tetrapods, namely Sec-to-Cys substitutions (*SELENOU1*, *GPX6*) and gene losses (*SELENOV*). In contrast, ray-finned fish exhibit larger and more dynamic selenoproteomes, reinforcing the hypothesis that the selective advantage of Sec is stronger in aquatic environments. We detected selenoprotein duplications spread across the actinopterygian clade involving 13 families, mainly involved in antioxidant defense. The richest selenoproteomes were found in Salmonidae and Cyprinoidei fish with 56 and 44 selenoproteins, respectively, owing to whole genome duplications. Among our findings, the *SELENOP* family stands out in lampreys, carrying up to an unprecedented 162 UGAs putatively recoded to Sec.

Our study presents the most comprehensive evolutionary map of vertebrate selenoproteins to date and delineates the specific selenoproteome of each lineage, establishing a foundational framework for selenium biology research in the era of biodiversity genomics.

## BACKGROUND

Selenium is an essential trace element for human health. The biological effects of selenium are mostly mediated by its incorporation in the non-canonical amino acid selenocysteine (Sec) [1–3]. Sec is the essential component of selenoproteins, a unique class of proteins with crucial functions in both normal physiology and cancer. Sec is often referred to as the 21st amino acid due to its distinctive biosynthetic pathway and incorporation mechanism, which differ fundamentally from those of the 20 canonical amino acids. Importantly, Sec is inserted in correspondence to the UGA codon, which is normally a stop codon. The UGA is co-translationally recoded to Sec in selenoprotein mRNAs in response to local sequence elements, the major of which is called the SECIS (selenocysteine insertion sequence) [4, 5]. Moreover, unlike standard amino acids which are synthesized then charged onto their cognate transfer RNAs (tRNAs), Sec is synthesized directly on its own dedicated tRNA, tRNASec, through a multi-step enzymatic process. The selenium required for this reaction is provided in the form of selenophosphate, synthesized by protein SEPHS2, an essential enzyme in Sec metabolism which, intriguingly, is itself a selenoprotein [6].

Since their discovery in the 1970s, Sec and selenoproteins have been extensively studied across diverse fields of biology [7, 8]. Indeed, despite representing a small fraction of protein-coding genes, selenoproteins perform many critical functions in redox homeostasis, stress response, hormone maturation, and protein quality control, thus playing key roles in both physiology and cancer biology. This has prompted extensive research into their enzymatic functions, selenium metabolism, and the unique mechanism of Sec biosynthesis. Selenium itself, a trace element with a narrow margin between deficiency and toxicity, has also drawn interest in nutritional and physiological contexts [3]. As a result, selenoproteins are investigated in a broad range of species, from standard model organisms (e.g., *Mus musculus, Danio rerio*) to agriculturally relevant animals (e.g., *Bos taurus, Gallus gallus, Salmo salar*). Naturally, understanding the full set of selenoproteins encoded by an organism is essential for the meaningful design and interpretation of experiments to elucidate selenium biology.

The human and murine selenoproteomes were first described in 2003 as comprising 25 and 24 Sec-encoding genes, respectively [9]. Subsequent studies expanded the known selenoproteomes of selected vertebrates and metazoans [10–15]. In our seminal paper in 2012, we provided the first description of selenoproteome composition and evolution across diverse vertebrates [16]. Although highly influential, that study was constrained by the limited availability of sequenced genomes at the time (44).

In the last 10 years, major advances in sequencing and the rise of biodiversity genomics provided an unprecedented wealth of genomic data [17, 18]. However, due to their deviation from the standard genetic code, selenoproteins have not kept pace with the expansion of automated genomics resources. At present, ∼90% and ∼50% of selenoprotein genes have substantial annotation errors in Ensembl and NCBI genomes, respectively [19]. This also led to unreliable selenoprotein data in evolutionary resources dedicated to orthology (e.g. Ensembl Compara [20], EggNOG [21]).

In this manuscript, we present a refined comprehensive characterization of the selenoproteome of vertebrates, obtained by meticulously tracing the evolution of 19 protein families across hundreds of genomes through sequence analysis and phylogenetic reconstruction. Our work provides an essential evolutionary framework to understand the selenoprotein content of any vertebrate genome, expanding on previous efforts by an order of magnitude.

## RESULTS

We downloaded 314 genome assemblies from Ensembl v.108 representative of all major lineages of vertebrates, complemented by 26 NCBI genomes from early vertebrates and outgroups (Methods). Next, we used the software Selenoprofiles (Methods) to identify homologs belonging to any of the known metazoan selenoprotein families in those genomes, resulting in 13,517 gene predictions. Phylogenetic reconstruction was then performed for each family, and gene and species trees were manually inspected to infer gene evolutionary events (Methods). These comprise gene duplications, gene losses, and the selenoprotein-specific event of the conversion of a Sec residue to Cys. In selenoprotein evolution, Cys to Sec events are also possible, but have been observed only along evolutionary distances much larger than encompassed by vertebrates [22, 23]. Each branch of gene trees was manually annotated as a separate orthologous group, assigning gene names as consistent as possible with existing resources and literature. Tree inspection and orthology assignments were performed taking into account branch support values obtained by UltraFast Bootstrap (UFB). The great majority of orthologous groups were supported by high UFB support values in gene trees (Methods); all exceptions are explicitly mentioned in their relevant section. Each putative event was critically assessed, following criteria of maximum parsimony, and seeking consistency with the species tree and support by independent data (e.g. RNA sequences, other genome assemblies from NCBI) (Methods).

Thanks to the vast diversity of organisms in our dataset, the great majority of events were shared among multiple species, leading to phylogenetically coherent patterns of gene occurrence that allowed reliable evolutionary inferences. Yet, two lineages stood out for their divergence from this general trend. First, bird genomes exhibited many apparent gene losses scattered across lineages, in a pattern that would imply dozens of independent events (Supplementary Figure S1). This included the apparent absence of *SEPHS2*, a gene that nevertheless is required for biosynthesis of all other selenoproteins [22], and of *SELENOW*. We ascribed this pattern to a known bias of bird assemblies: they systematically lack certain GC-rich regions that present technical challenges for sequencing [24]. To further support this interpretation, we examined the syntenic block surrounding *SEPHS2*, which includes the essential genes *G6PD* and *IKBKG* (Figure S2). This genomic region is conserved across Sauria; yet, we observed the apparent absence of all three genes in most (but not all) Aves. In contrast, outgroup lineages such as Testudines retain the complete syntenic block, including the three genes. Strikingly, we found RNA evidence of expression for both genes (*SEPHS2* and *SELENOW*) in multiple species despite their apparent absence from their available genome assemblies, demonstrating that these genes are present in the actual genomes of birds. Ultimately, we dismissed all selenoprotein losses in Sauria (which includes birds) as artifactual.

Second, the fish lineage of Cyprinoidei (e.g. carps, *Danio rerio*) exhibited a massive amount of apparent gene duplications (Supplementary Figure S1), which we could not precisely disentangle by phylogenetic analysis. We realized that the sublineage of Cyprininae (e.g. carps) reportedly underwent extensive recent whole genome duplications and polyploidy, which greatly complicated this task. Therefore, we decided to limit our analysis to the evolutionary events that occurred at the root of Cyprinoidei, i.e. shared between *D. rerio* and Cyprininae, and dismiss the rest.

Several of the outgroups we used have a characterized selenoproteome: *Ciona intestinalis* [15], *Branchiostoma floridae* [14], *Drosophila melanogaster* [25], *Strigamia maritima* [26, 27]. This allowed us to critically assess their gene predictions, and compare them with those of vertebrates to phase evolutionary events, as detailed later for each family.

### Overview of the vertebrate selenoproteome

We identified selenoproteins belonging to 19 protein families. The most represented families were glutathione peroxidases with 1,529 Sec-encoding genes, iodothyronine deiodinases with 988, and thioredoxin reductases with 718. In general, Actinopterygii (ray-finned fishes) exhibited richer selenoproteomes (35-56 selenoprotein genes per genome) than Tetrapoda (24-25 genes per genome) (Figure 1).

**Figure 1.**
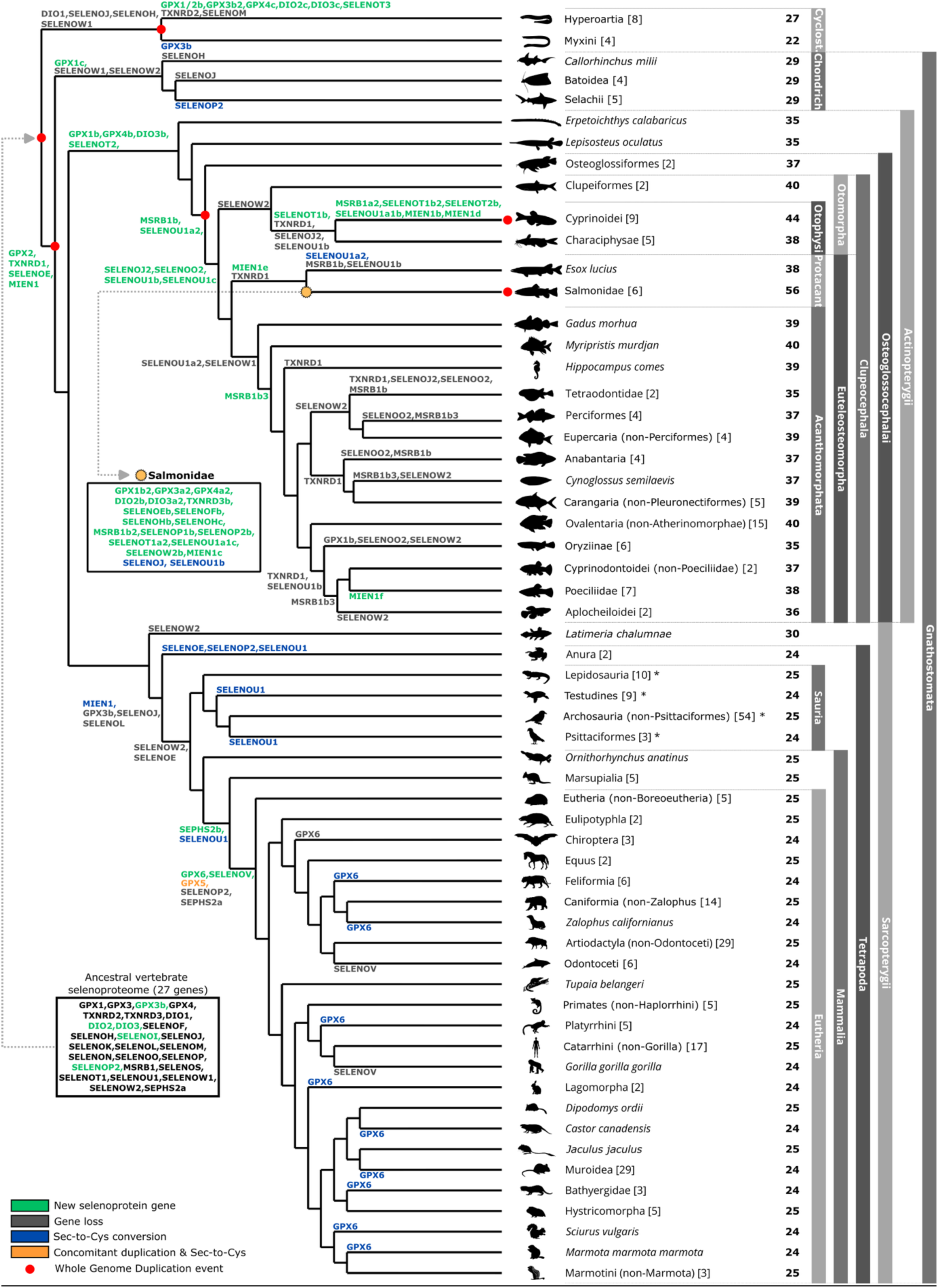
Evolution of the vertebrate selenoproteome. The evolutionary events shaping selenoprotein genes are shown along the species tree. Leaf nodes represent either individual species (italicized) or collapsed lineages; in the latter case, the number in brackets indicates how many species from that lineage are included in our main dataset. Asterisks (‘*’) denote lineages with apparent gene losses that we interpret as artifacts caused by known issues in genome assembly quality. Bold numbers in the rightmost column indicate the number of selenoprotein genes detected in each species or lineage. Grey rectangles highlight taxonomic clades of interest; “Cyclost.” Stands for Cyclostomata, and “Protacant” for Protacanthopterygii. Red dots mark whole genome duplications (WGD) reported in literature [74–79]. Animal silhouettes were sourced from https://www.phylopic.org.

While our analyses recapitulated earlier reports performed at a smaller scale, we also detected many novel events, particularly gene duplications. The majority affected Actinopterygii and its sublineages, whose selenoproteome appeared not only larger, but also more dynamic. In total, we detected 56 gene duplications (37 in Actinopterygii), and 50 gene losses (29 in Actinopterygii). The most selenoprotein-rich organisms were Salmonidae (aka salmonids, e.g. trouts, salmons) with 56 selenoprotein genes, 17 of which emerged by duplication specifically in this lineage, and Cyprinoidei with 44 selenoproteins, 6 of which lineage-specific. On the other hand, Sec-to-Cys conversions were more prevalent in Tetrapoda: we detected 21 of these events overall, 16 of which were in Tetrapoda. Hereafter, we describe the selenoproteome evolution for each family separately.

#### Glutathione peroxidases (GPX)

Glutathione peroxidases (GPX) are key antioxidant enzymes that protect cells from oxidative damage by reducing hydrogen peroxide and lipid peroxides using glutathione as an electron donor [16, 28]. Mammals possess eight *GPX* homologs with specialized tissue distribution and functions, including five selenoproteins: *GPX1* (ubiquitous), *GPX2* (gastrointestinal), *GPX3* (extracellular, kidney), *GPX4* (membrane-associated, ubiquitous), *GPX6* (olfactory system); and three Cys paralogs: *GPX5*, *GPX7*, and *GPX8*. Our phylogenetic analysis (Figures 2 and 3, Supplementary Figure S3) confirms a well-supported evolutionary classification of GPX proteins, identifying three major groups: GPX1/GPX2, GPX3/GPX5/GPX6, and GPX4/GPX7/GPX8. These findings align with previous studies on GPX evolution [16, 28, 29].

**Figure 2.**
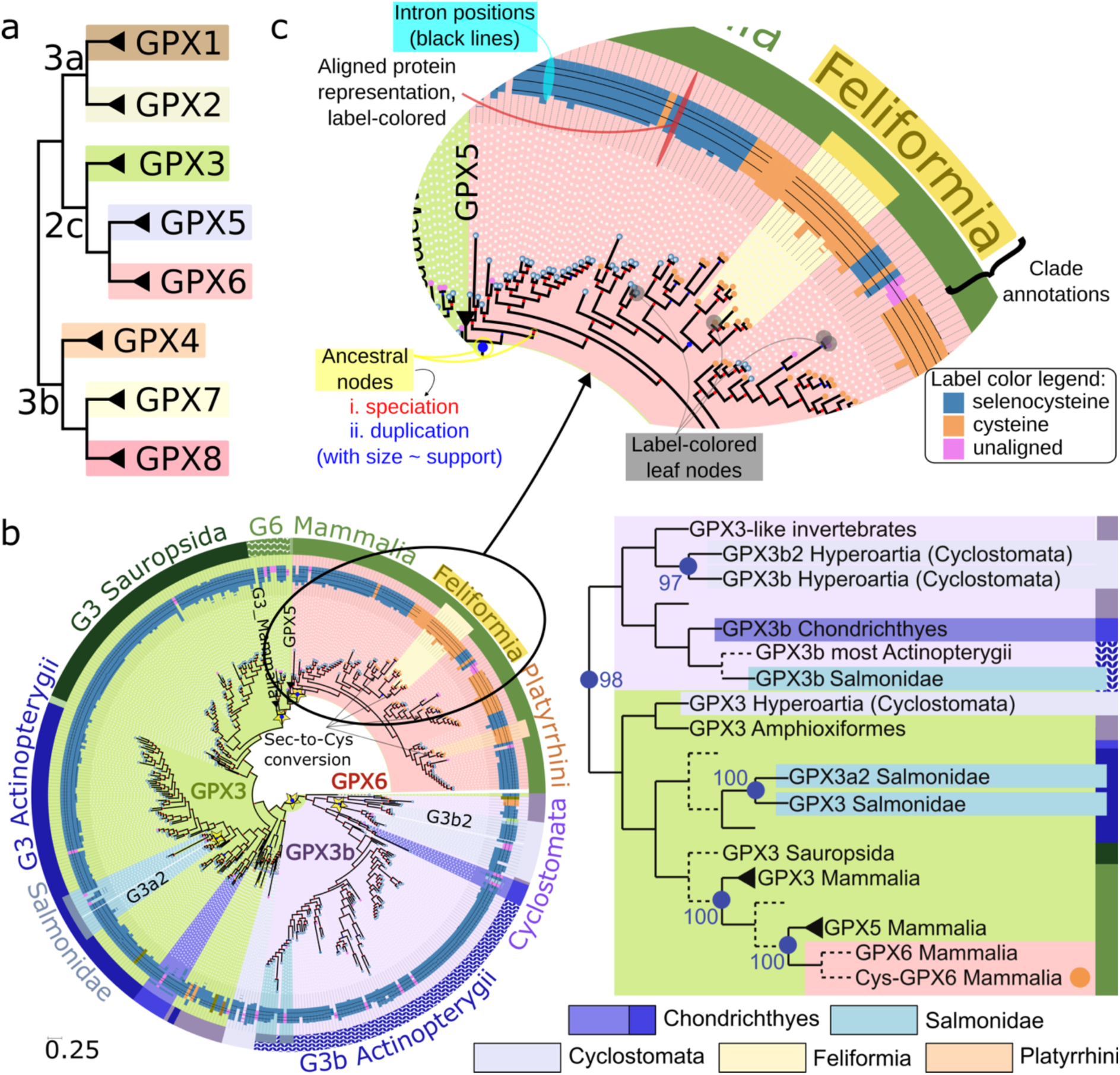
Phylogeny of the GPX family. The GPX gene tree was computed collating all sequences of this family, but it is split in different representations for visualization purposes. **a)** Topology of the GPX family, split into three clades shown in panels b, Figure 3a and 3b. **b)** Phylogeny of GPX3/5/6 clade. To the right of the main tree, a summarized tree representation is shown, displaying the evolutionary events identified in the full tree, with selected clades collapsed for visualization. Dotted lines in the summary tree represent paraphyletic, non-collapsible branches. Duplication events are indicated by blue nodes labeled with their UFB support values. The legend below shows the color scheme used to represent the taxonomic groups of the species highlighted in the tree. Note that certain monophyletic clades of the gene tree were collapsed (e.g. see GPX5). The full expanded (non-collapsed) gene tree is available as Supplementary Figure S3. **c)** Zoomed section of the GPX6 gene tree, used here to illustrate this form of representation, abundantly used throughout this paper. Each leaf corresponds to a putative gene whose protein sequence was included in the tree. Leaves are color-coded by their label, assigned by Selenoprofiles by examining which codon aligns to the Sec position in the profile (e.g. blue for Sec-UGA, orange for Cys, pink for unaligned). Moving outward, you will see an equally colored representation of the corresponding aligned protein, which shows its non-gap positions and includes black lines to map where introns are located in the coding sequence. Most intronless genes are pseudogenes, which constitute the prevalent false positives of Selenoprofiles. The outmost circle provides a manual annotation of gene clades, with colors that indicate the taxonomy of the source of those genes (e.g. blue for Actinopterygii, green for Mammalia). Internal nodes of the gene tree represent either speciation events (red) or duplication events (blue), determined via the species overlap algorithm. The latter are displayed with size proportional to the number of species supporting the duplication. The nodes that define duplication events deemed genuine via manual analysis are marked with a star. Branch lengths represent the expected number of substitutions per site and were estimated by maximum likelihood under the best-fitting substitution model in IQ-TREE.

**Figure 3.**
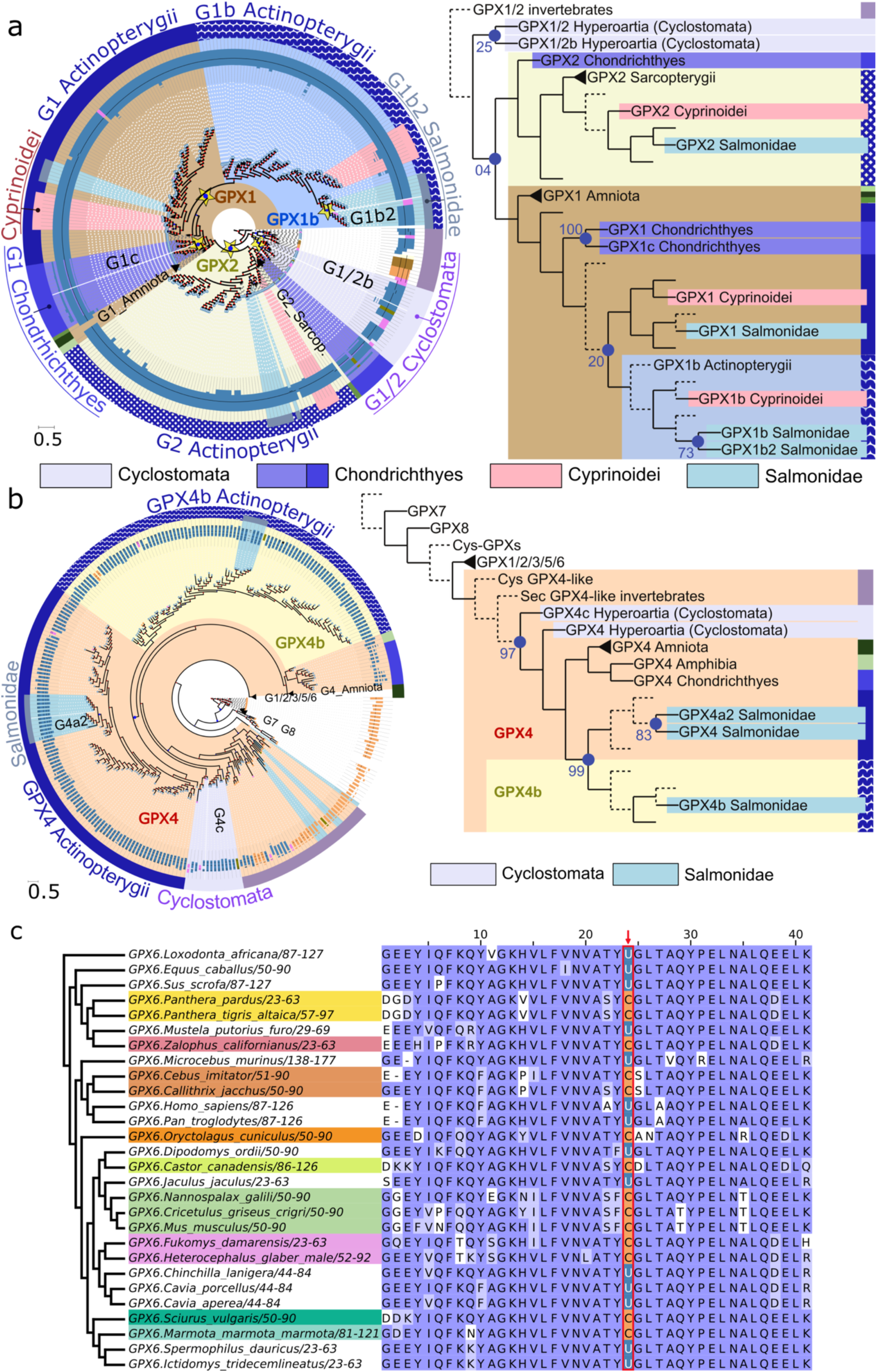
Phylogeny of the GPX family, continued from figure 2. **a, b)** Gene tree visualization of the GPX1/2 and GPX4/7/8 clades, respectively. See full feature description in Figure 2c The full gene tree is available as Supplementary Figure S3. **f)** Multiple sequence alignment of GPX6 proteins including representatives that underwent Cys conversions. Sec residues are shown in blue, while Cys residues at this position are in orange. The identifiers of GPX6-Cys homologs are highlighted.

*GPX1, GPX2, GPX3, GPX4* are present in all vertebrates, with a single notable exception. In the GPX1/GPX2 group, we observed that the Hyperoartia class of jawless fish (Cyclostomata superclass) possessed two similar genes which phylogenetic reconstruction placed as outgroups of the GPX1 and GPX2 clusters of other vertebrates (Figure 3a). We then noticed that the other class of Cyclostomata, Myxini, has instead a single orthologous gene (Supplementary Figure S1). We infer that GPX1 and GPX2 split in Gnathostomata (jawed vertebrates) and that Cyclostomata likely reflect the ancestral status, so that we named this gene GPX1/2 and its Hyperoartia-specific paralog GPX1/2b. Although the UFB support of the corresponding node in the reconstructed protein tree is only 25%, we argue that we can confidently map the origin of GPX1/2b to Hyperoartia based on the species-tree analysis.

*GPX6* originated at the root of placentals and exhibited a very dynamic evolution. Multiple independent *GPX6* Sec-to-Cys conversions had been reported [16], which we confirmed and expanded in our analyses (Figures 1, 2b, 2c, 3c; Supplementary Figure S3). We observed Cys conversions in multiple taxonomic groups, including Platyrrhini primates (New World monkeys, e.g. *Callithrix jacchus*, *Cebus imitator*), Muroidea rodents (e.g. *Mus musculus*, *Rattus norvegicus*, *Mesocricetus auratus*), Lagomorpha (e.g. *Oryctolagus cuniculus*, *Ochotona princeps*), Feliformia (e.g. *Felis catus*, *Panthera pardus*), Bathyergidae rodents (e.g. *Fukomys damarensis*). In addition, we also detected species-specific conversions, namely in *Marmota marmota marmota, Sciurus vulgaris, Zalophus californianus and Castor canadensis* (Figure 3c). Furthermore, *GPX6* was lost altogether in Chiroptera (bats).

We identified several GPX gene duplications across vertebrates. We confirmed two previously reported GPX gene duplications at the root of Actinopterygii leading to the emergence of *GPX4b* and *GPX1b* (the latter with only 20% UFB, but consistent occurrences along the species tree). These genes were found in all investigated species within this lineage, except for Oryziinae fish (e.g. *Oryzias latipes*), which we infer to have lost *GPX1b*. In two Oryziinae species, we initially identified a putative Cys-containing *GPX4b* paralog, yet we dismissed it due to lack of sequence support from RNA data. A third gene, *GPX3b*, was also reported in Actinopterygii; our analysis indicates that it originated at the root of Vertebrata, and was subsequently converted to Cys in Myxini and lost in Tetrapoda.

We also observed additional selenoprotein gene duplications in specific branches of Actinopterygii. In Hyperoartia, we observed three lineage-specific events that yielded *GPX1/2b*, *GPX3b2* and *GPX4c*. Then, a lineage-specific duplication of *GPX1* gave rise to a distinct Sec-paralog, *GPX1c*, in Chondrichthyes (cartilaginous fish). We mapped three additional duplications in Salmonidae yielding *GPX1b2*, *GPX3a2*, and *GPX4a2*. All these events were supported by gene tree nodes with high UFB support values (>70%), except the aforementioned *GPX1/2b.* Notably, these duplications were previously reported in an analysis of 8 fish genomes [29]. Our study demonstrates that they are lineage-specific events, so that we have assigned new names to the duplicated genes to reflect their precise evolutionary trajectories. The correspondence with [29] is provided in Supplementary Table T1.

#### Iodothyronine deiodinases (DIO)

Iodothyronine deiodinases (DIO) regulate thyroid hormone metabolism by catalyzing the activation or inactivation of thyroxine (T4) and triiodothyronine (T3) [30]. The vertebrate DIO family includes three main genes: *DIO1*, which is involved in peripheral thyroid hormone conversion; *DIO2*, which plays a critical role in local T3 activation; and *DIO3*, which inactivates excess thyroid hormones, preventing hyperthyroidism. Through our analyses (Figures 1, 4, Supplementary Figures S1, S4), we detected these three genes in all vertebrate lineages, with the exception of Cyclostomata that lacked *DIO1*. This gene is present in all investigated Chordates, indicating that *DIO1* predates the vertebrate radiation and was then lost in Cyclostomata. All Actinopterygii additionally carried a *DIO3* duplicate (*DIO3b*). The placement of *DIO3b* genes in the reconstructed protein tree was mostly consistent with an origin by *DIO3* duplication at the root of Actinopterygii (UFB support of relevant node: 68%), with the exception of Cyprinoidei *DIO3b* which was placed closer to the tree root, possibly due to long branch attraction.

**Figure 4.**
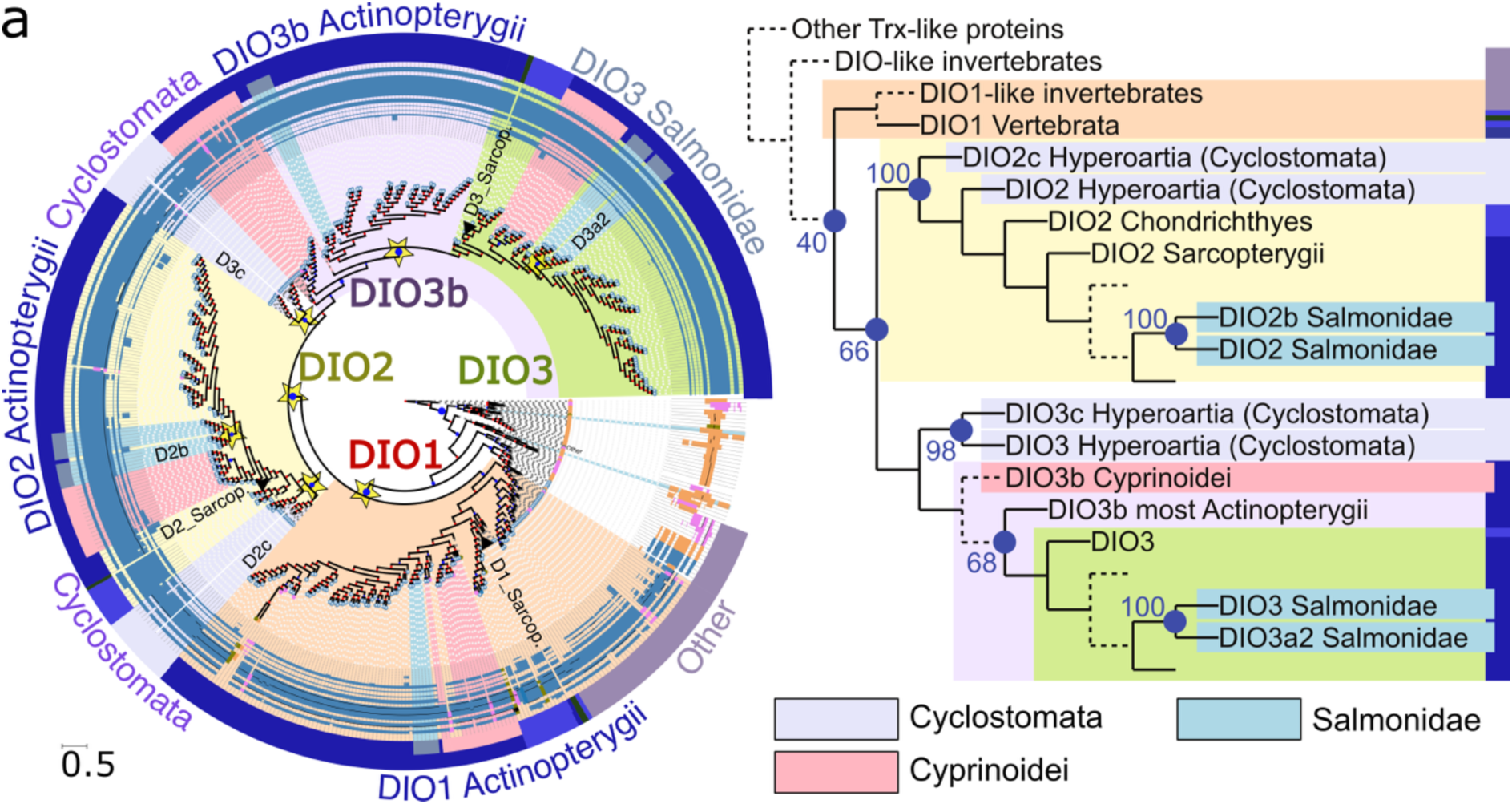
Phylogeny of the DIO family. **a)** Gene tree visualization of the DIO family. See full feature description in Figure 2c. Expanded gene and species trees are available as Supplementary Figures S4 and S5.

We also confirmed two more duplications, one of the ancestral *DIO3* gene (also called *DIO3a*) [31], to generate *DIO3a2*, and the other of *DIO2* to yield *DIO2b* (Figures 1, 4, Supplementary Figures S4, S5). Both were mapped to Salmonidae with high support, as previously reported [31]. In Hyperoartia (Cyclostomata), we identified two lineage-specific duplications resulting in *DIO2c* and *DIO3c* (absent from Myxini, the other class of Cyclostomata; Supplementary Figure S1). Both were highly supported, with the caveat that *DIO2c* was placed as an outgroup of all *DIO2* genes, rather than at its expected position as sister of Cyclostomata *DIO2*.

#### Thioredoxin reductases (TXNRD)

Thioredoxin reductases (TXNRDs) play a central role in maintaining cellular redox balance by catalyzing the reduction of thioredoxins, which are essential for regulating oxidative stress responses, cell proliferation, and redox-regulated signaling cascades [32]. Tetrapods have three TXNRD genes: *TXNRD1* (cytosolic), *TXNRD2* (mitochondrial), and *TXNRD3* (also known as *TGR*; testis-specific, containing a peculiar N-terminal Grx domain), each of which participates in distinct physiological processes. In phylogenetic reconstruction, *TXNRD1* and *TXNRD3* clearly cluster together, indicating a shared origin.

Reportedly, fishes lack the *TXNRD3/TGR* gene, but their *TXNRD1* gene encodes for a Grx-containing protein as its major isoform, which duplicated to give rise to TXNRD3/TGR in tetrapods [16, 33]. Yet, our present analysis (Figure 5, Supplementary Figures S6-S9) did not recapitulate TXNRD evolution as previously proposed. While indeed nearly all fish possess a single gene in the *TXNRD1/TXNRD3* group, we claim here that it is the ortholog of *TXNRD3*, rather than *TXNRD1*. This classification is supported by both the topology of reconstructed gene trees (Figure 5a, Supplementary Figure S6) and by syntenic conservation (Supplementary Figures S7, S8). Furthermore, we detected both genes (*TXNRD1* and *TXNRD3)* in Chondrichthyes. Because this lineage serves as an outgroup to both Actinopterygii and Tetrapoda, we argue that *TXNRD1* emerged earlier than tetrapods. To explain its sparse occurrence across Actinopterygii, we hypothesize that it was subsequently lost independently in six Actinopterygian lineages (Figure 1). Curiously, fish *TXNRD1* features an atypical GNCUG C-terminus (rather than GCUC found across TXNRDs). We did not find *TXNRD1* in Cyclostomata, and thus we map the *TXNRD1/TXNRD3* duplication to the root of Gnathostomata. Finally, because the Cyclostomata gene clearly clusters with *TXNRD3*, we identify the *TXNRD3* clade as the ancestral one and *TXNRD1* as the gene derived from its duplication.

**Figure 5.**
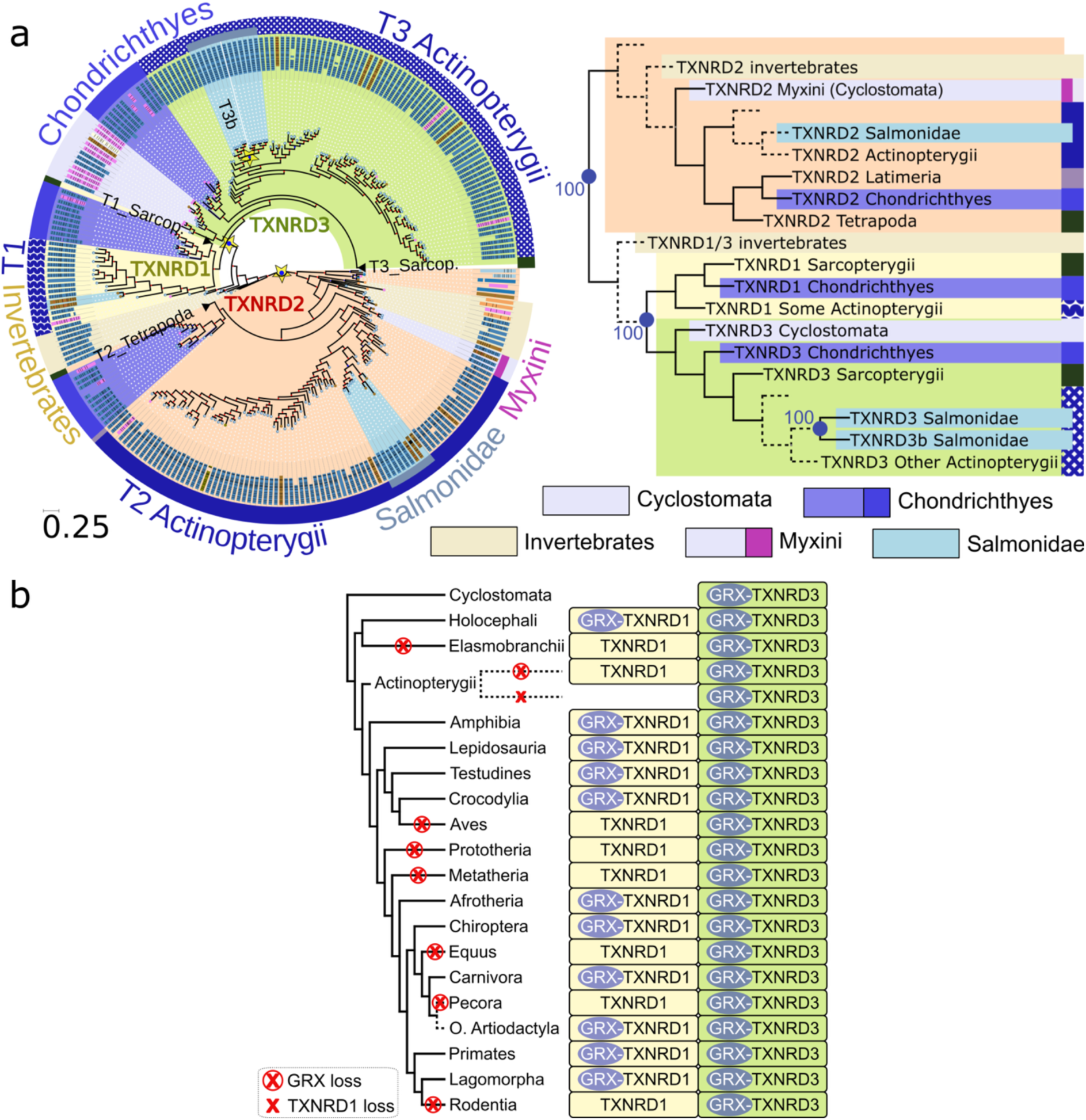
Phylogeny of the TXNRD family. **a)** Gene tree visualization of the TXNRD family. See full feature description in Figure 2c. Expanded gene and species trees are available as Supplementary Figures S6, while Supplementary Figures S7 and S8 contain analyses of the synteny of TXNRD1 and TXNRD3 genes. **b)** Phylogenetic distribution of the Grx domain in TXNRD1 and TXNRD3 proteins across vertebrate lineages. Colors are assigned according to the TXNRD gene tree. A non-collapsed version of this figure is available as Supplementary Figure S9.

It is reported that the human *TXNRD1* gene encodes for a minor transcript isoform that includes the Grx domain (“TxnRd1_v3”), which is a predominant feature of *TXNRD3* genes. This transcript isoform is reportedly absent in *M. musculus TXNRD1* [32]. Therefore, we systematically investigated the occurrence of Grx domains in the genomic regions of *TXNRD1* and *TXNRD3* across all vertebrates (Methods). Our results show that Grx domains are a widespread feature of *TXNRD1* genes (Figure 5b, Supplementary Figure S9). However, they have been lost in several tetrapod lineages independently (Rodentia, Aves, Equus, Prototheria, Metatheria and Pecora) and in all Actinopterygii retaining the *TXNRD1* gene, possibly due to functional redundancy with *TXNRD3*. Altogether, we argue that the Grx-domain is an ancestral feature of *TXNRD3* which was initially retained in *TXNRD1*, but that was later lost in multiple lineages through convergent evolution.

We further investigated the ancestral state of vertebrates by examining the position of other metazoan groups in the TXNRD gene tree (Figure 5a), as well as an extended gene tree featuring additional outgroups available in PhylomeDB [34] (QFO2020 seed, *Caenorhabditis elegans* phylome ID: 791). Our phylogenetic analyses indicate that the divergence between *TXNRD1/TXNRD3* and *TXNRD2* likely predates the origin of metazoans, as representatives from both clades are found in diverse animal lineages, including the nematode *C. elegans*, the annelid *Helobdella robusta*, and the cnidarian *Nematostella vectensis*. Secondary events are evident in several lineages, and detailed hereafter. The tunicate *Ciona intestinalis* lost *TXNRD2*: we only detected one TXNRD homolog in this species, consistent with an earlier report [15], which clusters with *TXNRD1/TXNRD3* in the reconstructed gene tree (Supplementary Figure S6). Analogous observations indicate that arthropods such as *Ixodes scapularis* and *D. melanogaster* lost *TXNRD1* (Supplementary Figure S6). Notably, *D. melanogaster* retains two *TXNRD2*-derived genes, presumably the result of a secondary duplication. Fitting with our analysis, non-vertebrate animals carried Grx-domains solely in *TXNRD3* genes. Due to the presence of this domain, these invertebrate genes were referred as *TGR* in literature, despite weak or absent glutathione reductase activity [35]. Altogether, we reconstruct that ancestral vertebrates carried a cytosolic Grx-containing *TXNRD3* gene and a mitochondrial *TXNRD2* gene, the former later giving rise to *TXNRD1* in Gnathostomata.

We further detected one *TXNRD3* duplication occurred in Salmonidae and yielded *TXNRD3b*, which retained the Grx domain. Lastly, the Hyperoartia clade of Cyclostomata lost *TXNRD2* (despite retaining its neighboring genes) and thus encodes *TXNRD3* as their sole gene in this family. *TXNRD2* was detected in its sister clade, Myxini (Supplementary Figure S1).

#### SELENOJ

Selenoprotein J (SELENOJ) shows significant similarity to jellyfish J1-crystallins and, together with them, forms a distinct subfamily within the large family of ADP-ribosylation enzymes. While the homology may suggest a structural role which would be unique among selenoproteins, SELENOJ remains functionally uncharacterized. The *SELENOJ* gene is reportedly restricted to actinopterygian fishes and sea urchins, with Cys homologs reported in cnidarians [11]. Our analyses replicated these earlier findings, and additionally highlighted the presence of *SELENOJ* in Amphioxiformes (Supplementary Figure S10). On the other hand, *SELENOJ* was lost in Cyclostomata and Batoidea (rays e.g *Leucoraja erinaceus*)(Supplementary Figure S1). Moreover, we had previously reported a *SELENOJ* duplication in *O. latipes* and *Gasterosteus aculeatus* [16], yielding *SELENOJ2*. Here, we traced back this duplication to the root of Clupeocephala (76% UFB support), which included more than half of fishes in our dataset (Figures 1, 6a, Supplementary Figures S10, S11). Later, *SELENOJ2* was lost independently in Otophysi (e.g *Danio rerio)* and Tetraodontidae (e.g. *Tetraodon nigroviridis*). On the other hand, *SELENOJ* underwent a Sec-to-Cys conversion in Salmonidae.

**Figure 6.**
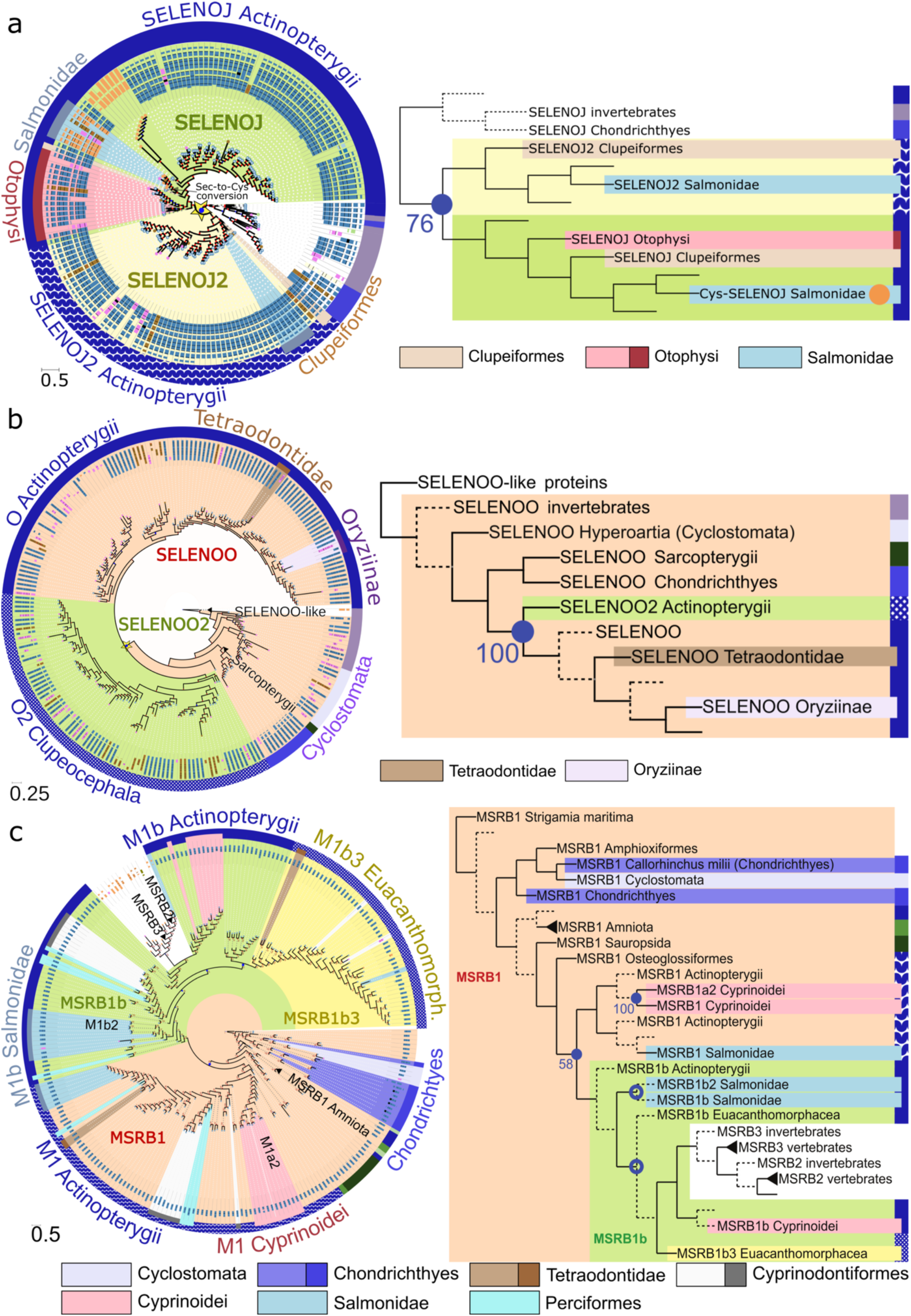
Phylogeny of the SELENOJ, SELENOO, and MSRB families. **a, b, c)** Gene tree visualizations of the SELENOJ, SELENOO, and MSRB families, respectively. See full feature description in Figure 2c. Expanded gene and species trees are available: Supplementary Figure S10, S11 (SELENOJ), S12, S13 (SELENOO), S14, S15 (MSRB).

#### SELENOO

Selenoprotein O (SELENOO) is the largest known selenoprotein by molecular weight, and belongs to a widespread protein family recently functionally characterized as a mitochondrial pseudokinase that transfers AMP to various protein substrates, implicated in oxidative stress response and metastasis [36, 37]. Previous studies identified a *SELENOO* paralog named *SELENOO2* in *D. rerio*, which was absent in the genomes of *Takifugu rubripes*, *G. aculeatus*, *O. latipes*, *and T. nigroviridis* [16]. In our analysis, we replicated these results, and yet we observed the presence of *SELENOO2* in many other fish genomes, including among others all of Cyprinodontoidei (toothcarps) and Carangaria (e.g. jacks, flatfishes), which implies a complex pattern of gene evolution (Figures 1, 6b, Supplementary Figures S12, S13). Ultimately, we hypothesize that *SELENOO2* emerged at the root of Clupeocephala but was subsequently lost in specific lineages, namely Tetraodontidae, Oryziinae, Anabantaria, and almost all Perciformes (e.g. *G. aculeatus*).

#### Methionine sulfoxide reductase B (MSRB)

Methionine sulfoxide reductase B (MSRB, previously known as SELENOR), are enzymes that reduce oxidized methionine residues, with roles for antioxidant defense and redox-based modulation of various pathways including actin assembly [38]. Humans possess one Sec-containing gene, *MSRB1*, and two Cys paralogs, *MSRB2* and *MSRB3*. Our analyses (Figure 6c, Supplementary Figures S14 and S15) indicate that the three genes predate the origin of vertebrates, since members of each subfamily were detected in invertebrates. Within fish lineages, *MSRB1* underwent a complex history of duplications and losses. We previously identified a Sec-containing gene, *MSRB1b*, in five actinopterygian species [16]. Based on the protein tree topology (Figure 6c, Supplementary Figure S14; UFB support 58%) and the pattern of gene occurrence across the species tree (Supplementary Figure S15), we mapped its origin to the subclade of Osteoglossocephalai, which comprises all Actinopterygii in our dataset except the *MSRB1b*-lacking species *Erpetoichthys calabaricus* and *Lepisosteus oculatus*. Gene counts per species further indicated secondary *MSRB1b* duplications; yet, the reconstructed protein tree exhibited a scattered topology, without the typical expected clustering pattern (Figure 6c). Based mostly on the species tree, we concluded that *MSRB1b* likely duplicated in Salmonidae yielding *MSRB1b2,* and in Euacanthomorphacea yielding *MSRB1b3;* and that both *MSRB1b* and the duplicated copy *MSRB1b3* have been later lost in multiple lineages: *MSRB1b* in Esociformes, Tetraodontidae and Anabantaria and *MSRB1b3* in Perciformes, *Cynoglossus semilaevis* and Cyprinodontiformes. Notably, this reconstruction highlights at least three convergent events of *MSRB1b3* loss in Percomorphacae, and another three loss events of *MSRB1b* in Euteleosteomorpha. Relevantly, other selenoprotein families show similar patterns of convergent losses in this lineage (Figure 1), as discussed later.

#### SELENOT

Selenoprotein T (SELENOT) is a thioredoxin-like enzyme anchored to the endoplasmic reticulum (ER) membrane. It is highly conserved throughout evolution, and it has an essential role in cellular homeostasis and redox regulation [39]. While mammals have a single *SELENOT* gene, previous studies reported three homologs in actinopterygian fishes [16], here refined and expanded. We found that most actinopterygian groups possessed two *SELENOT* genes, namely *SELENOT* (aka *SELENOT1*) and *SELENOT2*; yet, Salmonidae, Otophysi and Cyprinoidei exhibited further lineage-specific duplications. Ultimately, we mapped the origin of *SELENOT2* to a duplication of *SELENOT1* at the root of Actinopterygii (Figure 7a). *SELENOT1* duplicated again to yield *SELENOT1b* in Otophysi, i.e. the group including Cyprinoidei (e.g*. D. rerio* and carps), and Characiphysae (e.g. *Electrophorus electricus, Pygocentrus nattereri).* In Cyprinoidei, further duplications of *SELENOT2* and *SELENOT1b* gave rise to the genes *SELENOT2b* and *SELENOT1b2*, respectively (Figures 1, 7a, Supplementary Figure S16). Finally, we observed yet another *SELENOT1* duplication in Salmonidae, yielding the gene *SELENOT1a2*. Interestingly, Hyperoartia also independently expanded this family through one gene duplication, which yielded a divergent paralog that we designated *SELENOT3*. We deem all *SELENOT* duplications highly reliable due to the consistency between the species tree and the protein tree topology, and to the high UFB values in the latter (Figure 7a).

**Figure 7.**
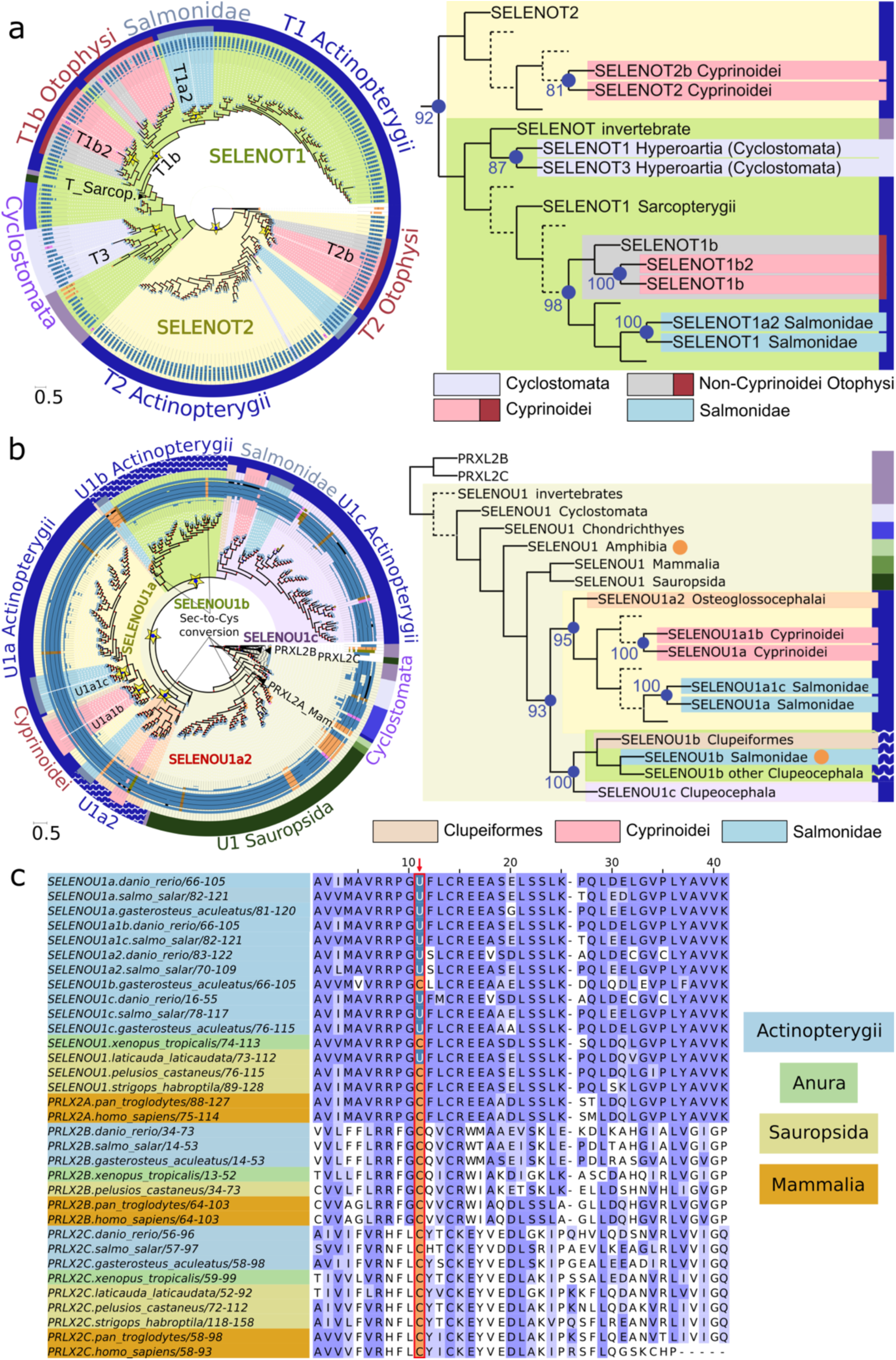
Phylogeny of the SELENOT, SELENOU families. **a, b)** Gene tree visualizations of the SELENOT and SELENOOU families, respectively. See full feature description in Figure 2c. Expanded gene trees are available as Supplementary Figures S16 (SELENOT) and S17 (SELENOU). **c)** SELENOU multiple sequence alignment. Species are color-labelled according to lineage. Sec residues are marked in blue and Cys residues in orange.

#### SELENOU

The Selenoprotein U (SELENOU) family includes selenoprotein genes in birds and fishes, the latter reported to carry up to three Sec-containing paralogs [10, 16]. Mammals and amphibians are reported to encode only Cys-homologs, with human containing three paralogs (*SELENOU1/PRXL2A*, *SELENOU2/PRXL2C*, *SELENOU3/PRXL2B*), involved in diverse functions such as antioxidant defense, prostaglandin biosynthesis, and glycolysis regulation. The SELENOU gene family exhibits a scattered pattern of occurrence, implying a complex evolutionary history involving gene duplications, losses, and Sec-to-Cys conversions. Here, we provide our best attempt to explain their history with current data (Figures 1, 7b, 7c, Supplementary Figures S1, S17).

The split of *SELENOU1/2/3* predates vertebrates, as members of the three subfamilies were detected in invertebrate species (Supplementary Figures S17). All Sec-encoding genes detected in this family belonged to the *SELENOU1* branch. Within fish, we mapped a first well-supported duplication of the *SELENOU1* gene in Osteoglossocephalai, yielding *SELENOU1a2* (Figure 7b). This gene was then lost at the root of the large group of Acanthomorphata, while it converted to Cys homolog in *Esox lucius*. Prior to both of these events, we traced two additional *SELENOU1* duplications in Clupeocephala, yielding the *SELENOU1b* and *SELENOU1c* genes. *SELENOU1b* was then lost independently in three lineages: Otophysi, Esocidae and Atherinomorphae (Supplementary Figure S1). Finally, we detected another two independent well-supported *SELENOU1* duplications in Cyprinoidei and Salmonidae, yielding the gene *SELENOU1a1b* and *SELENOU1a1c*, respectively (Figure 7b). Lastly, within tetrapods, *SELENOU1* underwent at least four convergent Sec-to-Cys conversions (Figure 7c), specifically in Theria (i.e. placentals and marsupials), Anura (frogs), Testudines (turtles), and Psittaciformes birds (parrots).

#### SELENOW, SELENOV and MIEN1

Selenoprotein W (SELENOW) is a widespread glutathione-dependent antioxidant oxidoreductase implicated in resolution of inflammation, with reported roles in muscle growth and differentiation, as well as in protecting neurons from oxidative stress during neuronal development [40, 41]. Selenoprotein V (SELENOV) is a placental-specific putative oxidoreductase expressed in testis and potentially involved in antioxidant defense and ER stress, whose C-terminus shares substantial sequence similarity to SELENOW [42]. Our present analyses are consistent with SELENOV emerging in placentals by duplication of *SELENOW* (Figure 8a). It was previously reported that the *SELENOV* gene was lost in *Gorilla gorilla* [16]. By analyzing the species tree annotated with *SELENOV* gene predictions (Supplementary Figure S1), we pinpointed additional potential losses, which we sought to corroborate by searching RNA datasets (Methods). Due to *SELENOV* detection in RNA data, we ultimately dismissed all apparent losses except two: the known one in *Gorilla gorilla*, and a novel one in Odontoceti (toothed whales, including dolphins) (Figure 1).

**Figure 8.**
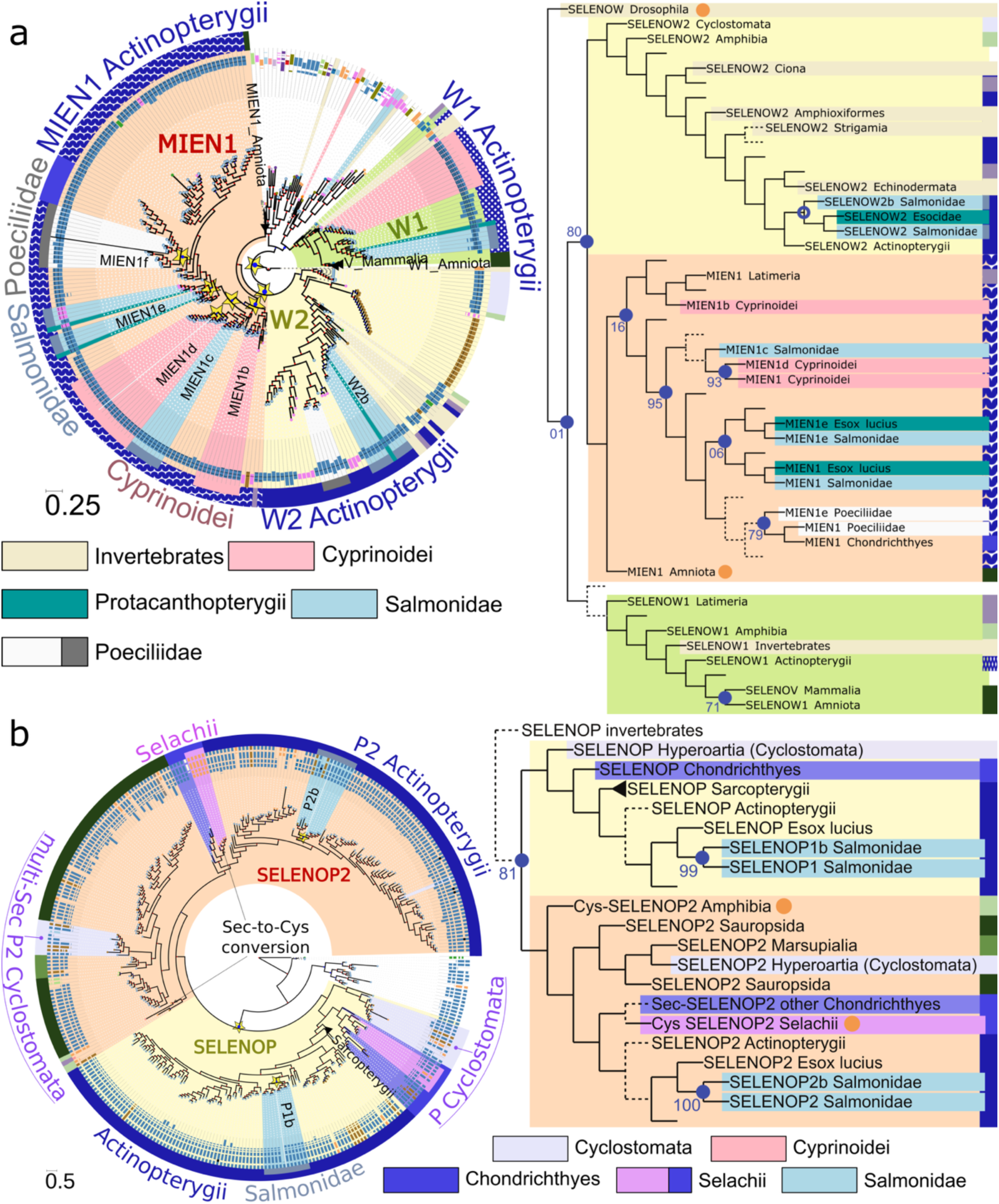
Phylogeny of the SELENOW and SELENOP families. **a, b)** Gene tree visualizations of the SELENOW and SELENOOP families, respectively. For tree annotation and legend details, see Figure 2c. Expanded gene trees are available as Supplementary Figures S18 (SELENOW) and S22 (SELENOP).

Besides *SELENOV*, mammals have a single other homolog in this gene family: *SELENOW*, also referred to as *SELENOW1*. Our analyses indicate that this gene was independently lost in Cyclostomata, Chondrichthyes and within Actinopterygii, specifically in Acanthomorphata, resulting in its absence from the majority of extant fish genomes (Supplementary Figure S1).

Earlier work pinpointed multiple *SELENOW* homologs in non-mammalian vertebrates, including two subfamilies named *SELENOW2* and *MIEN1* (formerly Rdx12) [43]. Indeed, our analysis recovered these two clades: they were defined by a duplication node supported by ∼35 species, and by performing BLASTP of each sequence, we noted that the majority of one clade matched sequences annotated as SELENOW2, while the other matched MIEN1 annotations. These clades were confidently placed as sister groups, with the *SELENOW* and *SELENOV* clade as their closest relative (Figure 8a). Within the SELENOW2/MIEN1 group, all genes belonging to invertebrates were placed in the *SELENOW2* branch (Supplementary Figure S18). This suggests that *SELENOW2* predates vertebrates, while *MIEN1* emerged within this lineage. We detected at least one *MIEN1* gene in each vertebrate genome, except Cyclostomata wherein it was consistently absent. Therefore, we propose that *MIEN1* originated by a *SELENOW2* duplication at the root of Gnathostomata. In Amniota, MIEN1 genes carry Cys in place of Sec, implying a Sec-to-Cys conversion in this clade. On the other hand, several Actinopterygian lineages contained multiple Sec-containing *MIEN1* copies, which we ascribed to secondary lineage-specific gene duplications, detailed hereafter. Because of poor consistency between the protein tree topology and species taxonomy, here we heavily relied on the species tree to infer gene histories. Cyprinoidei contained three copies; in the protein tree, *MIEN1* and *MIEN1d* exhibited the expected topology for duplication, while *MIEN1b* was placed closer to the root, possibly due to long branch attraction (Figure 8). We consider most likely that two *MIEN1* duplications at the root of Cyprinoidei generated *MIEN1b* and *MIEN1d*. Salmonidae also exhibited three copies: *MIEN1*, *MIEN1e*, *MIEN1c*. We argue that these emerged independently of those in Cyprinoidei, since virtually all other Clupeocephala contained a single copy (Supplementary Figure S1). *MIEN1e* was found in Salmonidae and also in their outgroup *E. lucius*, which together form the superorder of Protacanthopterygii. We ascribe the *MIEN1e* origin to a duplication at the root of this lineage; the protein tree topology is consistent with this scenario, albeit with very low UFB support in the corresponding node (6%). The other copy (*MIEN1c*), in contrast, was detected in Salmonidae only. Surprisingly, this clustered with Cyprinoidei *MIEN1* and *MIEN1d*. We consider that the most likely scenario is that *MIEN1c* emerged in Salmonidae after the split from *E. lucius*, then its high divergence confounded its phylogenetic position. Lastly, Poeciliidae contained two copies, *MIEN1* and *MIEN1f*. We ascribed this to a gene duplication at the root of this lineage; the protein tree topology is consistent with this, despite an unexpected clustering with *MIEN1* of Chondrichthyes.

The *SELENOW2* branch of the gene tree includes selenoprotein genes from invertebrates, Cyclostomata, Amphibia, and diverse Actinopterygii. The absence of Amniotes implies a gene loss in this lineage, as previously reported [16]. In Actinopterygii, on the other hand, the history of this gene is more obscure and difficult to solve. *SELENOW2* occurs in a scattered pattern across the species tree, apparently implying many independent losses, which may seem unlikely (Supplementary Figure S1). Therefore, we first set to assess the quality of our tree-based orthology assessment using a complementary syntenic analysis. We observed limited but detectable syntenic conservation around *SELENOW1* across Actinopterygii and Tetrapoda, shown by consistent nearby presence of *EHD2/EHD2A* (Supplementary Figure S19). Similarly, genes in the *SELENOW2* tree branch are consistently located in the vicinity of the *TRPM7* gene in all tested Actinopterygii and Amphibia (Supplementary Figure S20). As per MIEN1, syntenic conservation was clear within Actinopterygii (*SLC4A1A, UBTF, ATXN7I3/ATXN7I3A*) and within Amniota (*IKZF3*, *PGAP3, GRB7, ERBB2*), but not in-between (Supplementary Figure S21). No common nearby genes were observed between *SELENOW1*, *SELENOW2* and *MIEN1*. Altogether, these analyses corroborated the true orthology of these protein clades. Therefore, we proceeded to map *SELENOW2* losses within Actinopterygii by examining the species tree annotated with gene presence (Supplementary Figure S1) and seeking validation of putative losses in RNA data. Strikingly, we identified 13 independent losses (after dismissing two due to *SELENOW2* detection in RNA sequences; Supplementary Table T2), displayed in Figure 1. Although additional *SELENOW2*-like RNA sequences were recovered in certain genomes (e.g., *Oryzias melastigma, Ictalurus punctatus*), further inspection indicated that these sequences likely represented contamination, and they were therefore excluded.

#### SELENOP

Selenoprotein P (SELENOP) is the only mammalian selenoprotein with multiple Sec residues, e.g. 10 in human and *M. musculus*, due to its Sec-rich C-terminal tail. SELENOP is a key player in selenium transport and distribution, serving as the primary selenium carrier in the bloodstream, delivering selenium to peripheral tissues via receptor-mediated uptake. We recently characterized the evolution of the *SELENOP* gene family, showing that its Sec content can vary by two orders of magnitude across Metazoa [44]. Its C-terminal tail is challenging to identify using genomic data due to its fast evolution, so that here we mostly focused on gene-level detection. As expected, we identified two phylogenetic clades of vertebrate genes, multi-Sec *SELENOP/SELENOP1*, and single-Sec *SELENOP2/SELENOPB*. While the former is reportedly ubiquitous in vertebrate genomes, the latter was lost in Eutheria [44]. Moreover, *SELENOP2* was converted to a Cys homolog in Anura. Our present analysis (Figure 8b, Supplementary Figure S22) confirmed these findings, and offered additional insights into early vertebrate evolution.

Invertebrate *SELENOP* homologs are consistently recovered outside of the clade containing both vertebrate paralogs, supporting the hypothesis that the *SELENOP1/SELENOP2* duplication arose at the root of vertebrates. Most fish genomes contained the two Sec-*SELENOP* genes; a notable exception was the Selachii division within Chondrichthyes (i.e. sharks), where *SELENOP2* was converted to Cys homolog. In Salmonidae, we traced one lineage-specific gene duplication for each paralog, yielding *SELENOP1b* and *SELENOP2b*. Like most fish, Cyclostomata contains two genes, one within the *SELENOP1* branch and the other within *SELENOP2*. Interestingly, the latter displays a unique CXXU motif (rather than the more typical UXXC) in its N-terminal thioredoxin-like domain. Moreover, though *SELENOP2* of other vertebrates encodes a single Sec, in Cyclostomata it exhibits an extraordinary expansion of Sec content: it harbors 162 and 139 in-frame UGA codons in the lampreys *Lethenteron reissneri* and *Lampetra fluviatilis*, respectively. These genes were detected in genome assemblies, and their sequence was confirmed by highly similar matches in transcriptome shotgun assembly (TSA) data of other Cyclostomata, spanning all the putative transcript (Supplementary Document D1). Analogously to other *SELENOP* genes [44], the vast majority of putative Sec residues are encoded within an expanded modular C-terminal region, and two SECIS elements are detected downstream. We argue that all these in-frame UGAs are translated as Sec: (i) homologous SECIS elements from human, *M. musculus* and *D. rerio SELENOP* have been shown to support processive insertion at multiple Sec-UGA sites in a variety of *in vitro* and *in vivo* models [4, 44]; and (ii) the lamprey coding sequence contains more than 130 stop codons exclusively of the UGA type (rather than a mixture with UAA or UAG), providing strong evidence of selective pressure to preserve the same peculiar gene architecture that is observed in hundreds of multi-Sec SELENOP genes across Metazoa [44]. With this record, lampreys surpasses all other metazoans in the number of Sec-UGAs in *SELENOP*, including the previous high of 132 reported in bivalves [44]. Between the SECISes, we noted the presence of a distinctive region rich in CA dinucleotides (∼1500-1800nt) of unknown significance (Supplementary Document D1).

#### Other Salmonidae-specific duplications: SELENOE, SELENOF, SELENOH

Our analysis revealed a substantial selenoproteome expansion specific to the lineage of Salmonidae, which may suggest an increased functional reliance of this clade on selenium. Besides the aforementioned expansions in the GPX, DIO, TXNRD, MSRB, SELENOT, SELENOU and SELENOW protein families (Figure 1), we detected well-supported gene duplications of *SELENOE* and *SELENOF*, yielding *SELENOEb* and *SELENOFb*, respectively, and another two for *SELENOH*, yielding *SELENOHb* and *SELENOHc* (Figure 9, Supplementary Figures S23, S24, S25). Briefly, SELENOF is a ER-resident selenoprotein involved in redox quality control [45], recently pinpointed as tumor suppressor [46, 47]. Our results show that *SELENOF* is ubiquitous to vertebrates. On the other hand, SELENOE (formerly Fep15), a functionally uncharacterized selenoprotein distantly related to SELENOF [12], was found as selenoprotein only in fishes: Anura carry a Cys-homolog, while the rest of tetrapods (Amniota) lost the gene altogether. Notably, we mapped the origin of the *SELENOE* gene to the root of Gnathostomata, since we did not find it in Cyclostomata nor in the non-vertebrate outgroups. Finally, SELENOH is a widespread nucleolar oxidoreductase that regulates redox homeostasis and suppresses DNA damage through the p53 pathway, and it is implicated in cancer proliferation [48–50]. Apart from the Salmonidae-specific duplications, no other gene evolution event was detected for *SELENOE*, *SELENOF*, and *SELENOH*, with the only exception of the loss of *SELENOH* in Cyclostomata, and in independently in *Callorhinchus milii*.

**Figure 9.**
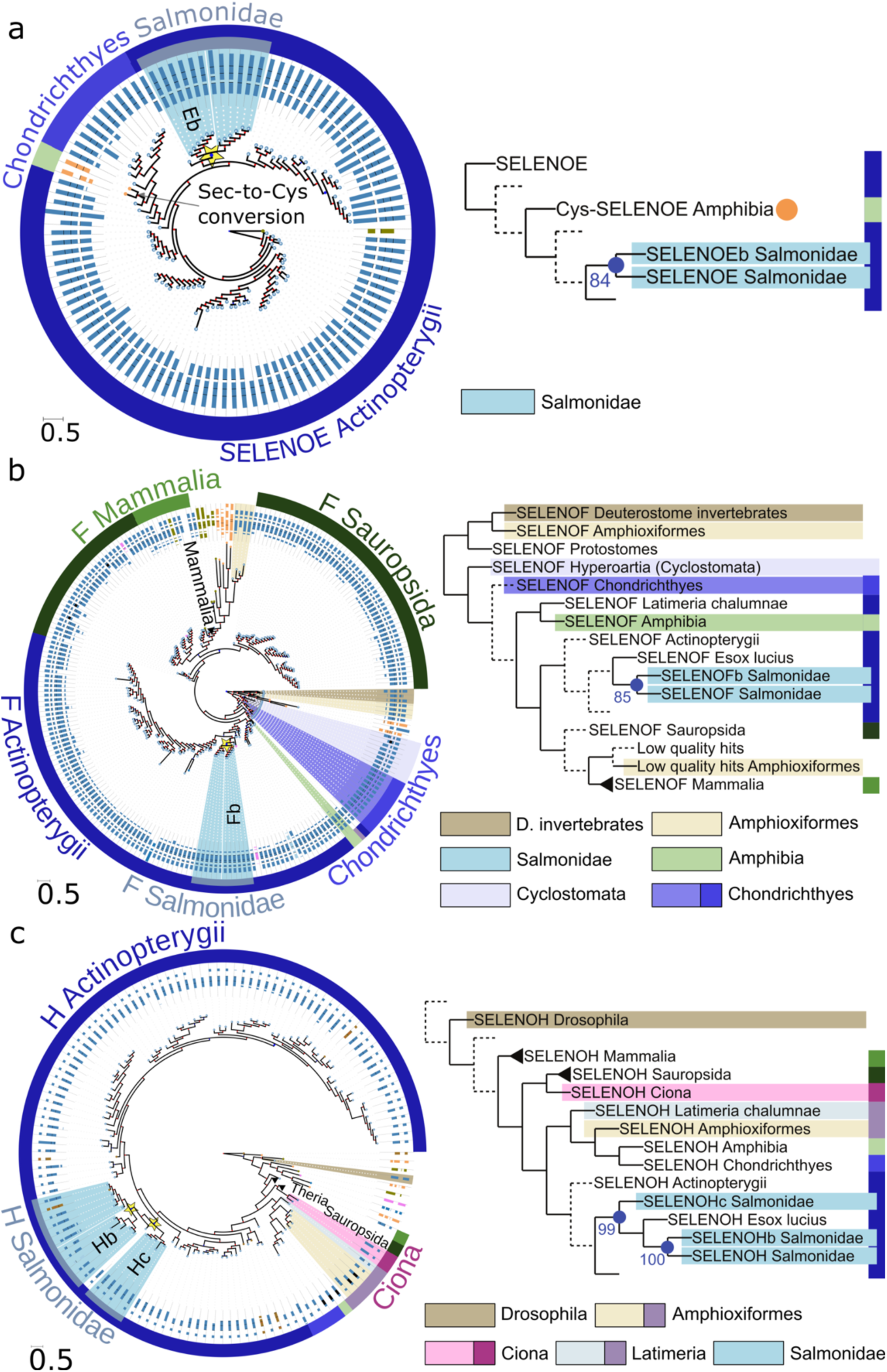
Phylogeny of SELENOE, SELENOF, and SELENOH. **a, b, c)** Gene tree visualizations of the SELENOE, SELENOF, and SELENOH families, respectively. For tree annotation and legend details, see Figure 2c. Expanded gene trees are available as Supplementary Figure S23 (SELENOE), S24 (SELENOF), and S25 (SELENOH).

It is worth mentioning that, because all the Salmonidae genomes in our dataset belong to its major subgroup, Salmoninae, we had initially parsimoniously mapped all the gene duplications described above to Salmoninae. However, inspection of intron positions relative to protein sequence alignments (Figures 2, 3, 4, 6) revealed that they were conserved across these lineage-specific duplications. This pattern is consistent with DNA-based gene duplication rather than RNA-based retrotransposition. Consequently, the whole-genome duplication (WGD) previously reported at the root of Salmonidae [51] provides the most plausible explanation for the origin of these genes, leading us to map these events to Salmonidae as a whole rather than exclusively to Salmoninae.

#### Other families: SEPHS2, SELENOL, SELENOK, SELENOM, SELENON, SELENOS

Hereafter, we briefly summarize the remaining vertebrate selenoproteome families, wherein our present analysis recapitulated earlier results without novel findings. As already introduced, SEPHS2 (aka SPS2) is a selenoprotein whose function is critical for Sec biosynthesis. As expected [16, 22], we identified the *SEPHS2* gene in all vertebrate lineages. (As discussed in the first section of the Results, the gene appears to be absent from many assemblies of birds, but we attribute this to a sequencing bias rather than true gene loss). The *SEPHS2* gene is intronless only in genomes of Eutheria (placentals). As previously shown [16], the ancestral intron-containing *SEPHS2* gene (referred to as *SEPHS2a*) duplicated by retrotransposition at the root of Mammalia, yielding an intron-less copy referred to as *SEPHS2b*. Both copies are present in extant genomes of Marsupialia, while *SEPHS2a* was then lost in Eutheria. Our results confirmed these findings (Supplementary Figures S1, S26). Thus, the sole placental *SEPHS2* gene is actually *SEPHS2b*. Of note, *SEPHS2* duplicated at the root of vertebrates, and independently also in various other metazoan lineages, to yield a non-Sec/non-Cys paralog known as *SEPHS1/SPS1* [22]. The vertebrate *SEPHS1* gene carries threonine in place of Sec. This kind of conversion is highly unusual for selenoproteins, since Sec is typically catalytic and it is highly dissimilar from canonical amino acids other than Cys. Indeed, SEPHS1 appears to lack the enzymatic activity of SEPHS2 (selenophosphate synthesis from selenide), and instead has a function in gene expression-mediated control of redox homeostasis [6, 52].

SELENOL is a peculiar selenoprotein featuring two nearby Sec residues that can form a diselenide bond. It features a peroxiredoxin-like motive suggestive of a redox-related function [13]. As expected [16], we detected the *SELENOL* gene in nearly all vertebrates except Tetrapoda, confirming a gene loss at the root of this lineage (Supplementary Figure S1).

The remaining selenoprotein families were detected in a single Sec-containing copy in all investigated genomes (with the only exception of *SELENOM*, lost in Hyperoartia; Supplementary Figure S1), barring pseudogenes and spurious absence attributed to imperfect assembly quality. Curiously, all these single-copy selenoproteins localize to the ER. They include: SELENOK, involved in calcium homeostasis and resolution of ER stress [53]; SELENOM and SELENOS, involved in unfolded protein responses [54]; and SELENON, involved in calcium and redox homeostasis [55].

#### Shifts in selenoproteome-to-proteome ratio across vertebrate evolution

Our analyses revealed that the vertebrate selenoproteome expanded through many gene duplications in fish lineages. The conservation of intron structures and the phylogenetic correspondence with documented WGD suggests that many of these gene duplications resulted from such events (Figure 1). This raises the question of whether such selenoproteome expansions are adaptive, or are neutral consequences of having a larger proteome. To gain insights in this matter, we compared the size of our predicted selenoproteome with that of the annotated proteome for each species (Figure 10). We fit a zero-intercept linear model separately for each phylogenetic clade of interest (Mammalia, Sauropsida, Chondrichthyes, Salmonidae, Cyprinoidei, and non-Salmonidae non-Cyprinoidei Actinopterigyii), with the hypothesis that the slope (i.e. the selenoproteome-to-proteome ratio) may differ among clades. We found that Actinopterygii have significantly larger selenoproteome-to-proteome ratios than Mammalia (Figure 10). Importantly, aquatic environments have long been hypothesized to favor selenoprotein usage [56], and our present analysis mostly supports this pattern. Yet, some lineages diverge from this trend: aquatic Salmonidae and Cyprinoidei, which have the biggest selenoproteomes across vertebrates, show low ratios (albeit with substantial statistical uncertainty, due to the few available genomes and their spread in apparent proteome size; see their wide confidence intervals in Figure 10). On the other hand, Sauropsida exhibits relatively high ratios despite being land-dwelling. Overall, this data is best explained by an ancestral state of rich selenoproteome-to-proteome ratio, subsequently strongly decreased in Mammalia, and possibly slightly increased in Actinopterygii.

**Figure 10.**
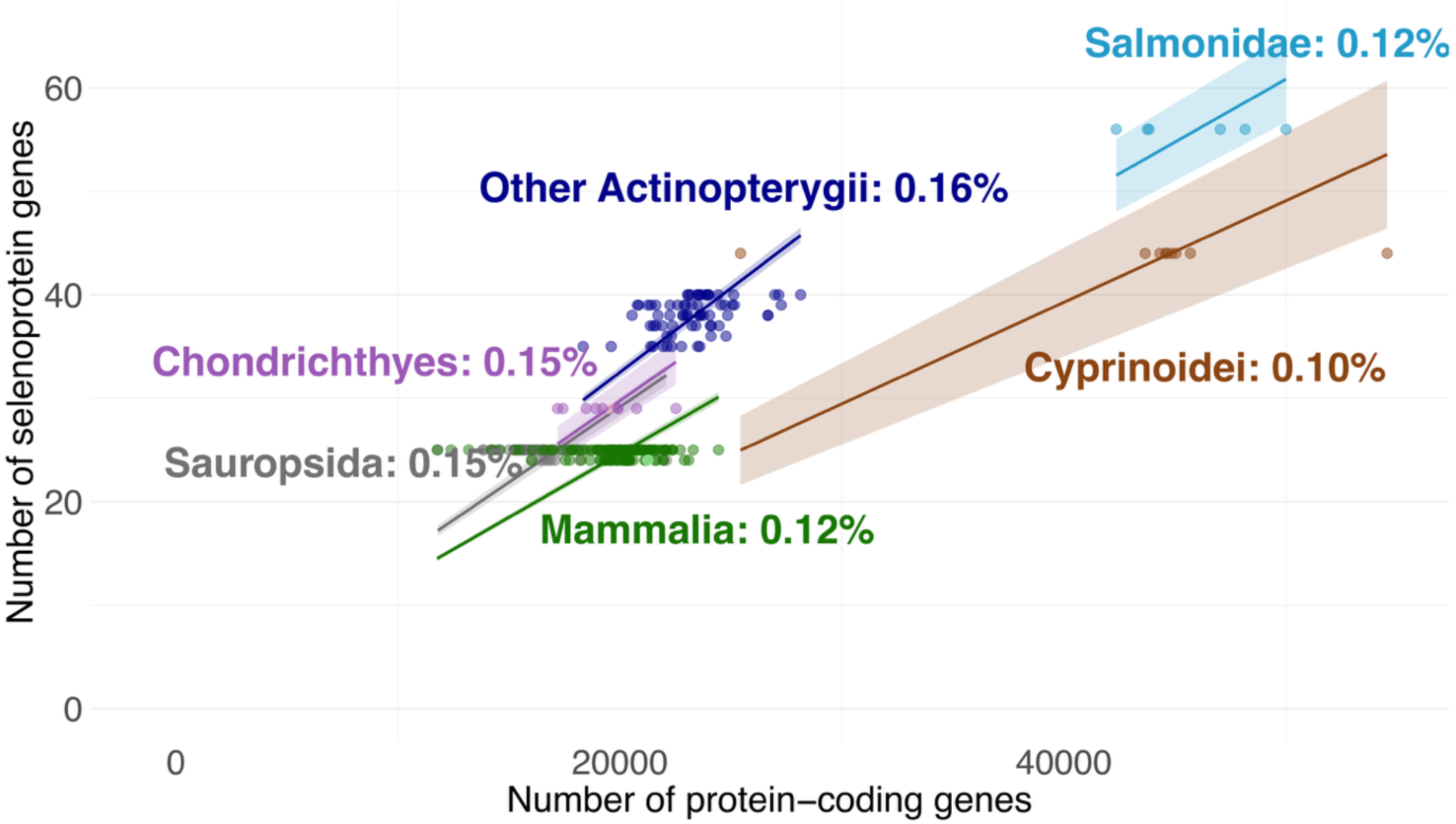
Genomic correlation between annotated proteome size and the number of predicted selenoproteins. Each point represents a species and is color-labelled according to lineage. Trend lines corresponding to zero-intercept linear models fit separately for each lineage (Methods) are displayed. Shading indicates the 95% confidence interval on the slope. The colored text provides the color key and includes the average selenoproteome-to-proteome ratio as percentage.

To formally assess the impact of habitat on selenoproteome size, we fitted a series of regression models relating proteome size (denoted as P) to selenoproteome size (S), with habitat (H: aquatic vs. terrestrial) included either as an additive term or as an interaction with proteome size. Because species are not statistically independent, we implemented phylogenetic generalized least squares under both Brownian motion (neutrality) and Ornstein–Uhlenbeck (balancing selection) models, thus incorporating the species tree as a covariance structure. We also tested alternative codings of habitat in which, besides fish, Cetacea and Amphibia were classified as aquatic. Additionally, we considered models in which only Cetacea were classified as aquatic, while Amphibia were treated as terrestrial. Across all tested models, the effect of aquatic habitat was highly significant (Table 1), consistently indicating an enrichment of the selenoproteome in aquatic lineages. Models with the interaction term (S ∼ P + H*P + 0) provided a better fit than additive models (S ∼ P + H + 0), consistent with aquatic species having a proportionally larger selenoproteome relative to proteome size. Likelihood ratio tests further supported the habitat effect, as each model outperformed its corresponding null model lacking H (Table 1).

**Table 1.** Analysis of selenoproteome-to-proteome ratio in aquatic vs terrestrial vertebrates. The table contains results of different regression models fitted with data from 263 vertebrates (Methods). Two formulas were evaluated: an additive model (S ∼ P + H + 0) and an interaction model (S ∼ P + H*P + 0*), where S is the number of predicted selenoproteins (Selenoprofiles), P the number of annotated proteins (Ensembl), and H a binary variable for aquatic vs. terrestrial habitat. The “+0” denotes that the intercept was constrained to zero. Models included simple linear regression (LM) and phylogenetic generalized least squares (PGLS) under either Brownian motion (BM, neutrality) or Ornstein–Uhlenbeck (OU, balancing selection). All fish were considered aquatic; in some models, Cetacea and/or Amphibia were also classified as aquatic (see the third column). The fourth column reports the P-value of the habitat coefficient, and the fifth column the likelihood ratio test P-value comparing each model to its corresponding null model without H (S ∼ P + 0).

| Formula | Model | Tetrapods aquatic? | H coefficient<br>P-value | LRT p-value |
| --- | --- | --- | --- | --- |
| $S \sim P + H * P + 0$ | PGLS + BM | No | 2.2e-16 | 2.51e-77 |
| $S \sim P + H * P + 0$ | PGLS + BM | Cetacea + Amphibia | 2.11e-12 | 9.08e-68 |
| $S \sim P + H * P + 0$ | PGLS + BM | Cetacea only | 2.2e-16 | 1.45e-72 |
| $S \sim P + H * P + 0$ | PGLS + OU | No | 2.2e-16 | 2.51e-77 |
| $S \sim P + H * P + 0$ | PGLS + OU | Cetacea + Amphibia | 2.11e-12 | 9.08e-68 |
| $S \sim P + H * P + 0$ | PGLS + OU | Cetacea only | 2.2e-16 | 1.45e-72 |
| $S \sim P + H * P + 0$ | LM | No | 2.2e-16 | 3.38e-184 |
| $S \sim P + H * P + 0$ | LM | Cetacea + Amphibia | 2.2e-16 | 7.44e-88 |
| $S \sim P + H * P + 0$ | LM | Cetacea only | 2.2e-16 | 9.11e-142 |
| $S \sim P + H + 0$ | PGLS + BM | No | 8.24e-08 | 9.8e-69 |
| $S \sim P + H + 0$ | PGLS + BM | Cetacea + Amphibia | 8.69e-07 | 2.45e-63 |
| S ~ P + H + 0 | PGLS + BM | Cetacea only | 6.15e-07 | 1.24e-63 |
| S ~ P + H + 0 | PGLS + OU | No | 8.24e-08 | 9.8e-69 |
| S ~ P + H + 0 | PGLS + OU | Cetacea + Amphibia | 8.69e-07 | 2.45e-63 |
| S ~ P + H + 0 | PGLS + OU | Cetacea only | 6.15e-07 | 1.24e-63 |
| S ~ P + H + 0 | LM | No | 2.2e-16 | 2.19e-134 |
| S ~ P + H + 0 | LM | Cetacea + Amphibia | 2.2e-16 | 1.26e-84 |
| S ~ P + H + 0 | LM | Cetacea only | 2.2e-16 | 5.02e-106 |

## DISCUSSION

The recent acceleration of genome sequencing and the rise of global biodiversity initiatives has generated an unprecedented abundance of genomic data across vertebrate lineages. This surge presents a unique opportunity to revisit fundamental questions about gene evolution, functional diversity, and lineage-specific adaptations. Here, we leveraged this genomic wealth to provide an updated view of the composition and evolution of the vertebrate selenoproteome, obtained through analyses of hundreds of genomes and meticulous phylogenetic reconstruction which greatly expanded upon previous endeavors. We reinforced and refined existing findings, and uncovered new genes and evolutionary events.

Our reconstruction of the ancestral vertebrate selenoproteome consists of 27 selenoprotein genes representing all the 19 known vertebrate Sec-encoding gene families (Figure 1). Our conclusions mostly overlap with our earlier analysis [16], with the exceptions of *GPX2* and *SELENOE* (both now mapped to Gnathostomata rather than Vertebrata), and *GPX3b* (Vertebrata rather than Actinopterygii). Besides, we now consider *TXNRD3* as the ancestral gene in the *TXNRD1/TXNRD3* group, with *TXNRD1* emerging in Gnathostomata. We also concluded that these selenoproteins emerged specifically at the root of Vertebrata: *GPX3b, DIO2, DIO3, SELENOI, SELENOP2*. Among these, *SELENOI* stands out in that the whole family appeared at the root of vertebrates: there is no homolog that includes the Sec position, regardless of the aligned amino acid, outside of vertebrates.

On the other hand, our present work substantially expands the catalog of selenoproteins in fish. Aquatic environments have long been hypothesized to favor selenoprotein usage, a trend first suggested by Lobanov et al. from limited genomic data [56]. Subsequent analyses further corroborated this model by showing relaxed purifying selection on selenoprotein genes in terrestrial vertebrates [57], alongside reduced Sec content in SELENOP, an established proxy for selenium demand [44, 57]. Our present analysis of

the selenoprotein-to-proteome ratio, leveraging the largest dataset to date and phylogeny-aware models, provides further support for this evolutionary pattern. However, we note that the difference between the selenoproteome of aquatic and land-dwelling species appears primarily driven by mammalian loss and actinopterygian expansion.

The underlying cause of the selective advantage of Sec in aquatic environments is arguably yet unclear. It has been proposed that land-dwelling organisms must face (i) reduced selenium bioavailability and (ii) increased oxygen exposure [56, 57]. While the latter seems intuitively relevant, it contradicts one of the prevalent proposed biochemical rationale for usage of selenium (Sec) over sulphur (Cys) [58, 59]: its resistance to (over)oxidation - a trait that would suggest selenoproteins should be more beneficial in terrestrial environments. Moreover, our data shows that selenoproteins belonging to antioxidant families are more prevalent in fish compared to tetrapods. While further research is required to fully resolve this question, we argue that the current evidence better supports selenium bioavailability as the most likely evolutionary driver of Sec usage in aquatic vs land environments.

These observations may depict our own selenoproteins as an unstable remnant of an aquatic past. Yet, it is worth noting that (i) land-living tetrapods exhibit substantial purifying selection, preventing Sec to Cys conversions in most genes [60]; and (ii) the aquatic selenoproteome is far from static, having expanded under putative adaptive forces at the root of Actinopterygii and in multiple of its clades. We found that these duplications occurred primarily in protein families linked to antioxidant defense, either with well-established roles (GPX, MSRB) or with putative/indirect functions (SELENOT, SELENOU, SELENOW). In contrast, no variation was observed in ER-resident proteins with functions in calcium homeostasis (SELENOK, SELENON) or protein quality control / unfolded protein response (SELENOM, SELENOS). While this trend was not absolute - e.g. ER-resident SELENOE and SELENOF duplicated in Salmonidae, and the frequently duplicated SELENOT is also ER-resident - it suggests that selenoproteome expansions in Actinopterygii may have been driven primarily by selection for enhanced antioxidant capacity. While we did not investigate the exact mechanism of gene duplications, we note that, strikingly, 40 selenoprotein duplications out of the 56 detected across vertebrates were mapped to taxonomic nodes corresponding with reported WGD events (Figure 1). This observation reinforces the importance of WGDs in shaping vertebrate proteomes.

Among tetrapods, our most striking finding is the recurrence of convergent evolutionary events, including Sec-to-Cys conversions in SELENOU1 and GPX6, as well as independent losses of SELENOV, all of which are supported by robust phylogenetic evidence. It is important to note that, because our reconstruction is based on maximum parsimony, our estimate of such events represents a conservative lower bound. Interestingly, both SELENOV and GPX6 are the “youngest” mammalian selenoproteins and they are specifically expressed in the testis, a tissue recognized for its role in the evolution of new genes [61].

Our study has some limitations. First, we restricted our search to genes homologous to known selenoprotein families, because existing methods for identifying novel selenoproteins suffer from low specificity [62]. As a result, we may have overlooked lineage-specific selenoproteins, particularly in fish lineages where selenoproteome expansions are evident. Second, we could only detect events within lineages represented in our dataset. Given the dynamic nature of the Actinopterygii selenoproteome, additional genome sequences from this group will likely reveal further duplications. Third, our analysis assumes the species tree as ground truth, which may not be entirely accurate. For instance, the convergent loss of the same genes in Tetraodontidae, Perciformes, Anabantaria, and *Cynoglossus semilaevis* (Figure 1) could be explained as a single ancestral loss if the non-Perciform Eupercaria and Carangaria were instead the outgroups of these clades - yet, we found no supporting evidence for this alternative phylogeny in the literature [63]. Fourth, further data or alternative methodological approaches could refine the evolutionary history of selenoprotein families with the most complex gene dynamics, such as SELENOW and SELENOU, and provide deeper insight into lineages with highly complex genome evolution, such as Cyprininae.

Despite these limitations, we anticipate that our characterization of vertebrate selenoproteins will largely hold up over time and provide a meaningful framework for future evolutionary and functional research.

## MATERIAL AND METHODS

### Genome sequences and species tree

We downloaded all genome assemblies and annotations available in Ensembl version v.108 [64] comprising 315 genomes, each corresponding to a different species or strain. These included 310 vertebrates, two tunicates, one nematode (*C. elegans*), one insect (*D. melanogaster*), and yeast *Saccharomyces cerevisae* (here excluded from all analyses). The phylogenetic tree of species was downloaded from Ensembl Compara v.e110 [20]. To refine our reconstruction of the ancestral vertebrate selenoproteome, we also analyzed NCBI assemblies from species with informative taxonomy; identifiers are provided in Supplementary Table T3. Specifically, we included various early-branching vertebrates: nine cartilaginous fish (Chondrichthyes) and nine jawless vertebrates (Hyperoartia class, belonging to the Cyclostomata superclass); then four amphioxiformes (Branchiostoma), serving as additional close relatives to vertebrates, besides the two tunicates already in Ensembl; and two remote outgroups: *Strongylocentrotus purpuratus* (Echinodermata, deuterostome) and *Strigamia maritima* (Arthropoda, protostome). These 24 genomes, together with the 314 Ensembl assemblies, constituted our main dataset. Later, we searched two additional Cyclostomata genomes, belonging to the class of Myxini, whose predictions were included in the protein tree reconstruction of TXNRD. For the rest of families, we distinguished between Cyclostomata, Myxini, and Hyperoartia specific events by inspecting the species tree annotated with selenoprotein predictions (Supplementary Figure S1). For follow up analyses to confirm or dismiss gene evolution events, we then inspected gene predictions on an extended set of 1,172 vertebrate genome assemblies downloaded from NCBI, using the NCBI taxonomy tree as the rough backbone for evolutionary interpretation.

### Selenoprotein gene prediction

Selenoprotein gene prediction was performed using Selenoprofiles [19, 65, 66] version v4.5.0, available at https://github.com/marco-mariotti/selenoprofiles. Selenoprofiles is a computational pipeline to identify members of the known selenoprotein families and related proteins. It uses several homology-based gene prediction programs, and employs a set of manually curated multiple sequence alignments of selenoprotein families to scan genomes for homologues. The profiles we used are available at https://github.com/marco-mariotti/selenoprotein_profiles (v1.3.0). We scanned genome assemblies for all known selenoprotein families in Metazoa. For most analyses, we focused on the predictions labelled as “selenocysteine”, i.e., the genes encoding for selenoproteins, with no apparent pseudogene features (e.g. frameshifts).

### Phylogenetic reconstruction of selenoprotein families

The protein sequences of all Selenoprofiles predictions across Ensembl genomes were collected using the Selenoprofiles *join* utility, and alignments were computed using ClustalO v.1.2.4 [67]. These were used as input to phylogenetic reconstruction using IQ-TREE v3.0.1 [68]. For each selenoprotein family, the best-fitting substitution model was automatically selected using ModelFinderPlus. The selected models are listed in Supplementary Table T4. Phylogenetic trees were inferred under the corresponding substitution model, and the branch support values were estimated using 1,000 ultrafast bootstrap replicates [68]. We visualized gene trees with the show_tree program in the biotree_tools package v0.0.6 (https://github.com/marco-mariotti/biotree_tools), based on the ETE3 python module [69], specifically adapted for custom visualization in this manuscript. The resulting vectorial images were further annotated manually using the Inkscape software. Sequence and tree manipulations throughout the project were carried out using the programs *alignment_tools* and *tree_tools* from the biotree_tools package, all based on ETE3 [69]. Sequences in the *SELENOP* gene family were trimmed prior to tree building to remove the highly divergent C-terminal Sec-rich tail, using custom code based on Pyaln v0.1.4 (https://github.com/marco-mariotti/pyaln).

### Analysis of gene evolution events

We manually inspected gene and species trees to determine gene evolution events: losses, duplications, and Sec-to-Cys conversions. To detect potential duplications, gene trees were processed using the species overlap algorithm: the number of common species between the left and right branch of every ancestral node was used as a quantitative indicator of a gene duplication in the last common ancestor of those species. Putative duplication nodes were colored in the gene tree visualization, and their size was made proportional to this indicator, facilitating our manual review.

A visualization of the species tree together with all gene predictions in each genome was produced using the *tree drawer* utility available in Selenoprofiles, customized as follows. First, each gene prediction corresponding to a multi-member gene family were assigned to a mammalian-centric homology group, hereafter referred to as gene “subfamily” (e.g. *GPX1, GPX2, GPX3*, etc), using the *orthology* utility of Selenoprofiles. Then, we encoded the expected selenoproteome for every lineage based on literature; e.g. bats, since they are mammals with no further lineage-specific events reported, were expected to carry one Sec-encoding gene for each of *GPX1, GPX2, GPX3, GPX4* and *GPX6*. In our custom *tree drawer* visualization (Supplementary Figure S1), we marked each gene prediction based on the deviation from such expectations; e.g., bats were labelled as missing a Sec-*GPX6*. This highlighted the presence of extra genes (putative duplications) and missing genes (putative losses). Ultimately, the joint analysis of gene and species trees led us to manually map each event into the species tree, following criteria of maximum parsimony and consistency with the known phylogenetic relationship among species.

Lastly, each event was critically evaluated to exclude potential artifacts. We required support by at least two sister species for most events. Single-species events were deemed reliable only if supporting independent data (e.g. RNA sequences) existed. Due to imperfect quality of genome assemblies, we sought confirmation of each individual gene loss using independent sequence data from the lineage of interest, as follows. At this stage, each putative gene loss consisted in the lack of an apparently functional gene (i.e., without mutations introducing frameshifts or in-frame stops) in the set of Selenoprofiles predictions on genome assemblies in our main dataset. First, we inspected the full set of Selenoprofiles predictions for the relevant gene family in an extended set of NCBI genomes from the relevant clade. If the loss was not dismissed, we then selected the protein sequence of the putatively lost gene from a closely related organism, and searched it using TBLASTN at the NCBI web server [70] in all RNA data available for the clade that included the focal species, i.e. TSAs, Expressed Sequence Tags (ESTs), and Reference RNA sequences (refseq_rna). While absence of transcriptomic data alone is not conclusive evidence of gene loss (e.g. due to tissue-specific expression), we argue that it provides strong support when it is combined with the concomitant absence from multiple genome assemblies in the same clade.

### Grx-domain analysis in TXNRD genes

To investigate the distribution of Grx domains encoded in TXNRD genes, we first collected a set of representative TXNRD protein sequences and scanned them for conserved domains using InterProScan [71]. We extracted the sequences containing the Grx domain and thus manually curated a high-confidence alignment of Grx domains of TXNRDs. These were used as sequence profiles to search for homologous matches in Ensembl vertebrate genomes using Selenoprofiles. Next, we cross-analyzed this data with those TXNRD gene predictions in the same genomes classified as either *TXNRD1* or *TXNRD3* by the Selenoprofiles *lineage* utility. We used a custom script based on pyranges v1 [72] to assign Grx domain predictions to *TXNRD1* and *TXNRD3* genes, if located within their extended genomic coordinates (±5,000 nucleotides). Unassigned Grx domains were discarded. Finally, we visualized domain architecture and evolutionary patterns by visualizing data through custom scripts based on the ETE3 library [63].

### Synteny analysis

Through custom scripts based on the library pyranges v1 [72], we selected all gene annotations that overlapped the genomic loci of the gene of interest (*TXNRD1*, *TXNRD3*, and *SELENOW2/MIEN1*; Supplementary Figures S7, S8, S19, S20) predicted by Selenoprofiles, extended on each side by 300,000 bp, in a set of representative species. For these syntenic analyses, we used Ensembl v.115 annotations, upon realizing their annotated gene names were improved compared to v.108, which we had used for the rest of our work. We then used python code based on the graphics library pyrangeyes v.1.0.7 (https://github.com/pyranges/pyrangeyes) to produce custom visualizations wherein genes with the same name were colored accordingly, allowing to assess their syntenic conservation. To inspect the homologous region in species lacking the gene of interest, we performed an analogous process on a gene that consistently occurred in the its neighborhood in other species (*CHST11* for *TXNRD1*; *TRPM* for *SELENOW2*; *RSPH4A* for *SELENOW1*). Synteny representations were manually collated next to the species tree using the Inkscape software.

### Gene nomenclature

We strived to assign gene names as consistent as possible with existing resources and literature. To do so, we consulted the relevant publications on vertebrate selenoprotein evolution and nomenclature [73], as well as current annotations at Uniprot, Ensembl, and NCBI. The reference recommendation for gene names is to derive them from the corresponding mammalian orthologs [73]. The genes emerged from non-mammalian duplications —many of them previously unreported— were named by appending alphabetical or numerical suffixes to the source gene, e.g. *GPX4b*, *SELENOW2*. To enhance readability, suffixes are written in lowercase throughout this work. To avoid ambiguity, when creating new gene names we systematically skipped the suffixes “a” and “1” (in favor of “b” or “2”), because these are inconsistently used in the literature to refer to non-mammalian orthologs of single-copy mammalian genes; e.g. *SELENOW* is sometimes referred to as *SELENOW1* in species that also possess the paralog *SELENOW2*.

### Regression models of selenoproteome and proteome sizes

Regression models were implemented in R, using libraries *stats* for linear models and likelihood ratio tests, and *phylolm* for phylogenetic generalized least squares. A first analysis with zero-intercept linear models (Figure 10) was run with 314 annotated species (Ensembl v.108), used to estimate proteome size (number of unique protein-coding genes), whereas Selenoprofiles was used to estimate selenoproteome size. A second analysis (Table 1) was run with the species subset that had a fully resolved taxonomic tree available in Ensembl Compara (263 genomes), using either linear models or phylogenetic generalized least squares; refer to the text and Table 1 for further details.

## Supporting information

All supplementary files

## DECLARATIONS

### Ethics approval and consent to participate

Not applicable

### Consent for publication

Not applicable

### Availability of data and materials

The latest Selenoprofiles code is available at https://github.com/marco-mariotti/selenoprofiles4 and documented at https://selenoprofiles4.readthedocs.io/.

Selenoprotein sequences used in this study split by family can be found in Supplementary Data D2.

### Authors’ contributions

MT analysed all the data, performed the evolutionary analyses, prepared all the figures and wrote the draft of the manuscript. JL-F contributed to data interpretation and revised the manuscript. MM supervised the project, secured funding, provided expert interpretation of the results and was a major contributor in writing the manuscript.

## Acknowledgements

Not applicable.

## SUPPLEMENTARY DATA

**Figure S1. Selenoproteome evolution across vertebrate species.** Each row represents a species and each column corresponds to a selenoprotein family. Rectangles represent individual Selenoprofiles predictions, color-coded according according to their Selenoprofiles labels, i.e. the codon found aligned to the Sec position: blue for Sec-UGA,orange for Cys, pink for unaligned). Selected lineages denoted with a colored background are labelled on the left.

**Figure S2. Presence of genes within the SEPHS2 syntenic block across Sauropsidan genome assemblies.** The species tree of Sauropsida (plus platypus, used as outgroup) is displayed on the left. Three columns display the presence of genes SEPSH2 (blue, third column) and the genes consistently located as its neighbors in tetrapods, G6PD (first column) and ikbkg (second column). The third column also shows the presence of SEPHS1 (purple), located in a different chromosomic location, here serving as positive control. Note the scattered lack of G6PD, ikbkg, and SEPHS2 from many bird genomes, attributed to imperfect quality of genome assemblies, as discussed in the text.

**Figure S3. Gene tree of the Glutathione peroxidase (GPX) family.** Subfamilies are color-labeled as follows: GPX4 in orange, GPX4b in yellow, GPX3b in purple, GPX3 in green, GPX6 in pink, GPX5 in light purple, GPX2 in beige, GPX1 in brown, and GPX1b in blue. Refer to Figures 2 and 3 for details.

**Figure S4. Gene tree of the Iodothyronine deiodinase (DIO) family.** Subfamilies are color-labeled as follows: DIO1 in orange, DIO2 in yellow, DIO3 in green and DIO3b in purple. See also Figures 2c (layout explanation) and 4 (compressed tree).

**Figure S5. Iodothyronine deiodinase (DIO) evolution across Salmonidae species.** Each row represents a species and each column corresponds to a selenoprotein family. Nodes represent individual Selenoprofiles predictions and are color-coded according to the codon aligned at the Sec position: blue for Sec-UGA, orange for Cys.

**Figure S6. Gene tree of the Thioredoxin reductase (TXNRD) family.** Subfamilies are color-labeled as follows: TXNRD1 in green, TXNRD2 in orange, TXNRD3 in yellow. See also Figures 2c (layout explanation) and 5 (compressed tree).

**Figure S7. Analysis of synteny of TXNRD1 across vertebrates.** The figure displays the genomic neighborhoods of TXNRD1 and/or CHST11 (see Methods) in representative genomes. Same-colored gene structures share the same annotated gene name. The species tree is shown on the left. Species highlighted in grey do not contain the TXNRD1 gene.

**Figure S8. Analysis of synteny of TXNRD3 across vertebrates.** The figure displays the genomic neighborhoods of TXNRD3 and/or CHST13 (see Methods) in representative genomes. Same-colored gene structures share the same annotated gene name. The species tree is shown on the left. Species highlighted in grey do not contain the TXNRD1 gene.

**Figure S9. Vertebrate species tree representing the presence/absence of Grx domain in TXNRD1/TXNRD3 sequences.** Colors are assigned according to the TXNRD gene tree.

**Figure S10. Gene tree of the Selenoprotein J (SELENOJ) family.** Subfamilies are color-labeled as follows: SELENOJ in green and SELENOJ2 in yellow. See also Figures 2c (layout explanation) and 6 (compressed tree).

**Figure S11. Selenoprotein J (SELENOJ) evolution across Actinopterygii species.** Each row represents a species and each column corresponds to a selenoprotein family. Nodes represent individual Selenoprofiles predictions and are color-coded according to their amino acid content at the UGA position: blue for Sec-containing sequences, orange for Cys-containing sequences, and pink for other classifications inferred by Selenoprofiles.

**Figure S12. Gene tree of the Selenoprotein O (SELENOO) family.** Subfamilies are color-labeled as follows: SELENOO in green and SELENOO2 in orange. Nodes highlighted in brown and light purple indicate sequences from Tetraodontidae and Oryziinae species, respectively. See also Figures 2c (layout explanation) and 6 (compressed tree).

**Figure S13. Selenoprotein O (SELENOO) evolution across Actinopterygii species.** Each row represents a species and each column corresponds to a selenoprotein family. Nodes represent individual Selenoprofiles predictions and are color-coded according to their amino acid content at the UGA position: blue for Sec-containing sequences, orange for Cys-containing sequences, and pink for other classifications inferred by Selenoprofiles.

**Figure S14. Gene tree of the Methionine sulfoxide reductase B (MSRB) family.** See also Figures 2c (layout explanation) and 6 (compressed tree).

**Figure S15. Methionine sulfoxide reductase B (MSRB) evolution across Actinopterygii species.** Each row represents a species and each column corresponds to a selenoprotein family. Nodes represent individual Selenoprofiles predictions and are color-coded according to their amino acid content at the UGA position: blue for Sec-containing sequences, orange for Cys-containing sequences, and pink for other classifications inferred by Selenoprofiles.

**Figure S16. Gene tree of the Selenoprotein T (SELENOT) family.** Subfamilies are color-labeled as follows: SELENOT1 in green and SELENOT2 in yellow. Nodes highlighted in light blue indicate sequences from Salmonidae species, while lightpink and grey indicate sequences from Cyprinoidei and Otophysi species, respectively. See also Figures 2c (layout explanation) and 7 (compressed tree).

**Figure S17. Gene tree of the Selenoprotein U (SELENOU) family.** Subfamilies are color-labeled as follows: SELENOU1a in yellow, SELENOU1a2 in orange, SELENOU1b in green, SELENO1c in purple and Cysteine-homolog PRLX2A in beige. Nodes highlighted in light blue indicate sequences from Salmonidae species, while lightpink indicate sequences from Cyprinoidei. See also Figures 2c (layout explanation) and 7 (compressed tree).

**Figure S18. Gene tree of the Selenoprotein V/W (SELENOV/W) family.** Subfamilies are color-labeled as follows: SELENOW1 from Actinopterygii species in green, SELENOW1 from tetrapoda species in beige, SELENOV in purple, SELENOW2a in yellow and SELENOW2c in orange. Nodes highlighted in light blue indicate sequences from Salmonidae species, while light pink and light grey indicate sequences from Cyprinoidei and Cyprinodontoidei species, respectively. See also Figures 2c (layout explanation) and 8 (compressed tree).

**Figure S19. Analysis of synteny of SELENOW1 across vertebrates.** The figure displays the genomic neighborhoods of SELENOW1 and/or rsph4a (see Methods) in representative genomes. Same-colored gene structures share the same annotated gene name. The species tree is shown on the left. Species highlighted in grey do not contain SELENOW1 gene.

**Figure S20. Analysis of synteny of SELENOW2 across vertebrates.** The figure displays the genomic neighborhoods of SELENOW2 and/or TRPM7 (see Methods) in representative genomes. Same-colored gene structures share the same annotated gene name. The species tree is shown on the left. Species highlighted in grey do not contain SELENOW2 gene.

**Figure S21. Analysis of synteny of MIEN1 across vertebrates.** The figure displays the genomic neighborhoods of MIEN1 (see Methods) in representative genomes. Same-colored gene structures share the same annotated gene name. The species tree is shown on the left.

**Figure S22. Gene tree of the Selenoprotein P (SELENOP) family.** Nodes highlighted in light blue indicate sequences from Salmonidae species. See also Figures 2c (layout explanation) and 8 (compressed tree).

**Figure S23. Gene tree of the Selenoprotein E (SELENOE) family.** See also Figures 2c (layout explanation) and 9 (compressed tree).

**Figure S24. Gene tree of the Selenoprotein F (SELENOF) family.** Nodes highlighted in light blue indicate sequences from Salmonidae species See also Figures 2c (layout explanation) and 9 (compressed tree).

**Figure S25. Gene tree of the Selenoprotein H (SELENOH) family.** Nodes highlighted in light blue indicate sequences from Salmonidae species. See also Figures 2c (layout explanation) and 9 (compressed tree).

**Figure S26. Gene tree of the SEPHS family.** See also Figure 2c (layout explanation).

**Table S1. Comparison of Glutathione Peroxidase (GPX) gene nomenclature between Xue et al. and this study.** The table lists the gene names assigned to GPX family members in Xue et al. and the corresponding nomenclature used in this study (Ticó et al.). The third column provides relevant notes on differences in naming, such as updated classification, subfamily assignment, or gene duplication interpretations.

**Table T2. Detection of SELENOW1, SELENOW2 and MIEN1 transcripts across vertebrate species.** Boolean values denote presence (True) or absence (False) of RNA-derived sequences assigned to each family based on similarity scoring against the curated reference dataset extracted from the SELENOW gene tree. Corresponding taxonomic lineage is provided for each species.

**Table T3. List of NCBI species and their corresponding genome assembly accession IDs used in this study.**

**Table T4. Substitution models selected by IQ-TREE for phylogenetic tree reconstruction in each selenoprotein family.**

**Document D1. Sequence analysis of SELENOP2 in lampreys**. The document contains the sequences of SELENOP2 genes in two Cyclostomata, obtained via Selenoprofiles in genome data, then corroborated by RNA sequences. SECIS elements were predicted using SECISearch3.

**Document D2. Protein sequences of all selenoproteins predicted and analyzed in this study.** For details on file structure and content, please refer to the accompanying README file.

