## Supplementary material for "Tracing the vertebrate selenoproteome evolution reveals expansions in ray-finned fishes and convergent depletions in tetrapods": All supplementary files: Supplementary document D1.lamprey_SELENOP_analysis.docx

Legend: CDS (3’region) SECIS	CA-rich region

Underlined= RNA evidence: homologous matches (>80% identity, 0.0 Evalue) found via blastn in TSAs of other cyclostomata species.

The list of such IDs is provided at the end of this document

>Lethenteron_reissneri_SELENOP_protein 162 UGAs

MGPQPWGKPPVALLALLAGAASTLAARPRHVAVPAGGGGGGRGGDEVGRVDEAKDALALCRPAPSWRDADGVDPMAEHAGCVRVLALLAGGCRLURSQATELDKLRARLDARGLSDVAYALLSERGPHSRSSARRLARRLLPSSSSPSSSSSTSTSSAVTAHAQASSGPDLWTLLGGQRNDIFVYDRCGRLTVRLSLPYSFLDFPYVESAIVATHAKEVCGACPPSGEATATATVATADVAKVRPGDGGGVGGGEEAERPAIVDYDCKDDDDDKRTEAHGEEDAETRGKIDALRRQLKKSPPNHHGHHGHHHGDGGAGSGSRUPLPPGRVRFGYRUAMERAGVGGGDGGDGGDGDGVAURUHURTLGPAAKALPGAAAAAVVATVGURCRGATRLPASCAURGGAGDVGGDVTEETUQURPRATVADURRTGKRQQSURUEAAAAAGRSSVUQUEAAGGCREEUKAATKDURULTSEUKEGVKEUKULTAEUKEGVKEUKULTSEUQEGVKEUKULTSEUKEGVKEUKULSSEUQEGVKGUKUFTSEUQEGVKEUKCLTSEUREGVKEUKULTSEUKEGVKEUKULTADUQEGVKEUKULTSEUQVGVKEUKULTSEUQEGVKEUKULTSEUKEGVKEUKULSSEUQEGVKGUKUFTSEUQEGVKEUKULTSEUREGVKEUKULTSEUKEGVKEUKULTADUQEGVKEUKULTSEUQVGVKEUKULTSEUQEGVKEUKULTSEUKEGVKEUKULSSEUKEGVKEUKULMSEUKEGVKEUKULSSEUQEGVKEUKULSSEUQEGVKEUKULTSEUQDGVKEUKULSSEUQEGVKEUKULTSEUQEGVKEUKULTSEUQEGVKEUKULSSEUKEGVKEUKULTSEUQEGVKEUKEGTSEUKEGVKEUKULTSEUQEGVKEUKQGTSEUKEGVKEUKULTAEUQEAVKEUKGTGEUQDGVKEUKULSSEUQEGVKEUKULSSEUQEGVKEUKULTSEUQEGVSEUKULTAEUQEGVKEUKULSSEUQEGVKEUKULNSEUKEGVKEUKULTSEUQEGVKEUKULTSEUQEGVKEUKULTSEUQEGVKEUKULTAEUQEGVKEUKULSSEUQEGVKEUKULTAEUKAEVKEUKULTAEUKAGVKEUKULTSEUQEGVKEUKUMTSEUKRGGVAAEAGRGGGSPLTSDPAN

>Lethenteron_reissneri_SELENOP_nt CDS + putative 3’ UTR

ATGGGGCCTCAGCCGTGGGGCAAACCCCCGGGGCGACCGCCACCGCCACCGTCATCACGGCCACCACGGCCGGTGGCGCTGACGGTGGCGGTGGCGTTCCTGGCCCTGTTGGCCGGAGCCGCGTCCACGCTGGCGGCTAGACCTCGGCACGTCGCAGTCCCGGCCGGGGGAGGAGGAGGAGGAGGGGACGAGGTGGGGAGGGTGGACGAGGCGAAGGACGCCCTCGCGCTCTGCCGACCGGCGCCCTCCTGGAGGGACGCGGACGGCGTCGACCCGATGGCGGAGCACGCGGGATGCGTGCGCGTGCTGGCGCTGCTGGCCGGAGGCTGCCGCCTCTGACGATCGCAGGCCACGGAGCTGGACAAGCTGCGCGCACGCCTGGACGCGCGCGGCTTGAGCGACGTGGCCTACGCGCTGCTCAGCGAGCGCGGCCCCCACTCGCGCTCCTCCGCCCGCCGCCTGGCACGACGCCTCCTCCCCTCCTCCTCCTCCCCATCATCCTCCTCCTCGTCGTCCGCGGTGACGGCCCACGCTCAGGCCTCGTCCGGCCCCGACCTGTGGACCTTGTTGGGCGGGCAGCGCAACGACATCTTCGTCTACGACAGGTGCGGACGCCTAACGGTGCGCCTTTCGCTGCCCTACTCCTTCCTGGACTTCCCGTACGTGGAGTCGGCCATCGTGGCCACCCACGCCAAGGAGGTGTGCGGGGCCTGCCCGCCCAGCGGCGAGGCCACCGCCACCGCCACCGTCGCCACCGCCGACGTCGCGAAGGTTCGCCCTGGCGACGGCGGCGGCGTCGGAGGTGGGGAGGAGGCGGAGCGGCCGGCGATCGTGGACTACGACTGCAAAGACGACGACGACGACAAGAGAACCGAGGCGCACGGAGAAGAAGACGTGGAGACGCGCGGCAAGATCGACGCGCTGCGACGGCAGCTCAAGAAGTCGCCTCCAAACCACCACGGCCACCACGGCCACCACCACGGGGACGGCGGCGCCGGCAGCGGTAGCCGGTGACCGCTGCCGCCCGGTCGCGTTCGTTTCGGCTACCGGTGAGCGATGGAGCACGCGGGCGTCGGTGGCGGTGACGGTGGTGACGGTGGTGACGGTGACGGCGTCGCGTGACGGTGACATTGACGGACGCTCGGCCCTGCGGCCAAAGCGTTGCCGGGCGCCGCCGCCGCCGCGGTCGTGGCGACCGTCGGCTGACGGTGCCGCGGGGCGACGCGGCTGCCGGCGTCGTGCGCCTGACGGGGCGGCGCCGGTGACGTCGGCGGTGACGTCACGGAGGAGACGTGACAGTGACGGCCCCGGGCGACGGTCGCCGACTGACGCCGGACGGGCAAGCGGCAGCAGTTGTGACGCTGAGAGGCGGCGGCTGCCGCCGGGCGGAGCTCCGTGTGACGGTGAGAGGCGGCGGGGGGCTGTCGGGAGGAATGAAAGGTGGCGACGAAGGACTGACGGTGACTGACGAGCGAATGAAAGGAGGGAGTCAAGGAATGAAAGTGACTCACCGCTGAATGAAAGGAAGGAGTGAAGGAATGAAAGTGACTGACGAGTGAATGACAGGAAGGAGTGAAGGAATGAAAGTGACTCACCAGCGAATGATGAAAGGAAGGAGTGAAGGAATGAAAATGACTCAGCAGTGAATGACAGGAGGGAGTCAAGGGATGAAAGTGATTCACGAGTGAATGACAGGAAGGAGTGAAGGAATGAAAGTGACTGACAAGTGAATGACGGGAAGGAGTGAAGGAATGAAAATGACTGACGAGTGAATGAAAGGAGGGAGTGAAGGAATGAAAGTGACTCACCGCTGATTGACAGGAAGGAGTGAAGGAATGAAAGTGACTCACTAGTGAATGACAGGTAGGAGTGAAGGAATGAAAGTGACTGACGAGTGAATGACAGGAAGGAGTGAAGGAATGAAAGTGACTCACCAGCGAATGAAAGGAAGGAGTGAAGGAATGAAAATGACTCAGCAGTGAATGAAAGGAAGGAGTGAAGGAATGAAAGTGACTGATGAGTGAATGAAAGGAAGGAGTGAAGGAATGAAAGTGACTCAGCAGTGAATGACAGGAAGGAGTGAAGGAATGAAAATGACTCAGCAGTGAATGACAGGAGGGAGTGAAGGAATGAAAGTGACTCACGAGTGAATGACAGGATGGAGTGAAGGAATGAAAATGACTCAGCAGTGAATGACAGGAGGGAGTGAAGGAATGAAAGTGACTCACTAGTGAATGACAGGAAGGAGTGAAGGAATGAAAGTGACTCACGAGTGAATGACAGGAAGGAGTGAAGGAATGAAAGTGACTCAGCAGTGAATGAAAGGAAGGAGTGAAGGAATGAAAGTGACTCACGAGTGAATGACAGGAAGGAGTGAAGGAATGAAAGGAGGGGACGAGTGAATGAAAGGAGGGAGTGAAGGAATGAAAGTGATTAACTAGTGAATGACAGGAAGGAGTGAAGGAATGAAAGCAGGGGACGAGTGAATGAAAGGAAGGAGTGAAGGAATGAAAGTGACTCACCGCTGAATGACAGGAAGCAGTGAAGGAATGAAAGGGGACGGGTGAATGACAGGATGGAGTGAAGGAATGAAAGTGACTCAGCAGTGAATGACAGGAAGGAGTGAAGGAATGAAAGTGACTCAGCAGTGAATGACAGGAAGGAGTGAAGGAATGAAAGTGACTGACGAGTGAATGACAGGAAGGAGTGAGTGAATGAAAGTGACTCACCGCTGAATGACAGGAAGGAGTGAAGGAATGAAAGTGACTCAGCAGTGAATGACAGGAGGGAGTGAAGGAATGAAAGTGACTCAACAGTGAATGAAAGGAAGGAGTGAAGGAATGAAAGTGACTCACGAGTGAATGACAGGAGGGAGTGAAAGAATGAAAGTGACTGACTAGTGAATGACAGGAAGGAGTGAAGGAATGAAAGTGACTGACGAGTGAATGACAGGAAGGAGTGAAGGAATGAAAGTGACTCACTGCTGAATGACAGGAAGGAGTGAAGGAATGAAAGTGACTCAGCAGTGAATGACAGGAAGGAGTGAAGGAATGAAAGTGACTGACCGCTGAATGAAAGGCAGAAGTGAAGGAATGAAAGTGACTGACCGCTGAATGAAAGGCAGGAGTGAAGGAATGAAAGTGACTGACGAGTGAATGACAGGAAGGAGTGAAGGAATGAAAGTGAATGACGAGTGAATGAAAGCGAGGAGGGGTCGCGGCTGAGGCCGGGAGAGGTGGGGGCTCGCCTCTGACCTCTGACCCCGCCAACTAAGCCGCCCCCCTCCTCTGAACCTCGCGAGATGTTGACTCCCTCGCCTGGCACGCAGGAGGTTGTGAGTTCGACCCCGACCCCTCACGACGGACGCCTGCTGGATGTGTGGGAGTTTGCGGGTCTCTCTCTCTCTCCCCACGTGGTCGCTTGGATTCTTCTACCGGGGTCTCCGCGCTTAGAGAGTATTTGTCGGCTGTAAATTCGTCGGCGGGCTCCCATCGCTCAAATTCGTAGCGACCCGCACGTGGGCCGTCGATTCGTGCCGTTGAAGGGGGAAGGCACCGGACGAATCCACGCGTCCCACGGGCAGAGCGATCGATGGGGGGGGGGAGAGAGAGAGAGGGAGCCACGCAGACCCCCAAATATACAGCGGACGACGTCAGGAGGTGGGGTCGGTCGTGATCGAACCCACGGAGACCTCCTGCGCACCCGGCGAGGTAGTTAACATCTCGAGCAAATGAACGGATGATGGGTGGGAACGGAACGGATGACGGTCGGACGAGTGGACGGACGACTCGGAAGTTTGGGTGAAGTTTTGGGAATTTTTTTATTATTTTGCTGCCCCCCCATTTGGTTACTCGGGGTTAGCCCCCCCCCACGTTCATCCCTACCCCCCCGCGGGGGTCGCGCACATGAAGGGGGGAGCAGAAACTACGCCGTAGGGGCTCTCGTCGGAACGGGGGGCGGTCACGGCGGGGGTTGGTCTGTTGGGGGGGGGGGCGCCTTGGGTTATGGGCAAGTGGTCGAGTCGGACCACGCACACTCACACGTACACACACATGTACACACACACGCATACACACACACTCACACGTACACACACATGTACACACACACGCACACTTACACATACACTCACACGTACACACACGCACACACCTACACATACACACACACTCACACGTACACACACGCACATGCACACACATACGCACACACACGCACACACACATGTACACACACATGCACACTTACACATACAAGCACACATGCACACATACACACACACACTTACACATACACACACACACGTACACACACATACACACATACGCACGCACTTAAACACACACTTACATATACACACACTCACAAGCACATGTACACACACACACGCACACACTTACACATACACACACGCACGTACACACACGCACACACACATGTACACACACACATACACACACGCACACACTTACACATACACACACACTTACACATACACACACACTCACACGCATGTACACACACGCACTCACACACGCGTACACACGCACGTACACACCCACGCACACATACACACTTACACATACACACACACGCGCACACACGCACGCCCATACACACACGCACACGCGCACACACACACGCACACATACACACACGCACATTTACACATACACGCACGTGCGCACACACACGCACGTACACACACTTACACAGGCACACGCACGCACACAGGCGCACGCACGCACACACACGCACGCACATACACGCACGCACATACACACATGCACATACGCACATGTACACGCGCATACAACGGACACGCACATACAGTATATGACCGAAGGTCGATGTTTTTTTCATTTCTTAATGTTGATCTTGTTTCCTCGTTGTAGTTCGTGCTGTCGGATGTGAACCGTTACTCGTGTTGTGAGAGGCTGTCGTGGGGATTGTGTACACACACACACACGCGTACACACACACGCACACACATACACACGCACACACACACACACGCGCACACAAAGACACACGCACACACACACGCGTACACACACACGCACACACACACACACGCGTACACACACACGCACACACATACACACGCACACACACACACGCACACACAAAGACACACGTACACACACACACACGCGTACACACACACGCACACACACACACGCGTACACACACACGCACACACATACACACGCACACACACACGCGCACACAAAGACACACGCACACACACACACGCACACACAAAGACACACGTACACACACACACACGCGTACACACACACGCACACACACACACGCGTACACACACACGCACACACATACACACGCACACACACACGCGCACACAAAGACACACGTACACACACACGTACACACACACGCACACACACGCGCGCGTCTCATCCGCACGCACACGCGTGCAGACGCGTCACGCACACAGACACACAAACCTGTACGCACACACAAACTTGCCAGTGTCACGCGTACACACACACACAAACCCGCACGTGTCACACACACATGGACACACACAAACCCACGTGTCACACACACATGGACACACACAAACCCACGTGTCACACACACATGTACACACACATGTACACACACACACGTGTAGACACACACGCGTCACACTCGCGTCACACACACATGTACACACGCACGCGTAGACACACACACACAAACCCGCACGCGTCACACACACATGTACACACACACACATGTACACACACACACGCGTACACACACACACGTACACACACACGTGTACACACACACACGCACATGACCGCTCCCCACGACATCCCAGCGGACGACACGTACAACTTGTTCTTAATTCTCTCTCTAGTCCATGAAGTGGTTTAAACATTTAAAAACAAAATAAAAAGATCGTGGGCGATAGCGACGGCTTTCTTCTCGCCCCCCCCCCCCCATCATGGGGGTGCGGAGCGGGGGAGGGGTCACACCAAAATCGTGCGCGAGCCACAGCGGCGAGGAAATGTTAACTCCCTCGCCTGGCGTGCAGGAGGTCCGTGGCTTAGACGCTCGATTCCTGACCCCCCCCCCCCCGCTCCCCCACGAGACGTTGACCGCCCGCTTCCAGCCTGGGTTCGATTCCCCGGTGTGTGGCGGGCGGGCAAAGGGGGGGGGGGGGGGGGTGGCAGGCGAGAGGAAACCGGTGAAAGGTAGCGCCGACACACGTCACGATCACGCGTGCCGAGGATGAACCTCCGAGCGGTCGGACGGGACGCGACGGACGGGACGCGACGGATCGACTGAACTTGACGGATTCGACCGTTCGAGTCGCGGACGCGGCGGCACCGGCGACCTCCCCACGTGCATGAAGGGCTCCCGGGAAAAGCGCGACCACGTTACCGGGACTCCGGATGGCGAGCGAGCGAGTCGCGATGGGCTCGCCGTGCCGCCGCGTCCGCCGAGGTTGCGGCGGTGGTGGTGGTGGTGGGAGGGCGGCGATGAAGAGACGCGTGTTAGAGGCTCCCACACCGCTGTGTTGTTGTGTGTGCGGGTGGTCGTTCGGACTGTGTCGGGGGGCCTCAATAAAATCTTGAGACGCTGAGCGTTTGGTAAGCGCGTGTTTATTTCACGGCGCGCTGGCGTTGTTTGTTGGTGTTGTTGGTTGTGGTTTTTGTTTGTTTGTTGTTGTTGTGTGTCCCTGTGCTACGCTGCGCGACGAACCTGGCGAGCCGGGTTCGATTCCTGCTCACGGCCAACATCGGTGACGATTATCCGAACGGGTCCTGGGCCTCCCCTAGGTCGATGCAGCCTCAAATGAATACCTGGGGCGGTGCGCGAACATCGCAACTTGGTATAGCCATGGAAAGCTTCTCGAACTACGTGGTCAACGATGATGTTAAGAAATGTGGGAGCCAGAGGACATCCCTGTCGCACCCCAAGGTGTACAGGGAATGGCTCCGACTCCTGGCCACGTAGCTCTAACCCGCTCTGCCGTGCCCACCATCACGAAGGCCCTTTAATGATTGCGAGCAGAGAGTCAGGAACTCCGAAATCACGAAGAATTGCCCACAATTGTGGCCTGTTGATGGTATCATAGGCCTTGACCAGGTCGATAAAGCAGAGGTGTAGGTCTTGTTGGAATTCCCTACACCTCTGCTGGAGCAGATGGACAGTGAACGTGGCTTCCGTGCAGGACCGCCCTGGTCTGAAACCGCATTGGGGCTCGGCAAGCCTCTCCTCAGTGTGCCTGGCAAGGCGATGTTGAATAATCCTTGCCAGGACCTTTCCCGGCACAAGACAAGACACACACATATATACAAACATTTCACTCAAGCTCGCCGTTAATCAGTGGGCCAACGTAGGCAGTGAAAATGAATCGGGGTTCAAGTTCATTTATTAGGTGTAGCCCTGTGAGCCATTCGGCTCTTGAATCGGGTGCAACCGCAACAAACAATAAACAAAAACATGAACAAGAAAACATCTTAAAGTGACTATGTGACTAAGTGCAAACAGAAAATTCAAGAAAAGACAGCAAAATATTGCAGAAAAAAGCACCGTACACCAACGCCGTGTGCAAAAATCCCAGTGGGATGCAAGTCAAACA

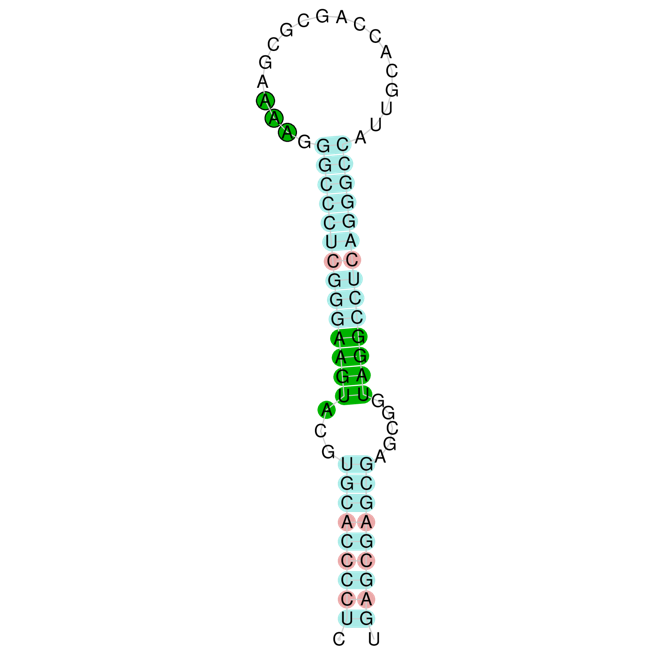

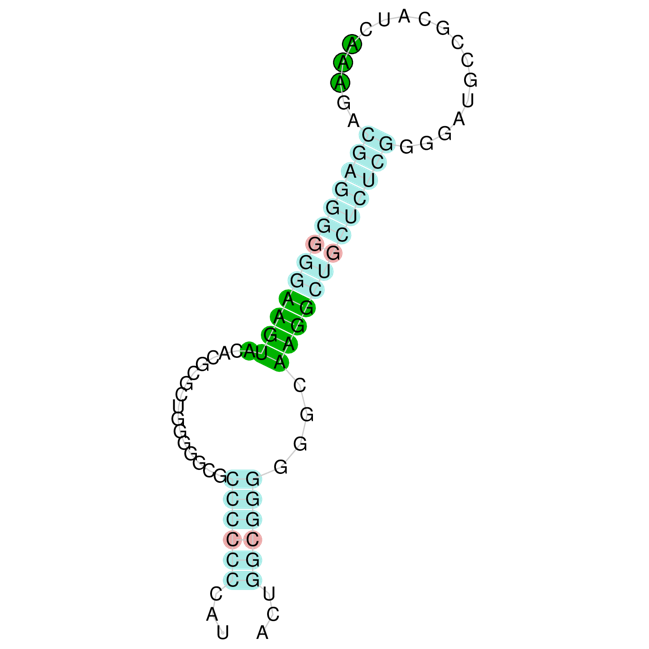

>Lampetra_fluvialis_SELENOP_protein 135 UGAs

MGPQPWGKPPGRPPPPPSSPPPRAVTVTVAVAFLALLAGAASTLAARPRHAAAAALASVPAGGGGGGGGDEVGRVDEAKDTLALCRPAPLWRDADGVDPMAEHAGCVRVLALLAGGCRLURSQATELDRLRARLDARGLSDVAYALLSERGPHSRSSARRLARRLPTPSSTSSSSSSSAVTVHAQASSGPDLWTFLGGQRNDIFVYDRCGRLTVRLSLPYSFLDFPYVESAIVATHAKEVCGACPPSGEAPATATVATADVAKVRPDGIVGGGGEEEERPAIVDYDCRDDDDDDDKRTEAHGEEDAETRGKINALRRQLKKSPPNHHGHHGHQHHGHHHGDGGAGSGSRUPLPPGRVRFGYRUAMERAGGGGGDGDGGDGGDGDGDGVAURUHURTLGPAAKATPAAVVVATVGURCRGATRLPASCAURGGDVGGDVTGDVTEGEETUQURPRATVADURRTGKRQQSURUEVAAAVAAGRSSVURUAAAGGCREEUKAATKDURULTSEUKEGVKGUKULTAEUTAGTSGUKULTAEUQGGVKEUKULSSEUKVGVKEUKULTSEUQEGVKEUKULSSEUKEGVKEUKULSSEUQEGVKGUKUFTSEUQEGLKEUKULTSEUREGVKEUKULTSEUKEGVSEUKULTAEUQEGVKEUKULTSDUKDGVKEUKULTSEUKEGVKEUKULSSQUQEGVKEUKULSSEUQEGVKEUKULSSEUKEGVKEUKULSSEUQEGVKEUKULTSEUQEGVKEUKEGTSEUKAGVKEUKULSSEUKEGVKEUKULSSEUQEGVKEUKULSSEUQEGVKEUKULTSEUKDGVKEUKULTSEUQEGVKEUKULSSEUQEGVKEUKULTSEUQEGVKEUKEGTSEUKEGVKEUKULTSEUQEGMKECKULTAEUQEGVKEUKAGTSEUKEGVKEUKULSSEUQEGVKEUKULTSEUQEGVKEUKEGTSEUKEGVKEUKULTSEUQEGVKEUKEGTSEUKEGVNEUKULTAEUQEGVKEUKEGTSEUKEGVKDUKULTSEUKEGVKEUKULSSEUKAGVNEUKULSSEUKEGVKEUKULTSEUQAGVTEUKUMTSEUERGGVAAEAGRGGGSPLTSDPAN

>Lampetra_fluvialis_SELENOP_nt CDS + putative 3’ UTR

ATGGGGCCTCAGCCGTGGGGCAAACCCCCGGGGCGACCGCCACCGCCACCGTCATCACCGCCACCACGGGCGGTGACGGTGACGGTGGCGGTCGCGTTCCTGGCCCTGTTGGCCGGAGCCGCGTCCACGCTGGCGGCTAGACCTCGGCACGCCGCAGCCGCGGCGCTGGCGTCTGTCCCGGCCGGGGGAGGAGGAGGAGGAGGAGGGGACGAGGTGGGGAGGGTGGACGAGGCGAAGGACACCCTCGCGCTCTGCCGACCGGCGCCCCTCTGGAGGGACGCGGACGGCGTCGACCCGATGGCTGAGCACGCGGGCTGCGTGCGCGTGCTGGCGCTGCTGGCCGGGGGCTGCCGCCTCTGACGATCGCAGGCCACGGAGCTGGACAGGCTGCGCGCACGCCTGGACGCGCGCGGCTTGAGCGACGTGGCCTACGCGCTGCTCAGCGAGCGCGGCCCTCACTCGCGCTCCTCCGCCCGCCGCCTGGCACGACGCCTCCCCACCCCATCCTCCACCTCCTCCTCCTCGTCGTCGTCCGCAGTGACGGTCCACGCGCAGGCCTCGTCCGGCCCCGACCTGTGGACCTTCTTGGGCGGGCAGCGCAACGACATCTTCGTCTACGACAGGTGCGGTCGCCTAACGGTGCGCCTTTCGCTGCCCTACTCCTTCCTGGACTTCCCCTACGTGGAGTCGGCGATCGTGGCCACCCACGCCAAGGAGGTGTGCGGGGCCTGCCCGCCCAGCGGCGAGGCCCCCGCCACCGCCACCGTCGCCACCGCCGACGTCGCGAAGGTTCGCCCTGACGGCATCGTCGGAGGAGGTGGGGAGGAGGAGGAGCGGCCGGCGATCGTGGACTACGACTGCAGAGACGACGACGACGACGACGACAAGAGAACCGAGGCGCACGGAGAAGAAGACGCGGAGACACGCGGCAAGATCAACGCGCTGCGACGGCAGCTCAAGAAGTCGCCTCCAAACCACCACGGCCACCACGGCCACCAGCACCACGGCCACCACCACGGGGACGGCGGCGCCGGCAGCGGTAGCCGGTGACCGCTGCCGCCCGGTCGCGTTCGTTTCGGCTACCGGTGAGCGATGGAGCGCGCGGGCGGCGGTGGCGGTGACGGTGACGGCGGTGACGGCGGTGACGGTGACGGTGACGGCGTCGCGTGACGGTGACACTGACGGACGCTCGGCCCTGCGGCCAAAGCGACGCCGGCCGCGGTCGTCGTCGCGACCGTCGGCTGACGGTGCCGTGGGGCGACGCGGCTGCCGGCGTCGTGCGCCTGACGGGGCGGTGACGTCGGCGGCGACGTCACCGGTGACGTCACGGAGGGGGAGGAGACGTGACAGTGACGGCCCCGGGCGACGGTCGCTGACTGACGCCGGACGGGCAAGCGGCAGCAGTCGTGACGCTGAGAGGTGGCGGCGGCGGTTGCCGCGGGGCGGAGCTCCGTGTGACGGTGAGCGGCGGCGGGGGGCTGTCGGGAGGAATGAAAGGCGGCGACGAAGGACTGACGGTGACTGACGAGCGAATGAAAGGAGGGAGTGAAGGGATGAAAGTGACTCACCGCTGAGTGAACGGCGGGGACGAGTGGATGAAAGTGACTCACCGCTGAGTGACAGGGAGGAGTGAAGGAATGAAAGTGACTCAGCAGTGAATGAAAGGTAGGAGTGAAGGAATGAAAGTGACTGACGAGTGAATGACAGGAAGGAGTGAAGGAATGAAAGTGACTCAGCAGTGAATGAAAGGAAGGAGTGAAGGAATGAAAATGACTCAGCAGTGAATGACAGGAGGGAGTCAAGGGATGAAAGTGATTCACGAGTGAATGACAGGAAGGATTGAAGGAATGAAAGTGACTGACAAGTGAATGACGGGAAGGAGTGAAGGAATGAAAATGACTGACGAGTGAATGAAAGGAAGGAGTGAGTGAATGAAAGTGACTCACCGCTGAATGACAGGAAGGAGTGAAGGAATGAAAGTGACTCACGAGTGATTGAAAGGATGGAGTGAAGGAATGAAAGTGACTCACGAGTGAATGAAAGGAAGGAGTGAAGGAATGAAAGTGACTCAGCAGTCAATGACAGGAAGGAGTGAAGGAATGAAAGTGACTCAGCAGTGAATGACAGGAAGGAGTGAAGGAATGAAAGTGACTCAGCAGTGAATGAAAGGAAGGAGTGAAGGAATGAAAGTGACTCAGCAGTGAATGACAGGAAGGAGTGAAGGAATGAAAGTGACTCACGAGTGAATGACAGGAAGGAGTGAAGGAATGAAAGGAAGGGACGAGTGAATGAAAGGCAGGAGTGAAGGAATGAAAGTGACTCAGCAGTGAATGAAAGGAGGGAGTGAAGGAATGAAAGTGACTTAGCAGTGAATGACAGGAAGGAGTGAAGGAATGAAAGTGACTCAGCAGTGAATGACAGGAAGGAGTGAAGGAATGAAAGTGACTCACGAGTGAATGAAAGGATGGAGTGAAGGAATGAAAGTGACTGACAAGTGAATGACAGGAAGGAGTGAAGGAATGAAAATGACTCAGCAGTGAATGACAGGAAGGAGTGAAGGAATGAAAGTGACTCACTAGTGAATGACAGGAAGGAGTGAAGGAATGAAAGGAGGGGACGAGTGAATGAAAGGAAGGAGTGAAGGAATGAAAGTGACTCACGAGTGAATGACAGGAAGGAATGAAGGAATGCAAGTGACTCACCGCTGAATGACAGGAAGGAGTGAAGGAATGAAAGGCGGGGACGAGTGAATGAAAGGAAGGAGTGAAGGAATGAAAGTGACTCAGCAGTGAATGACAGGAAGGAGTGAAGGAATGAAAGTGACTCACTAGTGAATGACAGGAAGGAGTGAAGGAATGAAAGGAGGGGACGAGTGAATGAAAGGAAGGAGTGAAGGAATGAAAGTGACTCACGAGTGAATGACAGGAAGGAGTGAAGGAATGAAAGGAGGGGACGAGTGAATGAAAGGAAGGAGTGAATGAATGAAAGTGACTGACCGCTGAATGACAGGAAGGAGTGAAGGAATGAAAGGAGGGGACGAGTGAATGAAAGGAAGGAGTGAAGGATTGAAAGTGATTAACTAGTGAATGAAAGGAAGGAGTGAAGGAATGAAAGTGACTCAGCAGCGAATGAAAGGCAGGAGTGAACGAGTGAAAGTGACTCAGCAGTGAATGAAAGGAAGGAGTGAAGGAATGAAAGTGACTGACGAGTGAATGACAGGCAGGAGTGACGGAATGAAAGTGAATGACGAGTGAATGAGAGCGAGGAGGGGTCGCGGCTGAGGCCGGGAGAGGGGGGGGCTCGCCTCTGACCTCTGACCCCGCCAACTAAGCCGCCCCTCCTCCTCCTCTGAATCTCGCGAGATGTTGACTCCCTCGCCTGGCACGCAGGAGGTCGTGAGTTCGACCCCGACCCCTCACGACGGACGCCTGCTGTATGTGTGGGAGTTTGCGGGTCTCTCTCTCTCTCCCCACGCGGTCGCGTGGATTCTTCTACCGGGGTCTCCGCGCTTAAAGAGTATTTGTCGGCTGTGAATTTGTCGGCGGGCTTTCATTTATTTGACCCGGGGAAGGCCCCCCCCCCCCCCCGACCCCAACTCGATTTTCAGAAGCGAACTTGCCCACCGCGAACGATCGCCCGGCGGAGCGGGCTCCCATCGCTCCCATCGTTCAAATTCGTTGCGACCCGCCGAACACTGTCAACGCAACGTGGGCCGTCGATTCGTGCCGTGGAAGGGGGGAAGGCACCGGACGAATCCACGCGTCCCACGGGCAGAGAGAGCGGGAGCCACGCAGACCCCCAAATATACAGCGGGCAACGTCAGGACGGGTCGGTCGGTCGTGATCGAACCCACGGAGACCTCCTGCGCACCAGGCGAGGTAGTTAACATCTCGAGCAAATGAACGAATGGGTGGGTGGGAACGGAGCGGATGACGGTCGGACGAATGGACGGACCACTCAGAAGTTTGGGTGAAGTTTTGGGATTTTTTTATTATTTTGCTGCCCCCCCATTTGGTTACTCGGGGTTAGCCCCCCCCACCTTCATCCCTACCCCCCCGCGGGGGTCGCGCACATGAAGGGGGGAGCAGAAACTACGCCGTAGGGGCTCTTGCTGGAACGAGGAGCGGTCACGGCGGGGGTTGGTCTGTTGGGGGGGGGGGGCGCCTTGGGTTATGGGCAGAGTGGTCGAGTCGGGCCACGCACACTCACACGTACACACACATGTACACACACACGCACACTTACACATACACGCACACTCACACACACATGTACACACACACGCACACTTACACATACACGCACACTCACACACACATGTACACACACACGCACACATACACGCACACTCACACGTACACACACGCACACACTTACACATACACACTCACACGTACACACACGCACACACACACATGTACACACACATGCACACTTACACATACAAGCACACACGCACACATACACACACACACACTTACACATACACACACACGTACACACACATACACACATACGCACGCACTTAAACACACACTTACACATACACACACTCACAAGCACATGTACACACACACACGCACACACTTACACATACACACACACGCATGTACACACACGCACACGTACACACACACACTTACACATACACACACACTTACACATACACACACACTCACACGCATGTACACACACGCACTCACACACGCGTACACACGCACGTACACACACGCACACACTTACACACGTACACACACACACCCATACACGCACGCACACGCGCACACACACACGCACACATACACACACGCACACTTGCACATACACGCACGTGCGCACACACACGCACGTACACACACGTACACACGCACACACACGCACACAGGCACACGCACGCACATACACGCACGCACATACGCGCATGCACACACGCGTACAACGGACACGCACACGCAGTATGTGACCGAAGGTCGATGTTTTTTTCTTTCTTAATGTTGATCTTGTTTCCTAGATGTAGTTCGCGCTGTCGGATGTGAACCGTTACTCGTGTTGTGAGAGGCTGTCGTGGGGATTGTGTACACACACGCACGTGCACACACGCACACACACGCGTACACACACACGCACACGCGTACACACACGCGTACACACACACGCACGCACACACGTACACACACACGCACGCACACACGCACACACGCGTACACACACACGCACACACACACGTACACACACACGTACACACACACGCATACACACACGTACACACACACGCACGCACACGCGTACACACACGCACACACACAAACCACAACGCACACACAAACTTGCCAGTGTCACGCGTACACACACACACACAAACCCGCACGTGTCACACACACATGTACACACACACAAACCCACGTGTCACACACACATGTACACACACACACGTGTAGACACACACGCGTACACACACACGCGTACACACACACGCGTCACACTCGCGTCACACACACATGTAGACACACACACACAAACCCGCACGCGTCACACACACATGTACACACACACACATGTACACACACACACACGCGTACACACACACACACATGTACACACGCACGCGTAGACACACACACACAAACCCGCACGCGTCACACACACATGTACACACACACACATGTACACACACACGTGTACACACACACACATGTACACACACACATGTACACACACACACACATGTACACACACGCGCACGACCGCTCCCCACGACATCCCAGCGGACGACGCGTACAACTTGTTCTTAATTCTCTCTCTCTAGTCCATGAAGTGGTTTAAACATTTAAAAACAAAATAAAAAGTTAGTGGGCGACAGCGACGGCTTTCTTCTCGCCCCCCCTCCCCCCCATCGTGGGGGTGCGGAGCGGGGGAGGGGGTCACACCAAAATCGTGCGCGAGCCACAGCGGCGAGGAAATGTTAACTCCCTCGCCTGGAGTGCAGGAGGTCCGTGGCTTCGACGCTCGATTCCTGACCCTCCCCCCCCGCTCCCCCACGAGACGTTGACCGCCCGTTTCCAGCCTGGGTTCGATTCCCGGCGGGCAGGAGGGGGGCAGGCGAGAGGGAACCGGTGAAAGGTAGCGTCGACACACGTCGCGATCACGCGTGCCGAGGATGAACCTCCGAGCGGTCGGACGGGACGGGACGCGACGGGACGCGACGGATCGACTGAACTTGACGGATTCGACCGTTCGAGTCGCGGACGCGGCGGCACCGGCGACCTCCCCACGTGCATGAAGGGCTCCCGGGAAAAGCGCGACCACGTTACCGGGACTCCGGATGGCGAGCGAGTGAGTCGCGATGGGCTCGCCGTGCCGCCGTGTCCGTGGTGGTGGTGGTGGGAGGGCGGCGATGAAGAGACGCGTGTTAGAGACTCCCACACCGCCGTGCGGTTGTGTGTGCGGGTGGTTGTTCGGACTGTGTCGGGGGGCCTCAATAAAATCTTGAGACGCTGAGCGTTTGGTGAGCGCGTGTTTATTTCGCGACGCGTTGGCGTTGTTTGTTGGTGTTGTTGGTTGTGGTTTGTTGTTGTTGTGTGTCCCAGTGCTACGCTGCGCGACGAACCTGGCGAGACGGGTTCGATTCCTGATCACGGCCAACATCGGTGACGATTATCCGAACGGGTCCTGGGCCTCCGCTAGGTCGACGCAGCCTCAAATGAATACCTGGGGCGGTGCGCGAACATCGCCACTTGGTATAGCCATGGAAAGCGTCTCGAACTACGTGGTCAAAGATGATGTTAAGAAGTGTGGGAGCCGGAGGACATCCCTGTCGCATCCCGAAGGTGTACAGGGAATGGCTCCGACTCCTGGCCACGTAGATCTACCCGGACCTGCCGTGCCCACCATCACGAAGGCCCTTTAATGATTGCGAGCAGAGAGTCAGGGACTCCGAAATCACGAAGAATTGCCCACAATTGTGGCCTGTTGATGGTATCATAGGCCTTGACCAGGTCGATAAAGCAGAGGTGTAGGTCTTGTTGGAATTCCCTACACCTCTGCTGGAGCAGATGGAGAGTGAACGTGGCTTTTGTGGTCTGAAACCGCATTGGGGCTCCGCAAGCCTCTCCTCAGCGTGCCTGGCAAGGCGATGTTGAATAATCCTTGCCAGGACCTTTCCCGGCACAAGACAAGACACACACACATATAAAAAACATTTCACTCAAGCTCGCCGTTAATCAGTGGGCCAACGCGGCAGCGAAAATGAATCGGGGTTCAAGTTCATTTATTAGGTATAGCCCTGTGAGCCATTCGGCTCTTGAATCGGGTGCAACCGCAACAAACAAAAAACAAAAACATGAACAAGAAAACATCTTAAAGTGAATATGTGACTGAGTGCAAACAGAAAATCCAAGAAAAGACAGCAAAATGTTGCAGAAAAAATCACCGTACACCAACGCTGTGTGCACAAATCCCAGTGGGATGCAAGTCAAACAACAACAACCACAAGGCATCTTCTTGGTTCTTTCGGTAAAGTCACGTAGAAGGGAA

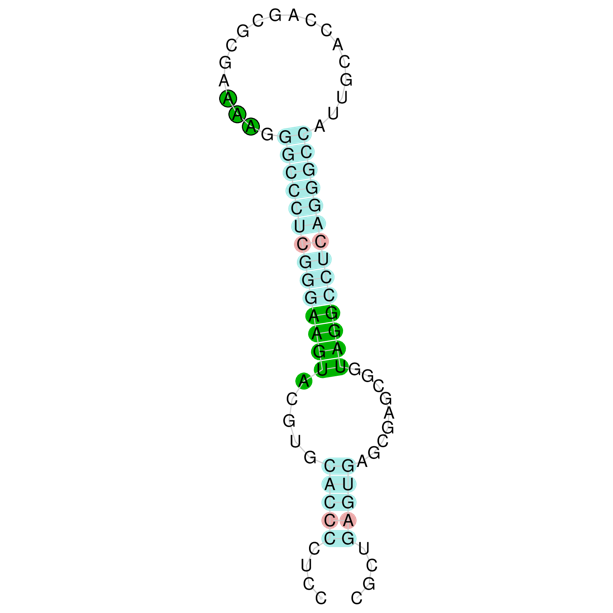

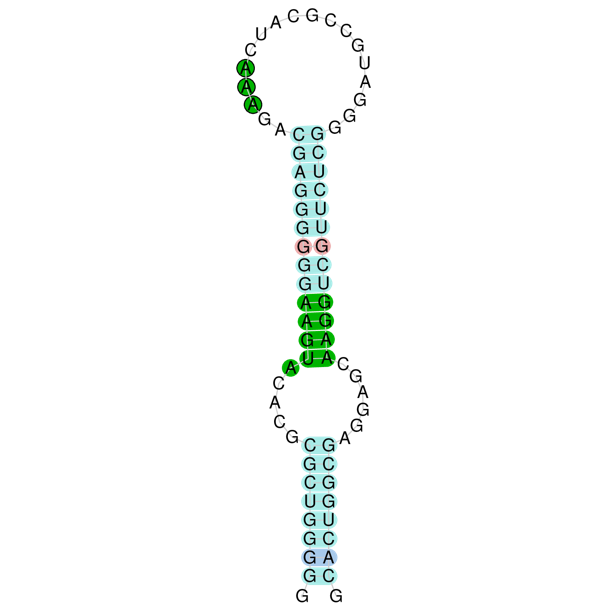

### List of RNA matches identifiers

#### TSA ids matching Lethenteron_reissneri_SELENOP_nt, indicated with their respective matched position range:

GIWK01068433.1:589-1926 GIWK01068433.1:762-1923 GIWK01068433.1:745-1926 GIWK01068433.1:745-1926 GIWK01068433.1:762-1926 GIWK01068433.1:742-1926 GIWK01068433.1:774-1923 GIWK01068433.1:762-1926 GIWK01068433.1:762-1926 GIWK01068433.1:762-1926 GIWK01068433.1:829-1926 GIWK01068433.1:745-1923 GIWK01068433.1:747-1926 GIWK01068433.1:745-1827 GIWK01068433.1:762-1926 GIWK01068433.1:762-1810 GIWK01068433.1:888-1905 GIWK01068433.1:783-1905 GIWK01068433.1:913-1926 GIWK01068433.1:745-1768 GIWK01068433.1:984-1905 GIWK01068433.1:1014-1926 GIWK01068433.1:762-1684 GIWK01068433.1:1077-1926 GIWK01068433.1:780-1600 GIWK01068433.1:2-586 GIWK01068433.1:1140-1926 GIWK01068433.1:1165-1926 GIWK01068433.1:1224-1926 GIWK01068433.1:742-1474 GIWK01068433.1:1249-1926 GIWK01068433.1:1291-1926 GIWK01068433.1:762-1348 GIWK01068433.1:762-1306 GIWK01068433.1:1417-1923 GIWK01053646.1:1961-3156 GIWK01053646.1:1961-3156 GIWK01053646.1:1961-3154 GIWK01053646.1:1961-3156 GIWK01053646.1:1961-3154 GIWK01053646.1:1961-3154 GIWK01053646.1:1961-3154 GIWK01053646.1:1964-3156 GIWK01053646.1:1961-3135 GIWK01053646.1:1961-3152 GIWK01053646.1:1961-3154 GIWK01053646.1:1961-3156 GIWK01053646.1:1961-3154 GIWK01053646.1:1961-3156 GIWK01053646.1:1961-3093 GIWK01053646.1:1961-3051 GIWK01053646.1:1989-3156 GIWK01053646.1:2060-3149 GIWK01053646.1:2132-3149 GIWK01053646.1:2153-3152 GIWK01053646.1:2090-3156 GIWK01053646.1:1961-2903 GIWK01053646.1:1961-2967 GIWK01053646.1:1961-2886 GIWK01053646.1:2216-3149 GIWK01053646.1:2241-3156 GIWK01053646.1:2300-3156 GIWK01053646.1:1961-2844 GIWK01053646.1:2325-3156 GIWK01053646.1:2367-3156 GIWK01053646.1:1961-2760 GIWK01053646.1:1961-2676 GIWK01053646.1:2493-3156 GIWK01053646.1:1961-2550 GIWK01053646.1:2678-3156 GIWK01053646.1:2780-3156 GIWK01063328.1:1-1464 GIWK01063328.1:302-1463 GIWK01063328.1:285-1464 GIWK01063328.1:282-1464 GIWK01063328.1:285-1464 GIWK01063328.1:302-1464 GIWK01063328.1:314-1463 GIWK01063328.1:302-1464 GIWK01063328.1:302-1464 GIWK01063328.1:369-1464 GIWK01063328.1:302-1464 GIWK01063328.1:287-1464 GIWK01063328.1:285-1463 GIWK01063328.1:285-1367 GIWK01063328.1:302-1464 GIWK01063328.1:302-1350 GIWK01063328.1:428-1445 GIWK01063328.1:302-1431 GIWK01063328.1:323-1445 GIWK01063328.1:453-1464 GIWK01063328.1:285-1308 GIWK01063328.1:524-1445 GIWK01063328.1:554-1464 GIWK01063328.1:302-1224 GIWK01063328.1:596-1463 GIWK01063328.1:617-1464 GIWK01063328.1:320-1140 GIWK01063328.1:680-1464 GIWK01063328.1:705-1464 GIWK01063328.1:764-1464 GIWK01063328.1:282-1014 GIWK01063328.1:789-1464 GIWK01063328.1:831-1464 GIWK01063328.1:302-888 GIWK01063328.1:302-846 GIWK01063328.1:957-1463 GIWK01066363.1:857-1435 GIWK01066363.1:851-1433 GIWK01066363.1:851-1433 GIWK01066363.1:858-1426 GIWK01066363.1:857-1431 GIWK01066363.1:851-1433 GIWK01066363.1:857-1433 GIWK01066363.1:857-1429 GIWK01066363.1:864-1433 GIWK01066363.1:851-1433 GIWK01066363.1:851-1431 GIWK01066363.1:851-1433 GIWK01066363.1:857-1431 GIWK01066363.1:857-1433 GIWK01066363.1:857-1431 GIWK01066363.1:857-1426 GIWK01066363.1:857-1433 GIWK01066363.1:857-1409 GIWK01066363.1:851-1370 GIWK01066363.1:851-1431 GIWK01066363.1:913-1433 GIWK01066363.1:1015-1431 GIWK01066363.1:1183-1433 GIWK01011613.1:191-2057 GIWK01011613.1:1546-2057 GIWK01011613.1:1566-2040 GIWK01011613.1:1566-2057 GIWK01011613.1:1549-2040 GIWK01011613.1:1566-2057 GIWK01011613.1:1566-2057 GIWK01011613.1:1566-2057 GIWK01011613.1:1566-2057 GIWK01011613.1:1566-2057 GIWK01011613.1:1566-2057 GIWK01011613.1:1566-2057 GIWK01011613.1:1584-2057 GIWK01011613.1:1546-2048 GIWK01011613.1:1549-2057 GIWK01011613.1:1566-2057 GIWK01011613.1:1633-2057 GIWK01011613.1:163-198 GIWK01059119.1:191-1715 GIWK01059119.1:1566-1709 GIWK01059119.1:1566-1715 GIWK01059119.1:1566-1715 GIWK01059119.1:1549-1712 GIWK01059119.1:1566-1712 GIWK01059119.1:1566-1715 GIWK01059119.1:163-198 GIWK01065380.1:163-1303 GIWK01065380.1:1290-1623 GIWK01065380.1:1473-1616 GIWK01065380.1:1473-1622 GIWK01065380.1:1473-1622 GIWK01065380.1:1456-1619 GIWK01065380.1:1473-1619 GIWK01065380.1:1473-1622 GIWK01067313.1:1-1359 GIWK01067313.1:1-1355 GIWK01067313.1:1-1353 GIWK01067313.1:1-1334 GIWK01067313.1:7-1353 GIWK01067313.1:1-1359 GIWK01067313.1:1-1355 GIWK01067313.1:1-1353 GIWK01067313.1:1-1353 GIWK01067313.1:1-1355 GIWK01067313.1:1-1292 GIWK01067313.1:62-1353 GIWK01067313.1:37-1353 GIWK01067313.1:121-1359 GIWK01067313.1:1-1250 GIWK01067313.1:1-1208 GIWK01067313.1:146-1353 GIWK01067313.1:188-1353 GIWK01067313.1:1-1141 GIWK01067313.1:1-1121 GIWK01067313.1:226-1348 GIWK01067313.1:1-1079 GIWK01067313.1:1-1040 GIWK01067313.1:314-1348 GIWK01067313.1:352-1351 GIWK01067313.1:1-995 GIWK01067313.1:1-956 GIWK01067313.1:457-1353 GIWK01067313.1:398-1348 GIWK01067313.1:1-914 GIWK01067313.1:1-872 GIWK01067313.1:499-1353 GIWK01067313.1:524-1353 GIWK01067313.1:13-724 GIWK01067313.1:601-1353 GIWK01067313.1:685-1353 GIWK01067313.1:1-788 GIWK01067313.1:1-707 GIWK01067313.1:769-1359 GIWK01067313.1:13-665 GIWK01067313.1:1-581 GIWK01067313.1:1-497 GIWK01067313.1:874-1353 GIWK01067313.1:958-1353 GIWK01067313.1:1105-1351 GIWK01067621.1:699-1494 GIWK01067621.1:2-708 GIWK01015682.1:1-1258 GIWK01015682.1:1-1375 GIWK01015682.1:1-1342 GIWK01015682.1:1-1374 GIWK01015682.1:1-1374 GIWK01015682.1:28-1375 GIWK01015682.1:1-1375 GIWK01015682.1:1-1375 GIWK01015682.1:1-1375 GIWK01015682.1:1-1375 GIWK01015682.1:1-1374 GIWK01015682.1:3-1374 GIWK01015682.1:1-1375 GIWK01015682.1:1-1165 GIWK01015682.1:1-1300 GIWK01015682.1:66-1375 GIWK01015682.1:154-1375 GIWK01015682.1:192-1375 GIWK01015682.1:1-1132 GIWK01015682.1:1-1048 GIWK01015682.1:1-1081 GIWK01015682.1:1-1006 GIWK01015682.1:1-964 GIWK01015682.1:339-1356 GIWK01015682.1:441-1356 GIWK01015682.1:364-1375 GIWK01015682.1:525-1374 GIWK01015682.1:1-880 GIWK01015682.1:1-796 GIWK01015682.1:528-1375 GIWK01015682.1:1-835 GIWK01015682.1:609-1375 GIWK01015682.1:606-1375 GIWK01015682.1:1-754 GIWK01015682.1:1-712 GIWK01015682.1:1-564 GIWK01015682.1:1-628 GIWK01015682.1:1-547 GIWK01015682.1:798-1374 GIWK01015682.1:1-505 GIWK01015682.1:823-1374 GIWK01015682.1:903-1375 GIWK01015682.1:966-1375 GIWK01015682.1:1092-1375 GIWK01066786.1:607-1455 GIWK01066786.1:3-682 GIWK01015113.1:992-1912 GIWK01015113.1:981-1910 GIWK01015113.1:992-1908 GIWK01015113.1:992-1910 GIWK01015113.1:985-1908 GIWK01015113.1:985-1906 GIWK01015113.1:985-1908 GIWK01015113.1:992-1903 GIWK01015113.1:992-1903 GIWK01015113.1:985-1912 GIWK01015113.1:985-1912 GIWK01015113.1:985-1903 GIWK01015113.1:995-1910 GIWK01015113.1:985-1908 GIWK01015113.1:985-1910 GIWK01015113.1:985-1910 GIWK01015113.1:992-1908 GIWK01015113.1:985-1908 GIWK01015113.1:985-1910 GIWK01015113.1:985-1889 GIWK01015113.1:992-1808 GIWK01015113.1:1054-1912 GIWK01015113.1:989-1766 GIWK01015113.1:985-1825 GIWK01015113.1:1138-1910 GIWK01015113.1:1222-1910 GIWK01015113.1:1180-1912 GIWK01015113.1:985-1682 GIWK01015113.1:993-1640 GIWK01015113.1:1276-1912 GIWK01015113.1:1306-1912 GIWK01015113.1:992-1511 GIWK01015113.1:985-1472 GIWK01015113.1:1415-1910 GIWK01031270.1:1-1242 GIWK01031270.1:9-1475 GIWK01031270.1:1-1473 GIWK01031270.1:9-1471 GIWK01031270.1:12-1471 GIWK01031270.1:5-1471 GIWK01031270.1:9-1452 GIWK01031270.1:9-1469 GIWK01031270.1:50-1473 GIWK01031270.1:9-1471 GIWK01031270.1:9-1326 GIWK01031270.1:1-1365 GIWK01031270.1:1-1410 GIWK01031270.1:138-1475 GIWK01031270.1:113-1473 GIWK01031270.1:1-1284 GIWK01031270.1:176-1473 GIWK01031270.1:1-1158 GIWK01031270.1:281-1475 GIWK01031270.1:222-1471 GIWK01031270.1:425-1473 GIWK01031270.1:1-1116 GIWK01031270.1:1-1023 GIWK01031270.1:323-1475 GIWK01031270.1:593-1475 GIWK01031270.1:509-1469 GIWK01031270.1:1-1074 GIWK01031270.1:348-1466 GIWK01031270.1:1-990 GIWK01031270.1:428-1466 GIWK01031270.1:512-1466 GIWK01031270.1:1-939 GIWK01031270.1:1-864 GIWK01031270.1:698-1473 GIWK01031270.1:656-1473 GIWK01031270.1:9-819 GIWK01031270.1:9-780 GIWK01031270.1:752-1475 GIWK01031270.1:9-738 GIWK01031270.1:1-696 GIWK01031270.1:782-1473 GIWK01031270.1:807-1475 GIWK01031270.1:1-548 GIWK01031270.1:1-612 GIWK01031270.1:9-531 GIWK01031270.1:950-1475 GIWK01031270.1:926-1471 GIWK01031270.1:1-489 GIWK01031270.1:1010-1473 GIWK01031270.1:1034-1471 GIWK01031270.1:1076-1473 GIWK01031270.1:1143-1471 GIWK01029243.1:656-1528 GIWK01029243.1:20-750 GIWK01068541.1:1-1265 GIWK01068541.1:1-1265 GIWK01068541.1:1-1271 GIWK01068541.1:1-1265 GIWK01068541.1:12-1267 GIWK01068541.1:12-1271 GIWK01068541.1:1-1271 GIWK01068541.1:12-1267 GIWK01068541.1:30-1271 GIWK01068541.1:15-1265 GIWK01068541.1:1-1265 GIWK01068541.1:1-1267 GIWK01068541.1:1-1265 GIWK01068541.1:12-1246 GIWK01068541.1:12-1204 GIWK01068541.1:55-1265 GIWK01068541.1:12-1162 GIWK01068541.1:114-1265 GIWK01068541.1:12-1120 GIWK01068541.1:156-1260 GIWK01068541.1:240-1260 GIWK01068541.1:210-1265 GIWK01068541.1:12-984 GIWK01068541.1:282-1263 GIWK01068541.1:12-1033 GIWK01068541.1:307-1260 GIWK01068541.1:366-1265 GIWK01068541.1:12-952 GIWK01068541.1:12-910 GIWK01068541.1:391-1265 GIWK01068541.1:433-1265 GIWK01068541.1:12-826 GIWK01068541.1:12-742 GIWK01068541.1:1-700 GIWK01068541.1:471-1265 GIWK01068541.1:559-1265 GIWK01068541.1:702-1271 GIWK01068541.1:12-616 GIWK01068541.1:12-574 GIWK01068541.1:1-532 GIWK01068541.1:769-1265 GIWK01068541.1:849-1265 GIWK01068541.1:1017-1263 GIWK01056333.1:35-1183 GIWK01056333.1:1-993 GIWK01056333.1:1-1180 GIWK01056333.1:1-1180 GIWK01056333.1:1-1178 GIWK01056333.1:1-1182 GIWK01056333.1:1-1183 GIWK01056333.1:1-1182 GIWK01056333.1:1-1182 GIWK01056333.1:1-1180 GIWK01056333.1:1-1180 GIWK01056333.1:1-1180 GIWK01056333.1:1-1080 GIWK01056333.1:1-1180 GIWK01056333.1:1-1182 GIWK01056333.1:1-1097 GIWK01056333.1:1-1183 GIWK01056333.1:1-1161 GIWK01056333.1:102-1182 GIWK01056333.1:179-1175 GIWK01056333.1:77-1175 GIWK01056333.1:1-1038 GIWK01056333.1:263-1175 GIWK01056333.1:1-912 GIWK01056333.1:182-1178 GIWK01056333.1:266-1182 GIWK01056333.1:347-1182 GIWK01056333.1:1-870 GIWK01056333.1:344-1183 GIWK01056333.1:12-828 GIWK01056333.1:410-1182 GIWK01056333.1:452-1183 GIWK01056333.1:12-744 GIWK01056333.1:506-1182 GIWK01056333.1:1-702 GIWK01056333.1:1-618 GIWK01056333.1:536-1183 GIWK01056333.1:561-1183 GIWK01056333.1:1-576 GIWK01056333.1:1-534 GIWK01056333.1:1-492 GIWK01056333.1:687-1182 GIWK01050416.1:1-782 GIWK01050416.1:1-770 GIWK01050416.1:1-770 GIWK01050416.1:1-770 GIWK01050416.1:1-770 GIWK01050416.1:1-782 GIWK01050416.1:1-770 GIWK01050416.1:1-770 GIWK01050416.1:1-770 GIWK01050416.1:1-770 GIWK01050416.1:1-770 GIWK01050416.1:1-770 GIWK01050416.1:1-770 GIWK01050416.1:1-770 GIWK01050416.1:1-770 GIWK01050416.1:1-770 GIWK01050416.1:1-770 GIWK01050416.1:1-770 GIWK01050416.1:1-771 GIWK01050416.1:1-770 GIWK01050416.1:30-769 GIWK01050416.1:1-770 GIWK01050416.1:1-756 GIWK01050416.1:2-672 GIWK01050416.1:1-630 GIWK01050416.1:114-769 GIWK01050416.1:1-546 GIWK01050416.1:215-770 GIWK01050416.1:272-769 GIWK01050416.1:275-770 GIWK01050416.1:356-770 GIWK01050416.1:359-770 GIWK01020752.1:1-1440 GIWK01020752.1:1-1440 GIWK01020752.1:8-1440 GIWK01020752.1:1-1440 GIWK01020752.1:1-1440 GIWK01020752.1:1-1438 GIWK01020752.1:1-1421 GIWK01020752.1:1-1440 GIWK01020752.1:1-1440 GIWK01020752.1:20-1440 GIWK01020752.1:1-1372 GIWK01020752.1:121-1440 GIWK01020752.1:1-1286 GIWK01020752.1:1-1253 GIWK01020752.1:79-1440 GIWK01020752.1:1-1211 GIWK01020752.1:1-1043 GIWK01020752.1:146-1440 GIWK01020752.1:1-1127 GIWK01020752.1:226-1440 GIWK01020752.1:1-1169 GIWK01020752.1:223-1440 GIWK01020752.1:310-1435 GIWK01020752.1:307-1440 GIWK01020752.1:1-1082 GIWK01020752.1:373-1440 GIWK01020752.1:1-959 GIWK01020752.1:1-1001 GIWK01020752.1:398-1435 GIWK01020752.1:440-1438 GIWK01020752.1:478-1435 GIWK01020752.1:1-917 GIWK01020752.1:541-1440 GIWK01020752.1:566-1440 GIWK01020752.1:604-1440 GIWK01020752.1:1-791 GIWK01020752.1:17-749 GIWK01020752.1:650-1440 GIWK01020752.1:1-623 GIWK01020752.1:18-662 GIWK01020752.1:1-578 GIWK01020752.1:751-1440 GIWK01020752.1:776-1440 GIWK01020752.1:853-1440 GIWK01020752.1:1-539 GIWK01020752.1:937-1440 GIWK01020752.1:940-1440 GIWK01044708.1:129-1410 GIWK01044708.1:1-1410 GIWK01044708.1:1-1406 GIWK01044708.1:6-1409 GIWK01044708.1:1-1409 GIWK01044708.1:1-1410 GIWK01044708.1:1-1406 GIWK01044708.1:1-1404 GIWK01044708.1:1-1387 GIWK01044708.1:1-1406 GIWK01044708.1:1-1412 GIWK01044708.1:1-1345 GIWK01044708.1:63-1412 GIWK01044708.1:1-1278 GIWK01044708.1:66-1412 GIWK01044708.1:1-1258 GIWK01044708.1:1-1209 GIWK01044708.1:171-1406 GIWK01044708.1:213-1412 GIWK01044708.1:1-1177 GIWK01044708.1:234-1412 GIWK01044708.1:1-1132 GIWK01044708.1:1-1093 GIWK01044708.1:297-1401 GIWK01044708.1:1-1051 GIWK01044708.1:339-1410 GIWK01044708.1:444-1401 GIWK01044708.1:381-1401 GIWK01044708.1:1-967 GIWK01044708.1:507-1409 GIWK01044708.1:532-1412 GIWK01044708.1:591-1409 GIWK01044708.1:1-715 GIWK01044708.1:1-841 GIWK01044708.1:1-796 GIWK01044708.1:616-1409 GIWK01044708.1:658-1412 GIWK01044708.1:1-757 GIWK01044708.1:1-589 GIWK01044708.1:927-1412 GIWK01044708.1:3-673 GIWK01044708.1:1-631 GIWK01044708.1:784-1412 GIWK01044708.1:1-547 GIWK01044708.1:994-1409 GIWK01069519.1:636-1657 GIWK01064888.1:439-1456 GIWK01064888.1:1-683 GIWK01052658.1:33-1068 GIWK01052658.1:1285-2258 GIWK01052658.1:814-1853 GIWK01011755.1:1-722 GIWK01011755.1:3-722 GIWK01011755.1:3-722 GIWK01011755.1:1-722 GIWK01011755.1:3-722 GIWK01011755.1:3-722 GIWK01011755.1:3-722 GIWK01011755.1:3-722 GIWK01011755.1:1-722 GIWK01011755.1:3-722 GIWK01011755.1:3-722 GIWK01011755.1:1-722 GIWK01011755.1:1-722 GIWK01011755.1:3-722 GIWK01011755.1:3-720 GIWK01011755.1:3-722 GIWK01011755.1:3-722 GIWK01011755.1:1-722 GIWK01011755.1:197-722 GIWK01011755.1:3-722 GIWK01011755.1:3-722 GIWK01011755.1:3-636 GIWK01011755.1:92-722 GIWK01011755.1:3-569 GIWK01011755.1:3-549 GIWK01011755.1:3-500 GIWK01011755.1:285-722 GIWK01019425.1:731-1992 GIWK01019425.1:1-1221 GIWK01064514.1:1-1366 GIWK01064514.1:1-1388 GIWK01064514.1:1-1388 GIWK01064514.1:1-1388 GIWK01064514.1:1-1385 GIWK01064514.1:1-1388 GIWK01064514.1:1-1307 GIWK01064514.1:1-1386 GIWK01064514.1:322-1389 GIWK01064514.1:1-1388 GIWK01064514.1:1-1349 GIWK01064514.1:77-1388 GIWK01064514.1:22-1388 GIWK01064514.1:1-1223 GIWK01064514.1:52-1386 GIWK01064514.1:1-1181 GIWK01064514.1:220-1388 GIWK01064514.1:119-1388 GIWK01064514.1:1-1139 GIWK01064514.1:157-1388 GIWK01064514.1:1-1052 GIWK01064514.1:406-1388 GIWK01064514.1:346-1363 GIWK01064514.1:1-1097 GIWK01064514.1:430-1363 GIWK01064514.1:1-1013 GIWK01064514.1:1-929 GIWK01064514.1:472-1388 GIWK01064514.1:514-1388 GIWK01064514.1:574-1388 GIWK01064514.1:1-887 GIWK01064514.1:658-1386 GIWK01064514.1:577-1388 GIWK01064514.1:661-1386 GIWK01064514.1:1-697 GIWK01064514.1:739-1386 GIWK01064514.1:763-1388 GIWK01064514.1:1-680 GIWK01064514.1:1-593 GIWK01064514.1:1-505 GIWK01064514.1:788-1388 GIWK01064514.1:830-1388 GIWK01064514.1:956-1385 GIWK01006535.1:1013-2127 GIWK01006535.1:1-1299 GIWK01032908.1:1-1690 GIWK01032908.1:1-346 GIWK01032908.1:1-346 GIWK01032908.1:1-344 GIWK01032908.1:1-346 GIWK01032908.1:2-344 GIWK01032908.1:1-346 GIWK01032908.1:1-346 GIWK01032908.1:1-339 GIWK01032908.1:1-346 GIWK01032908.1:1-346 GIWK01032908.1:1-321 GIWK01032908.1:1-344 GIWK01032908.1:1-346 GIWK01026969.1:1195-2333 GIWK01026969.1:1-866 GIWK01026969.1:397-1546 GIWK01000094.1:235-1376 GIWK01000094.1:1-592 GIWK01014775.1:89-1487 GIWK01014775.1:11-1487 GIWK01014775.1:5-1487 GIWK01014775.1:7-1481 GIWK01014775.1:9-1487 GIWK01014775.1:5-1445 GIWK01014775.1:6-1487 GIWK01014775.1:5-1487 GIWK01014775.1:5-1487 GIWK01014775.1:11-1403 GIWK01014775.1:8-1487 GIWK01014775.1:92-1487 GIWK01014775.1:155-1487 GIWK01014775.1:197-1487 GIWK01014775.1:5-1319 GIWK01014775.1:6-1277 GIWK01014775.1:222-1487 GIWK01014775.1:9-1210 GIWK01014775.1:281-1487 GIWK01014775.1:306-1487 GIWK01014775.1:390-1484 GIWK01014775.1:9-1141 GIWK01014775.1:801-1487 GIWK01014775.1:9-1025 GIWK01014775.1:533-1487 GIWK01014775.1:474-1487 GIWK01014775.1:449-1487 GIWK01014775.1:6-1064 GIWK01014775.1:1-800 GIWK01014775.1:6-983 GIWK01014775.1:9-802 GIWK01014775.1:558-1487 GIWK01014775.1:600-1487 GIWK01014775.1:1-738 GIWK01014775.1:6-783 GIWK01014775.1:638-1487 GIWK01014775.1:726-1487 GIWK01014775.1:9-615 GIWK01014775.1:9-573 GIWK01014775.1:926-1487 GIWK01014775.1:5-531 GIWK01014775.1:1010-1487 GIWK01014775.1:1081-1487 GIWK01014775.1:1174-1487 GIWK01015959.1:9-1424 GIWK01015959.1:8-1365 GIWK01015959.1:9-1424 GIWK01015959.1:11-1424 GIWK01015959.1:47-1424 GIWK01015959.1:14-1424 GIWK01015959.1:11-1410 GIWK01015959.1:9-1424 GIWK01015959.1:9-1424 GIWK01015959.1:9-1424 GIWK01015959.1:383-1435 GIWK01015959.1:9-1284 GIWK01015959.1:9-1326 GIWK01015959.1:131-1424 GIWK01015959.1:50-1424 GIWK01015959.1:128-1424 GIWK01015959.1:9-1242 GIWK01015959.1:6-1200 GIWK01015959.1:194-1424 GIWK01015959.1:236-1424 GIWK01015959.1:320-1424 GIWK01015959.1:290-1424 GIWK01015959.1:9-1116 GIWK01015959.1:9-1155 GIWK01015959.1:446-1424 GIWK01015959.1:488-1435 GIWK01015959.1:8-990 GIWK01015959.1:548-1424 GIWK01015959.1:9-1032 GIWK01015959.1:9-948 GIWK01015959.1:1-755 GIWK01015959.1:572-1424 GIWK01015959.1:9-906 GIWK01015959.1:632-1424 GIWK01015959.1:9-822 GIWK01015959.1:740-1424 GIWK01015959.1:698-1423 GIWK01015959.1:782-1423 GIWK01015959.1:11-696 GIWK01015959.1:10-645 GIWK01015959.1:807-1424 GIWK01015959.1:9-612 GIWK01015959.1:866-1424 GIWK01015959.1:9-528 GIWK01015959.1:891-1423 GIWK01015959.1:933-1424 GIWK01015959.1:11-486 GIWK01025856.1:1-1246 GIWK01025856.1:783-2027 GIWK01062408.1:2-1501 GIWK01062408.1:2-1521 GIWK01062408.1:1-1462 GIWK01062408.1:4-1521 GIWK01062408.1:7-1501 GIWK01062408.1:2-1445 GIWK01062408.1:2-1516 GIWK01062408.1:2-1521 GIWK01062408.1:379-1524 GIWK01062408.1:40-1521 GIWK01062408.1:64-1521 GIWK01062408.1:106-1521 GIWK01062408.1:2-1358 GIWK01062408.1:148-1501 GIWK01062408.1:190-1521 GIWK01062408.1:2-1319 GIWK01062408.1:257-1521 GIWK01062408.1:2-1109 GIWK01062408.1:316-1521 GIWK01062408.1:2-1235 GIWK01062408.1:232-1521 GIWK01062408.1:2-1277 GIWK01062408.1:376-1521 GIWK01062408.1:2-1193 GIWK01062408.1:460-1521 GIWK01062408.1:2-1148 GIWK01062408.1:457-1521 GIWK01062408.1:2-1064 GIWK01062408.1:523-1521 GIWK01062408.1:2-1025 GIWK01062408.1:2-983 GIWK01062408.1:607-1521 GIWK01062408.1:2-934 GIWK01062408.1:628-1521 GIWK01062408.1:2-848 GIWK01062408.1:2-815 GIWK01062408.1:2-773 GIWK01062408.1:775-1521 GIWK01062408.1:4-689 GIWK01062408.1:859-1521 GIWK01062408.1:835-1524 GIWK01062408.1:3-644 GIWK01062408.1:901-1521 GIWK01062408.1:943-1521 GIWK01062408.1:2-563 GIWK01062408.1:2-521 GIWK01062408.1:1052-1521 GIWK01059296.1:3-1521 GIWK01059296.1:7-1521 GIWK01059296.1:1-1521 GIWK01059296.1:2-1490 GIWK01059296.1:2-1521 GIWK01059296.1:2-1512 GIWK01059296.1:5-1521 GIWK01059296.1:50-1521 GIWK01059296.1:2-1448 GIWK01059296.1:7-1381 GIWK01059296.1:5-1028 GIWK01059296.1:109-1521 GIWK01059296.1:1-1361 GIWK01059296.1:211-1521 GIWK01059296.1:298-1521 GIWK01059296.1:2-1312 GIWK01059296.1:5-1235 GIWK01059296.1:1-1280 GIWK01059296.1:134-1521 GIWK01059296.1:214-1521 GIWK01059296.1:295-1521 GIWK01059296.1:4-1196 GIWK01059296.1:376-1521 GIWK01059296.1:379-1504 GIWK01059296.1:5-1154 GIWK01059296.1:5-1112 GIWK01059296.1:484-1504 GIWK01059296.1:1-1067 GIWK01059296.1:1-944 GIWK01059296.1:568-1504 GIWK01059296.1:538-1521 GIWK01059296.1:1-986 GIWK01059296.1:593-1521 GIWK01059296.1:635-1521 GIWK01059296.1:673-1521 GIWK01059296.1:5-902 GIWK01059296.1:1-860 GIWK01059296.1:736-1521 GIWK01059296.1:778-1521 GIWK01059296.1:2-776 GIWK01059296.1:2-734 GIWK01059296.1:1009-1521 GIWK01059296.1:845-1521 GIWK01059296.1:2-650 GIWK01059296.1:883-1521 GIWK01059296.1:5-605 GIWK01059296.1:5-566 GIWK01059296.1:2-524 GIWK01059296.1:7-482 GIWK01059296.1:1097-1521 GIWK01000457.1:4-1352 GIWK01000457.1:4-1352 GIWK01000457.1:4-1339 GIWK01000457.1:1-1352 GIWK01000457.1:7-1352 GIWK01000457.1:1-1352 GIWK01000457.1:5-1352 GIWK01000457.1:4-1352 GIWK01000457.1:4-1339 GIWK01000457.1:4-1339 GIWK01000457.1:5-1280 GIWK01000457.1:22-1352 GIWK01000457.1:47-1339 GIWK01000457.1:4-1196 GIWK01000457.1:442-1352 GIWK01000457.1:89-1352 GIWK01000457.1:4-1154 GIWK01000457.1:127-1352 GIWK01000457.1:190-1352 GIWK01000457.1:4-1028 GIWK01000457.1:376-1352 GIWK01000457.1:232-1336 GIWK01000457.1:292-1339 GIWK01000457.1:4-1070 GIWK01000457.1:316-1336 GIWK01000457.1:400-1336 GIWK01000457.1:4-986 GIWK01000457.1:484-1352 GIWK01000457.1:4-902 GIWK01000457.1:568-1352 GIWK01000457.1:509-1352 GIWK01000457.1:4-860 GIWK01000457.1:593-1352 GIWK01000457.1:664-1352 GIWK01000457.1:1-692 GIWK01000457.1:4-647 GIWK01000457.1:736-1352 GIWK01000457.1:757-1352 GIWK01000457.1:4-566 GIWK01000457.1:1-475 GIWK01000457.1:845-1339 GIWK01000457.1:820-1352 GIWK01000457.1:904-1352 GIWK01008296.1:1-1329 GIWK01008296.1:1454-2795 GIWK01008296.1:844-1964 GIWK01064060.1:1-1464 GIWK01064060.1:2-1462 GIWK01064060.1:3-1464 GIWK01064060.1:7-1381 GIWK01064060.1:7-1466 GIWK01064060.1:302-1467 GIWK01064060.1:2-1467 GIWK01064060.1:2-1464 GIWK01064060.1:5-1464 GIWK01064060.1:1-1364 GIWK01064060.1:4-1196 GIWK01064060.1:1-1277 GIWK01064060.1:4-1464 GIWK01064060.1:2-1445 GIWK01064060.1:109-1466 GIWK01064060.1:67-1466 GIWK01064060.1:235-1464 GIWK01064060.1:2-1322 GIWK01064060.1:169-1467 GIWK01064060.1:193-1466 GIWK01064060.1:5-1238 GIWK01064060.1:260-1467 GIWK01064060.1:403-1466 GIWK01064060.1:5-1112 GIWK01064060.1:463-1459 GIWK01064060.1:340-1459 GIWK01064060.1:5-1028 GIWK01064060.1:1-1067 GIWK01064060.1:466-1462 GIWK01064060.1:544-1459 GIWK01064060.1:547-1466 GIWK01064060.1:5-902 GIWK01064060.1:1-944 GIWK01064060.1:652-1466 GIWK01064060.1:1-860 GIWK01064060.1:610-1467 GIWK01064060.1:706-1466 GIWK01064060.1:736-1467 GIWK01064060.1:7-692 GIWK01064060.1:2-776 GIWK01064060.1:778-1466 GIWK01064060.1:803-1467 GIWK01064060.1:845-1467 GIWK01064060.1:5-566 GIWK01064060.1:2-502 GIWK01064060.1:971-1466 GIWK01052548.1:1413-4178 GIWK01052548.1:3015-4178 GIWK01052548.1:3022-4178 GIWK01052548.1:3015-4174 GIWK01052548.1:3015-4178 GIWK01052548.1:3057-4178 GIWK01052548.1:3015-4178 GIWK01052548.1:3015-4178 GIWK01052548.1:3017-4178 GIWK01052548.1:3015-4178 GIWK01052548.1:3015-4178 GIWK01052548.1:3099-4178 GIWK01052548.1:3015-4159 GIWK01052548.1:638-2047 GIWK01052548.1:3015-4178 GIWK01052548.1:3015-4165 GIWK01052548.1:3148-4178 GIWK01052548.1:3015-4077 GIWK01052548.1:3183-4178 GIWK01052548.1:3022-4102 GIWK01052548.1:1-1116 GIWK01052548.1:3015-4039 GIWK01052548.1:3225-4178 GIWK01052548.1:3017-3993 GIWK01052548.1:3267-4178 GIWK01052548.1:3015-3934 GIWK01052548.1:3351-4165 GIWK01052548.1:3438-4165 GIWK01052548.1:3393-4178 GIWK01052548.1:3015-3892 GIWK01052548.1:3015-3867 GIWK01052548.1:3015-3706 GIWK01052548.1:3477-4178 GIWK01052548.1:3519-4163 GIWK01052548.1:3017-3787 GIWK01052548.1:3017-3703 GIWK01052548.1:3649-4178 GIWK01052548.1:3015-3622 GIWK01052548.1:3684-4178 GIWK01052548.1:3015-3559 GIWK01038917.1:1-2544 GIWK01038917.1:1-944 GIWK01038917.1:1910-3319 GIWK01038917.1:1-944 GIWK01038917.1:1-937 GIWK01038917.1:2-944 GIWK01038917.1:5-902 GIWK01038917.1:2-944 GIWK01038917.1:1-942 GIWK01038917.1:7-944 GIWK01038917.1:5-944 GIWK01038917.1:5-944 GIWK01038917.1:2-944 GIWK01038917.1:5-944 GIWK01038917.1:7-944 GIWK01038917.1:2-942 GIWK01038917.1:5-944 GIWK01038917.1:10-919 GIWK01038917.1:25-944 GIWK01038917.1:5-944 GIWK01038917.1:1-860 GIWK01038917.1:2-811 GIWK01038917.1:67-944 GIWK01038917.1:92-944 GIWK01038917.1:253-944 GIWK01038917.1:2-725 GIWK01038917.1:7-692 GIWK01038917.1:172-942 GIWK01038917.1:5-608 GIWK01038917.1:256-942 GIWK01038917.1:5-566 GIWK01038917.1:2-521 GIWK01038917.1:337-944 GIWK01038917.1:7-482 GIWK01038917.1:400-944 GIWK01038917.1:2841-3510 GIWK01038917.1:3496-3748 GHVE01065598.1:1-744 GHVE01065598.1:1-744 GHVE01065598.1:2647-3537 GHVE01065598.1:1446-2270 GHVE01065598.1:1739-2729 GIWK01026002.1:556-1857 GIWK01026002.1:6-1034 GIWK01000702.1:428-1719 GIWK01000702.1:8-904 GIWK01032289.1:320-1615 GIWK01032289.1:1-798 GIWK01043700.1:525-1818 GIWK01043700.1:1-1003 GIWK01065725.1:1-1685 GIWK01065725.1:1-1668 GIWK01065725.1:1-1604 GIWK01065725.1:2-1545 GIWK01065725.1:8-1697 GIWK01065725.1:80-1687 GIWK01065725.1:56-1687 GIWK01065725.1:1-1587 GIWK01065725.1:140-1697 GIWK01065725.1:164-1689 GIWK01065725.1:539-1697 GIWK01065725.1:1-1500 GIWK01065725.1:248-1697 GIWK01065725.1:1-1419 GIWK01065725.1:347-1697 GIWK01065725.1:455-1697 GIWK01065725.1:1-1377 GIWK01065725.1:290-1697 GIWK01065725.1:350-1697 GIWK01065725.1:413-1689 GIWK01065725.1:606-1689 GIWK01065725.1:1-1209 GIWK01065725.1:1-992 GIWK01065725.1:1-1167 GIWK01065725.1:564-1682 GIWK01065725.1:1-1125 GIWK01065725.1:1-1083 GIWK01065725.1:644-1682 GIWK01065725.1:707-1685 GIWK01065725.1:749-1682 GIWK01065725.1:1-1041 GIWK01065725.1:1-873 GIWK01065725.1:809-1697 GIWK01065725.1:1-906 GIWK01065725.1:833-1697 GIWK01065725.1:1-789 GIWK01065725.1:893-1697 GIWK01065725.1:917-1697 GIWK01065725.1:959-1697 GIWK01065725.1:1001-1697 GIWK01065725.1:20-822 GIWK01065725.1:1022-1697 GIWK01065725.1:1068-1697 GIWK01065725.1:20-660 GIWK01065725.1:1-621 GIWK01065725.1:1194-1697 GIWK01063059.1:110-1400 GIWK01063059.1:1-586 GIWK01012354.1:1073-2370 GIWK01012354.1:142-1551 GIWK01017897.1:1299-2594 GIWK01017897.1:368-1773 GIWK01017897.1:1-1002 GIWK01028724.1:1-879 GIWK01028724.1:525-1990 GIWK01039335.1:2025-3316 GIWK01039335.1:658-2503 GIWK01039335.1:1-660 GIWK01039335.1:1-660 GIWK01039335.1:1-660 GIWK01039335.1:1-660 GIWK01039335.1:19-659 GIWK01039335.1:1-661 GIWK01039335.1:1-660 GIWK01039335.1:7-661 GIWK01039335.1:1-660 GIWK01039335.1:1-660 GIWK01039335.1:1-660 GIWK01039335.1:19-661 GIWK01039335.1:1-661 GIWK01039335.1:1-660 GIWK01039335.1:1-660 GIWK01039335.1:1-660 GIWK01039335.1:1-660 GIWK01039335.1:1-660 GIWK01039335.1:1-661 GIWK01039335.1:1-660 GIWK01039335.1:1-660 GIWK01039335.1:1-660 GIWK01039335.1:37-660 GIWK01039335.1:1-581 GIWK01039335.1:62-660 GIWK01039335.1:121-660 GIWK01039335.1:1-497 GIWK01039335.1:146-660 GIWK01039335.1:188-660 GIWK01029994.1:878-2146 GIWK01067018.1:1-1423 GIWK01067018.1:5-1423 GIWK01067018.1:275-1427 GIWK01067018.1:1-1427 GIWK01067018.1:3-1423 GIWK01067018.1:1-1421 GIWK01067018.1:1-1423 GIWK01067018.1:3-1340 GIWK01067018.1:1-1425 GIWK01067018.1:5-1404 GIWK01067018.1:65-1425 GIWK01067018.1:3-1281 GIWK01067018.1:23-1425 GIWK01067018.1:107-1427 GIWK01067018.1:1-1323 GIWK01067018.1:191-1423 GIWK01067018.1:2-1197 GIWK01067018.1:3-1236 GIWK01067018.1:167-1425 GIWK01067018.1:251-1427 GIWK01067018.1:342-1425 GIWK01067018.1:317-1418 GIWK01067018.1:3-1155 GIWK01067018.1:3-1113 GIWK01067018.1:1-1071 GIWK01067018.1:401-1418 GIWK01067018.1:3-987 GIWK01067018.1:3-1026 GIWK01067018.1:485-1418 GIWK01067018.1:3-903 GIWK01067018.1:545-1425 GIWK01067018.1:636-1425 GIWK01067018.1:1-861 GIWK01067018.1:674-1427 GIWK01067018.1:737-1425 GIWK01067018.1:779-1427 GIWK01067018.1:3-777 GIWK01067018.1:1-735 GIWK01067018.1:804-1427 GIWK01067018.1:1-651 GIWK01067018.1:1-525 GIWK01067018.1:5-474 GIWK01067018.1:947-1425 GIWK01061984.1:1-1659 GIWK01061984.1:1-1625 GIWK01061984.1:645-1659 GIWK01061984.1:118-1645 GIWK01061984.1:1-1645 GIWK01061984.1:1-1586 GIWK01061984.1:202-1659 GIWK01061984.1:1-1502 GIWK01061984.1:1-1415 GIWK01061984.1:205-1659 GIWK01061984.1:2-1376 GIWK01061984.1:286-1659 GIWK01061984.1:283-1659 GIWK01061984.1:391-1645 GIWK01061984.1:1-1250 GIWK01061984.1:445-1659 GIWK01061984.1:517-1642 GIWK01061984.1:1-1208 GIWK01061984.1:1-946 GIWK01061984.1:475-1659 GIWK01061984.1:647-1645 GIWK01061984.1:1-1166 GIWK01061984.1:1-1124 GIWK01061984.1:685-1642 GIWK01061984.1:1-1015 GIWK01061984.1:731-1659 GIWK01061984.1:790-1659 GIWK01061984.1:1-1082 GIWK01061984.1:815-1659 GIWK01061984.1:1-995 GIWK01061984.1:886-1659 GIWK01061984.1:1-830 GIWK01061984.1:958-1659 GIWK01061984.1:1-655 GIWK01061984.1:979-1659 GIWK01061984.1:1-788 GIWK01061984.1:1042-1659 GIWK01061984.1:1-641 GIWK01061984.1:1067-1659 GIWK01061984.1:1126-1659 GIWK01061984.1:1-550 GIWK01061984.1:1151-1645 GIWK01061984.1:1-515 GIWK01061984.1:1193-1659 GIWK01032646.1:936-2223 GIWK01032646.1:7-1410 GIWK01067527.1:652-1939 GIWK01067527.1:1-1128 GHVE01065605.1:1-811 GHVE01065605.1:1-811 GHVE01065605.1:2714-3550 GHVE01065605.1:1513-2337 GHVE01065605.1:1806-2796 GHVE01065593.1:69-1053 GHVE01065593.1:134-1053 GHVE01065593.1:2956-3846 GHVE01065593.1:1755-2579 GHVE01065593.1:2048-3038 GIWK01053399.1:1-2656 GIWK01053399.1:2146-3236 GIWK01053399.1:1-421 GIWK01053399.1:1-423 GIWK01053399.1:1-423 GIWK01053399.1:1-421 GIWK01053399.1:2-423 GIWK01053399.1:1-423 GIWK01053399.1:1-423 GIWK01053399.1:1-423 GIWK01053399.1:1-421 GIWK01053399.1:1-423 GIWK01053399.1:1-423 GIWK01053399.1:5-423 GHVE01065589.1:1-729 GHVE01065589.1:1-729 GHVE01065589.1:2632-3522 GHVE01065589.1:1431-2255 GHVE01065589.1:1724-2714 GHVE01065596.1:1-923 GHVE01065596.1:8-923 GHVE01065596.1:2826-3716 GHVE01065596.1:1625-2449 GHVE01065596.1:1918-2908 GIWK01065045.1:661-2013 GIWK01065045.1:6-1171 GIWK01033542.1:806-2151 GIWK01033542.1:3-1316 GIWK01000290.1:277-1623 GIWK01000290.1:1-787 GIWK01000327.1:398-1744 GIWK01000327.1:1-908 GIWK01050953.1:1048-2394 GIWK01050953.1:144-1558 GIWK01056241.1:607-1954 GIWK01056241.1:2-1117 GIWK01004636.1:773-2116 GIWK01004636.1:135-1283 GIWK01004636.1:1-649 GIWK01001255.1:71-1410 GIWK01001255.1:1-581 GIWK01042089.1:1627-3093 GIWK01042089.1:33-1330 GIWK01042089.1:856-2259 GKQY01000390.1:2666-3887 GKQY01000390.1:389-1646 GKQY01000390.1:3044-3887 GKQY01000390.1:164-341 GKQY01000393.1:389-3431 GKQY01000393.1:2369-3590 GKQY01000393.1:2747-3590 GKQY01000393.1:164-341 GHVE01065604.1:100-1064 GHVE01065604.1:184-1066 GHVE01065604.1:2957-3793 GHVE01065604.1:1756-2580 GHVE01065604.1:2049-3039 GKQY01000391.1:389-3431 GKQY01000391.1:2411-3632 GKQY01000391.1:2789-3632 GKQY01000391.1:164-341 GHVE01065586.1:92-1062 GHVE01065586.1:175-1064 GHVE01065586.1:2955-3937 GHVE01065586.1:1754-2578 GHVE01065586.1:2047-3037 GKQY01000389.1:389-3448 GKQY01000389.1:2470-3691 GKQY01000389.1:2848-3691 GKQY01000389.1:164-341 GIWK01050585.1:1-3083 GIWK01050585.1:2573-3669 GIWK01050585.1:1-726 GIWK01050585.1:1-595 GIWK01050585.1:1-728 GIWK01050585.1:1-726 GIWK01050585.1:1-728 GIWK01050585.1:1-728 GIWK01050585.1:1-728 GIWK01050585.1:1-728 GIWK01050585.1:1-726 GIWK01050585.1:1-728 GIWK01050585.1:1-728 GIWK01050585.1:1-728 GIWK01050585.1:1-728 GIWK01050585.1:1-728 GIWK01050585.1:1-728 GIWK01050585.1:1-728 GIWK01050585.1:1-686 GIWK01050585.1:1-728 GIWK01050585.1:1-644 GIWK01050585.1:142-728 GIWK01050585.1:1-509 GIWK01050585.1:1-476 GIWK01050585.1:167-728 GIWK01050585.1:310-728 GHVE01065599.1:149-1037 GHVE01065599.1:149-1037 GHVE01065599.1:2940-3830 GHVE01065599.1:1739-2563 GHVE01065599.1:2032-3022 GIWK01046963.1:1-3057 GIWK01046963.1:2547-3892 GIWK01046963.1:1-701 GIWK01046963.1:13-703 GIWK01046963.1:1-570 GIWK01046963.1:1-703 GIWK01046963.1:1-703 GIWK01046963.1:1-701 GIWK01046963.1:1-703 GIWK01046963.1:1-701 GIWK01046963.1:1-703 GIWK01046963.1:1-703 GIWK01046963.1:1-703 GIWK01046963.1:1-703 GIWK01046963.1:14-661 GIWK01046963.1:1-703 GIWK01046963.1:1-703 GIWK01046963.1:1-703 GIWK01046963.1:1-703 GIWK01046963.1:1-703 GIWK01046963.1:117-703 GIWK01046963.1:1-484 GIWK01046963.1:142-703 GIWK01046963.1:285-703 GIWK01046137.1:1-2930 GIWK01046137.1:2420-3515 GIWK01046137.1:1-578 GIWK01046137.1:1-578 GIWK01046137.1:1-576 GIWK01046137.1:1-576 GIWK01046137.1:1-578 GIWK01046137.1:1-578 GIWK01046137.1:1-578 GIWK01046137.1:18-578 GIWK01046137.1:1-578 GIWK01046137.1:1-568 GIWK01046137.1:55-578 GIWK01046137.1:17-578 GIWK01046137.1:17-578 GIWK01046137.1:1-578 GIWK01046137.1:1-536 GIWK01046137.1:1-578 GIWK01046137.1:1-494 GIWK01046137.1:139-578 GIWK01022639.1:1-1375 GIWK01022639.1:743-2150 GIWK01022639.1:1672-2466 GHVE01065606.1:50-1043 GHVE01065606.1:134-1043 GHVE01065606.1:2946-3698 GHVE01065606.1:1745-2569 GHVE01065606.1:2038-3028 GIWK01000847.1:2-1622 GIWK01039616.1:1-2617 GIWK01039616.1:2107-3457 GIWK01039616.1:36-262 GIWK01039616.1:1-262 GIWK01039616.1:1-260 GIWK01039616.1:1-260 GIWK01039616.1:2-262 GIWK01039616.1:2-262 GIWK01039616.1:1-262 GIWK01039616.1:3-262 GIWK01039616.1:2-262 GHVE01065620.1:631-1903 GHVE01065620.1:406-583 GIWK01016097.1:763-1976 GIWK01016097.1:1500-2180 GKQY01000392.1:389-2078 GKQY01000392.1:164-341 GIWK01005258.1:76-1483 GIWK01005258.1:1007-2053 GIWK01037455.1:2114-3221 GIWK01037455.1:1205-2622 GHVE01065581.1:5351-6898 GHVE01065581.1:889-2161 GHVE01065581.1:2466-3448 GHVE01065581.1:2567-3448 GHVE01065581.1:4150-4974 GHVE01065581.1:4443-5433 GHVE01065581.1:664-841 GHVE01065580.1:4988-6535 GHVE01065580.1:844-2116 GHVE01065580.1:2298-3085 GHVE01065580.1:2354-3085 GHVE01065580.1:3787-4611 GHVE01065580.1:4080-5070 GHVE01065580.1:619-796 GKQY01000394.1:389-1868 GKQY01000394.1:164-341 GHVE01065579.1:4590-6214 GHVE01065579.1:749-2021 GHVE01065579.1:3389-4213 GHVE01065579.1:3682-4672 GHVE01065579.1:2220-2687 GHVE01065579.1:2203-2687 GHVE01065579.1:524-701 GHVE01065578.1:3066-4690 GHVE01065578.1:224-1173 GHVE01065578.1:325-1175 GHVE01065578.1:1865-2689 GHVE01065578.1:2158-3148 GIWK01036431.1:818-3149 GIWK01036431.1:31-1172 GIWK01047599.1:1-1091 GIWK01047599.1:578-1373 GIWK01061608.1:286-1353 GIWK01061608.1:1-796 GIWK01061608.1:1353-1502 GIWK01049141.1:547-2485 GIWK01049141.1:1975-3314 GIWK01049141.1:4-546 GIWK01049141.1:4-538 GIWK01049141.1:4-543 GIWK01049141.1:1-475 GIWK01049141.1:4-549 GIWK01049141.1:1-546 GIWK01049141.1:1-549 GIWK01049141.1:4-543 GIWK01049141.1:4-538 GIWK01049141.1:4-549 GIWK01049141.1:4-543 GIWK01049141.1:7-549 GIWK01049141.1:4-543 GIWK01049141.1:4-549 GIWK01049141.1:4-549 GIWK01049141.1:4-546 GIWK01049141.1:22-543 GIWK01049141.1:47-543 GIWK01039343.1:324-2257 GIWK01039343.1:1747-3088 GIWK01039343.1:1-324 GHVE01065584.1:2674-3671 GHVE01065584.1:69-771 GHVE01065584.1:39-771 GHVE01065584.1:1473-2297 GHVE01065584.1:1766-2756 GIWK01009859.1:35-1496 GIWK01009859.1:1145-2027

#### TSA ids matching Lampetra_fluvialis_SELENOP_nt, indicated with their respective matched position range:

GKQY01000391.1:164-3431/1-3268 GKQY01000393.1:164-3431/1-3268 GKQY01000389.1:164-3448/1-3285 GIWK01065380.1:1308-1622/1-315 GIWK01015113.1:1222-1912/1-691 GIWK01015113.1:1306-1912/1-607 GIWK01015113.1:1327-1912/1-586 GIWK01038917.1:7-566/1-560 GIWK01050416.1:1-585/1-585 GIWK01050416.1:156-770/1-615 GIWK01050416.1:215-770/1-556 GIWK01050416.1:1-504/1-504 GIWK01053646.1:1961-3156/1-1196 GIWK01053646.1:1961-3152/1-1192 GIWK01053646.1:1961-3156/1-1196 GIWK01053646.1:1961-3149/1-1189 GIWK01053646.1:1963-3149/1-1187 GIWK01053646.1:1961-3154/1-1194 GIWK01053646.1:1961-3149/1-1189 GIWK01053646.1:1961-3093/1-1133 GIWK01053646.1:1961-3149/1-1189 GIWK01053646.1:1961-3152/1-1192 GIWK01053646.1:1964-3156/1-1193 GIWK01053646.1:1989-3156/1-1168 GIWK01053646.1:1961-3154/1-1194 GIWK01053646.1:2090-3156/1-1067 GIWK01053646.1:1961-3135/1-1175 GIWK01053646.1:2060-3149/1-1090 GIWK01053646.1:1961-2999/1-1039 GIWK01053646.1:2153-3154/1-1002 GIWK01053646.1:1961-2903/1-943 GIWK01053646.1:1961-2967/1-1007 GIWK01053646.1:2216-3156/1-941 GIWK01053646.1:2258-3156/1-899 GIWK01053646.1:1961-2799/1-839 GIWK01053646.1:2300-3154/1-855 GIWK01053646.1:1961-2760/1-800 GIWK01053646.1:2367-3149/1-783 GIWK01053646.1:1961-2718/1-758 GIWK01053646.1:2405-3156/1-752 GIWK01053646.1:1961-2589/1-629 GIWK01053646.1:2552-3156/1-605 GIWK01053646.1:1961-2466/1-506 GIWK01053646.1:1961-2508/1-548 GIWK01053646.1:2577-3156/1-580 GIWK01053646.1:2678-3156/1-479 GIWK01053646.1:1961-2424/1-464 GIWK01053646.1:1961-2382/1-422 GIWK01053646.1:1961-2340/1-380 GIWK01053646.1:1961-2298/1-338 GIWK01053646.1:2864-3156/1-293 GIWK01053646.1:2867-3156/1-290 GIWK01053646.1:1961-2088/1-128 GIWK01065380.1:163-1297/1-1135 GIWK01065380.1:1473-1619/1-147 GIWK01065380.1:1473-1619/1-147 GIWK01065380.1:1453-1611/1-159 GIWK01065380.1:1456-1614/1-159 GIWK01065380.1:1453-1616/1-164 GIWK01065380.1:1473-1619/1-147 GIWK01015113.1:992-1766/1-775 GIWK01015113.1:985-1721/1-737 GIWK01015113.1:1180-1910/1-731 GIWK01015113.1:992-1640/1-649 GIWK01015113.1:985-1682/1-698 GIWK01015113.1:992-1511/1-520 GIWK01015113.1:992-1388/1-397 GIWK01015113.1:985-1430/1-446 GIWK01015113.1:1474-1910/1-437 GIWK01015113.1:992-1304/1-313 GIWK01015113.1:994-1262/1-269 GIWK01015113.1:992-1220/1-229 GIWK01015113.1:985-1116/1-132 GIWK01015113.1:985-1084/1-100 GIWK01011755.1:3-342/1-340 GIWK01011755.1:3-213/1-211 GIWK01011755.1:3-174/1-172 GIWK01011755.1:3-132/1-130 GIWK01050585.1:1-1011/1-1011 GIWK01050585.1:1717-3138/1-1422 GIWK01050585.1:1170-2302/1-1133 GIWK01050585.1:2573-3669/1-1097 GIWK01050585.1:1-728/1-728 GIWK01050585.1:1-728/1-728 GIWK01050585.1:1-728/1-728 GIWK01050585.1:1-728/1-728 GIWK01050585.1:1-728/1-728 GIWK01050585.1:2-728/1-727 GIWK01050585.1:2-726/1-725 GIWK01050585.1:1-728/1-728 GIWK01050585.1:1-728/1-728 GIWK01050585.1:1-728/1-728 GIWK01050585.1:1-728/1-728 GIWK01050585.1:1-728/1-728 GIWK01050585.1:1-728/1-728 GIWK01050585.1:1-728/1-728 GIWK01050585.1:16-728/1-713 GIWK01050585.1:1-624/1-624 GIWK01050585.1:58-728/1-671 GIWK01050585.1:112-728/1-617 GIWK01050585.1:1-490/1-490 GIWK01050585.1:1-392/1-392 GIWK01050585.1:209-728/1-520 GIWK01050585.1:1-424/1-424 GIWK01050585.1:1-308/1-308 GIWK01050585.1:412-718/1-307 GIWK01050585.1:1-224/1-224 GIWK01050585.1:1-140/1-140 GIWK01067313.1:1-1359/1-1359 GIWK01067313.1:1-1348/1-1348 GIWK01067313.1:1-1348/1-1348 GIWK01067313.1:1-1353/1-1353 GIWK01067313.1:1-1334/1-1334 GIWK01067313.1:7-1348/1-1342 GIWK01067313.1:1-1355/1-1355 GIWK01067313.1:1-1351/1-1351 GIWK01067313.1:1-1292/1-1292 GIWK01067313.1:1-1359/1-1359 GIWK01067313.1:37-1353/1-1317 GIWK01067313.1:1-1250/1-1250 GIWK01067313.1:121-1348/1-1228 GIWK01067313.1:79-1353/1-1275 GIWK01067313.1:1-1208/1-1208 GIWK01067313.1:146-1353/1-1208 GIWK01067313.1:1-1121/1-1121 GIWK01067313.1:1-1141/1-1141 GIWK01067313.1:188-1353/1-1166 GIWK01067313.1:289-1353/1-1065 GIWK01067313.1:226-1348/1-1123 GIWK01067313.1:1-995/1-995 GIWK01067313.1:1-1040/1-1040 GIWK01067313.1:1-914/1-914 GIWK01067313.1:373-1355/1-983 GIWK01067313.1:1-956/1-956 GIWK01067313.1:398-1353/1-956 GIWK01067313.1:1-820/1-820 GIWK01067313.1:499-1353/1-855 GIWK01067313.1:1-788/1-788 GIWK01067313.1:601-1348/1-748 GIWK01067313.1:1-724/1-724 GIWK01067313.1:688-1353/1-666 GIWK01067313.1:685-1353/1-669 GIWK01067313.1:1-620/1-620 GIWK01067313.1:13-539/1-527 GIWK01067313.1:1-581/1-581 GIWK01067313.1:790-1353/1-564 GIWK01067313.1:832-1353/1-522 GIWK01067313.1:874-1353/1-480 GIWK01067313.1:1-410/1-410 GIWK01067313.1:1-329/1-329 GIWK01067313.1:1-287/1-287 GIWK01067313.1:1-245/1-245 GIWK01067313.1:2-203/1-202 GIWK01067313.1:1-161/1-161 GIWK01067313.1:2-119/1-118 GIWK01067313.1:1-77/1-77 GIWK01059119.1:191-1715/1-1525 GIWK01059119.1:1566-1712/1-147 GIWK01059119.1:1566-1712/1-147 GIWK01059119.1:1546-1704/1-159 GIWK01059119.1:1549-1707/1-159 GIWK01059119.1:1546-1709/1-164 GIWK01059119.1:1566-1712/1-147 GIWK01059119.1:163-198/1-36 GIWK01020752.1:1-1435/1-1435 GIWK01020752.1:1-1440/1-1440 GIWK01020752.1:1-1440/1-1440 GIWK01020752.1:1-1438/1-1438 GIWK01020752.1:1-1438/1-1438 GIWK01020752.1:1-1440/1-1440 GIWK01020752.1:8-1435/1-1428 GIWK01020752.1:1-1369/1-1369 GIWK01020752.1:1-1421/1-1421 GIWK01020752.1:20-1440/1-1421 GIWK01020752.1:1-1440/1-1440 GIWK01020752.1:1-1253/1-1253 GIWK01020752.1:1-1267/1-1267 GIWK01020752.1:79-1435/1-1357 GIWK01020752.1:146-1440/1-1295 GIWK01020752.1:121-1440/1-1320 GIWK01020752.1:1-1169/1-1169 GIWK01020752.1:1-1211/1-1211 GIWK01020752.1:307-1440/1-1134 GIWK01020752.1:226-1440/1-1215 GIWK01020752.1:1-1127/1-1127 GIWK01020752.1:1-1082/1-1082 GIWK01020752.1:310-1435/1-1126 GIWK01020752.1:373-1440/1-1068 GIWK01020752.1:1-1043/1-1043 GIWK01020752.1:1-959/1-959 GIWK01020752.1:440-1440/1-1001 GIWK01020752.1:17-1001/1-985 GIWK01020752.1:478-1440/1-963 GIWK01020752.1:1-917/1-917 GIWK01020752.1:1-872/1-872 GIWK01020752.1:541-1440/1-900 GIWK01020752.1:1-833/1-833 GIWK01020752.1:566-1440/1-875 GIWK01020752.1:625-1440/1-816 GIWK01020752.1:1-791/1-791 GIWK01020752.1:650-1435/1-786 GIWK01020752.1:709-1440/1-732 GIWK01020752.1:751-1440/1-690 GIWK01020752.1:17-539/1-523 GIWK01020752.1:1-578/1-578 GIWK01020752.1:856-1440/1-585 GIWK01020752.1:853-1440/1-588 GIWK01020752.1:1-497/1-497 GIWK01020752.1:17-413/1-397 GIWK01020752.1:17-242/1-226 GIWK01020752.1:2-203/1-202 GIWK01020752.1:1171-1440/1-270 GIWK01020752.1:1-161/1-161 GIWK01062408.1:70-1521/1-1452 GIWK01062408.1:2-1520/1-1519 GIWK01062408.1:4-1501/1-1498 GIWK01062408.1:2-1521/1-1520 GIWK01062408.1:64-1521/1-1458 GIWK01062408.1:2-1462/1-1461 GIWK01062408.1:2-1358/1-1357 GIWK01062408.1:2-1445/1-1444 GIWK01062408.1:106-1521/1-1416 GIWK01062408.1:46-1521/1-1476 GIWK01062408.1:131-1521/1-1391 GIWK01062408.1:2-1516/1-1515 GIWK01062408.1:190-1521/1-1332 GIWK01062408.1:1-1403/1-1403 GIWK01062408.1:215-1521/1-1307 GIWK01062408.1:316-1521/1-1206 GIWK01062408.1:4-1319/1-1316 GIWK01062408.1:2-1235/1-1234 GIWK01062408.1:257-1521/1-1265 GIWK01062408.1:2-1277/1-1276 GIWK01062408.1:376-1521/1-1146 GIWK01062408.1:379-1521/1-1143 GIWK01062408.1:457-1521/1-1065 GIWK01062408.1:2-1109/1-1108 GIWK01062408.1:2-1025/1-1024 GIWK01062408.1:2-1064/1-1063 GIWK01062408.1:2-983/1-982 GIWK01062408.1:565-1524/1-960 GIWK01062408.1:523-1521/1-999 GIWK01062408.1:607-1521/1-915 GIWK01062408.1:2-899/1-898 GIWK01062408.1:1-773/1-773 GIWK01062408.1:2-829/1-828 GIWK01062408.1:2-689/1-688 GIWK01062408.1:2-815/1-814 GIWK01062408.1:775-1521/1-747 GIWK01062408.1:733-1520/1-788 GIWK01062408.1:835-1521/1-687 GIWK01062408.1:859-1521/1-663 GIWK01062408.1:2-644/1-643 GIWK01062408.1:2-605/1-604 GIWK01062408.1:926-1521/1-596 GIWK01062408.1:964-1521/1-558 GIWK01062408.1:2-479/1-478 GIWK01062408.1:2-356/1-355 GIWK01062408.1:1111-1521/1-411 GIWK01062408.1:4-272/1-269 GIWK01062408.1:2-220/1-219 GIWK01062408.1:2-188/1-187 GIWK01068541.1:1-1271/1-1271 GIWK01068541.1:1-1260/1-1260 GIWK01068541.1:1-1265/1-1265 GIWK01068541.1:12-1267/1-1256 GIWK01068541.1:12-1265/1-1254 GIWK01068541.1:1-1260/1-1260 GIWK01068541.1:12-1246/1-1235 GIWK01068541.1:1-1265/1-1265 GIWK01068541.1:1-1265/1-1265 GIWK01068541.1:12-1263/1-1252 GIWK01068541.1:12-1260/1-1249 GIWK01068541.1:1-1265/1-1265 GIWK01068541.1:12-1271/1-1260 GIWK01068541.1:12-1162/1-1151 GIWK01068541.1:13-1260/1-1248 GIWK01068541.1:55-1265/1-1211 GIWK01068541.1:14-1204/1-1191 GIWK01068541.1:114-1265/1-1152 GIWK01068541.1:210-1265/1-1056 GIWK01068541.1:12-1053/1-1042 GIWK01068541.1:12-984/1-973 GIWK01068541.1:240-1265/1-1026 GIWK01068541.1:12-1033/1-1022 GIWK01068541.1:282-1267/1-986 GIWK01068541.1:324-1265/1-942 GIWK01068541.1:12-952/1-941 GIWK01068541.1:366-1265/1-900 GIWK01068541.1:12-865/1-854 GIWK01068541.1:12-826/1-815 GIWK01068541.1:433-1265/1-833 GIWK01068541.1:1-784/1-784 GIWK01068541.1:471-1260/1-790 GIWK01068541.1:618-1265/1-648 GIWK01068541.1:12-655/1-644 GIWK01068541.1:1-574/1-574 GIWK01068541.1:12-532/1-521 GIWK01068541.1:702-1265/1-564 GIWK01068541.1:12-490/1-479 GIWK01068541.1:769-1265/1-497 GIWK01068541.1:12-448/1-437 GIWK01068541.1:12-406/1-395 GIWK01068541.1:12-364/1-353 GIWK01068541.1:12-322/1-311 GIWK01068541.1:12-238/1-227 GIWK01068541.1:1-154/1-154 GIWK01052548.1:2739-4178/1-1440 GIWK01052548.1:3015-4178/1-1164 GIWK01052548.1:3015-4178/1-1164 GIWK01052548.1:3015-4178/1-1164 GIWK01052548.1:3015-4178/1-1164 GIWK01052548.1:3057-4178/1-1122 GIWK01052548.1:3015-4178/1-1164 GIWK01052548.1:3015-4178/1-1164 GIWK01052548.1:3015-4178/1-1164 GIWK01052548.1:3015-4138/1-1124 GIWK01052548.1:3015-4178/1-1164 GIWK01052548.1:3015-4178/1-1164 GIWK01052548.1:3016-4174/1-1159 GIWK01052548.1:3099-4178/1-1080 GIWK01052548.1:3015-4165/1-1151 GIWK01052548.1:1-1116/1-1116 GIWK01052548.1:3148-4178/1-1031 GIWK01052548.1:3183-4165/1-983 GIWK01052548.1:3015-4018/1-1004 GIWK01052548.1:3235-4178/1-944 GIWK01052548.1:3267-4178/1-912 GIWK01052548.1:1423-2577/1-1155 GIWK01052548.1:3393-4178/1-786 GIWK01052548.1:604-2077/1-1474 GIWK01052548.1:3309-4178/1-870 GIWK01052548.1:3015-3993/1-979 GIWK01052548.1:3015-3934/1-920 GIWK01052548.1:3015-3790/1-776 GIWK01052548.1:3477-4178/1-702 GIWK01052548.1:3015-3787/1-773 GIWK01052548.1:3017-3867/1-851 GIWK01052548.1:3438-4178/1-741 GIWK01052548.1:3015-3706/1-692 GIWK01052548.1:3015-3703/1-689 GIWK01052548.1:3015-3625/1-611 GIWK01052548.1:3603-4162/1-560 GIWK01052548.1:3771-4165/1-395 GIWK01052548.1:3810-4178/1-369 GIWK01052548.1:3852-4165/1-314 GIWK01052548.1:3016-3349/1-334 GIWK01052548.1:4065-4178/1-114 GIWK01059296.1:7-1521/1-1515 GIWK01059296.1:5-1521/1-1517 GIWK01059296.1:2-1521/1-1520 GIWK01059296.1:2-1504/1-1503 GIWK01059296.1:5-1521/1-1517 GIWK01059296.1:5-1490/1-1486 GIWK01059296.1:31-1521/1-1491 GIWK01059296.1:25-1521/1-1497 GIWK01059296.1:109-1521/1-1413 GIWK01059296.1:2-1361/1-1360 GIWK01059296.1:50-1504/1-1455 GIWK01059296.1:1-1448/1-1448 GIWK01059296.1:2-1381/1-1380 GIWK01059296.1:7-1312/1-1306 GIWK01059296.1:211-1521/1-1311 GIWK01059296.1:1-1280/1-1280 GIWK01059296.1:214-1521/1-1308 GIWK01059296.1:5-1196/1-1192 GIWK01059296.1:298-1521/1-1224 GIWK01059296.1:376-1521/1-1146 GIWK01059296.1:295-1504/1-1210 GIWK01059296.1:1-1235/1-1235 GIWK01059296.1:5-1154/1-1150 GIWK01059296.1:400-1504/1-1105 GIWK01059296.1:442-1521/1-1080 GIWK01059296.1:1-1028/1-1028 GIWK01059296.1:484-1521/1-1038 GIWK01059296.1:5-1067/1-1063 GIWK01059296.1:538-1521/1-984 GIWK01059296.1:5-944/1-940 GIWK01059296.1:2-986/1-985 GIWK01059296.1:5-902/1-898 GIWK01059296.1:635-1521/1-887 GIWK01059296.1:673-1521/1-849 GIWK01059296.1:1-818/1-818 GIWK01059296.1:5-860/1-856 GIWK01059296.1:2-776/1-775 GIWK01059296.1:778-1521/1-744 GIWK01059296.1:5-734/1-730 GIWK01059296.1:803-1521/1-719 GIWK01059296.1:946-1515/1-570 GIWK01059296.1:2-650/1-649 GIWK01059296.1:7-566/1-560 GIWK01059296.1:1030-1512/1-483 GIWK01059296.1:2-398/1-397 GIWK01059296.1:2-230/1-229 GIWK01059296.1:2-191/1-190 GIWK01059296.1:5-149/1-145 GIWK01059296.1:5-107/1-103 GIWK01056333.1:1-1182/1-1182 GIWK01056333.1:1-1178/1-1178 GIWK01056333.1:1-1183/1-1183 GIWK01056333.1:1-1175/1-1175 GIWK01056333.1:1-1180/1-1180 GIWK01056333.1:1-1182/1-1182 GIWK01056333.1:1-1175/1-1175 GIWK01056333.1:1-1180/1-1180 GIWK01056333.1:1-1183/1-1183 GIWK01056333.1:1-1183/1-1183 GIWK01056333.1:1-1175/1-1175 GIWK01056333.1:1-1161/1-1161 GIWK01056333.1:5-1182/1-1178 GIWK01056333.1:35-1182/1-1148 GIWK01056333.1:77-1175/1-1099 GIWK01056333.1:1-1097/1-1097 GIWK01056333.1:12-993/1-982 GIWK01056333.1:102-1182/1-1081 GIWK01056333.1:182-1180/1-999 GIWK01056333.1:179-1183/1-1005 GIWK01056333.1:1-912/1-912 GIWK01056333.1:263-1182/1-920 GIWK01056333.1:266-1183/1-918 GIWK01056333.1:1-954/1-954 GIWK01056333.1:1-870/1-870 GIWK01056333.1:1-783/1-783 GIWK01056333.1:1-828/1-828 GIWK01056333.1:410-1175/1-766 GIWK01056333.1:368-1183/1-816 GIWK01056333.1:452-1182/1-731 GIWK01056333.1:1-744/1-744 GIWK01056333.1:1-702/1-702 GIWK01056333.1:12-660/1-649 GIWK01056333.1:506-1183/1-678 GIWK01056333.1:561-1183/1-623 GIWK01056333.1:599-1183/1-585 GIWK01056333.1:12-534/1-523 GIWK01056333.1:746-1182/1-437 GIWK01056333.1:1-366/1-366 GIWK01056333.1:1-198/1-198 GIWK01056333.1:1-159/1-159 GIWK01056333.1:1-117/1-117 GIWK01038917.1:5-1220/1-1216 GIWK01038917.1:5-944/1-940 GIWK01038917.1:2-944/1-943 GIWK01038917.1:7-944/1-938 GIWK01038917.1:2-944/1-943 GIWK01038917.1:5-944/1-940 GIWK01038917.1:1-944/1-944 GIWK01038917.1:31-944/1-914 GIWK01038917.1:5-944/1-940 GIWK01038917.1:5-944/1-940 GIWK01038917.1:5-934/1-930 GIWK01038917.1:2-944/1-943 GIWK01038917.1:1383-2534/1-1152 GIWK01038917.1:1-944/1-944 GIWK01038917.1:1880-3355/1-1476 GIWK01038917.1:2-944/1-943 GIWK01038917.1:5-860/1-856 GIWK01038917.1:2-944/1-943 GIWK01038917.1:25-944/1-920 GIWK01038917.1:1-808/1-808 GIWK01038917.1:2-776/1-775 GIWK01038917.1:169-944/1-776 GIWK01038917.1:92-942/1-851 GIWK01038917.1:172-944/1-773 GIWK01038917.1:5-706/1-702 GIWK01038917.1:2-650/1-649 GIWK01038917.1:1-692/1-692 GIWK01038917.1:253-944/1-692 GIWK01038917.1:256-944/1-689 GIWK01038917.1:334-944/1-611 GIWK01038917.1:5-482/1-478 GIWK01038917.1:2841-3509/1-669 GIWK01038917.1:2-521/1-520 GIWK01038917.1:1-356/1-356 GIWK01038917.1:3492-3748/1-257 GIWK01038917.1:610-934/1-325 GIWK01038917.1:2-188/1-187 GIWK01038917.1:5-149/1-145 GIWK01038917.1:5-107/1-103 GIWK01044708.1:12-1410/1-1399 GIWK01044708.1:1-1401/1-1401 GIWK01044708.1:1-1404/1-1404 GIWK01044708.1:1-1401/1-1401 GIWK01044708.1:63-1401/1-1339 GIWK01044708.1:6-1409/1-1404 GIWK01044708.1:1-1406/1-1406 GIWK01044708.1:1-1404/1-1404 GIWK01044708.1:1-1412/1-1412 GIWK01044708.1:1-1387/1-1387 GIWK01044708.1:1-1258/1-1258 GIWK01044708.1:1-1209/1-1209 GIWK01044708.1:1-1278/1-1278 GIWK01044708.1:129-1412/1-1284 GIWK01044708.1:213-1410/1-1198 GIWK01044708.1:1-1177/1-1177 GIWK01044708.1:234-1409/1-1176 GIWK01044708.1:297-1401/1-1105 GIWK01044708.1:1-1093/1-1093 GIWK01044708.1:1-1051/1-1051 GIWK01044708.1:1-1132/1-1132 GIWK01044708.1:423-1406/1-984 GIWK01044708.1:339-1409/1-1071 GIWK01044708.1:1-1009/1-1009 GIWK01044708.1:444-1409/1-966 GIWK01044708.1:507-1412/1-906 GIWK01044708.1:1-925/1-925 GIWK01044708.1:1-880/1-880 GIWK01044708.1:549-1406/1-858 GIWK01044708.1:1-841/1-841 GIWK01044708.1:591-1412/1-822 GIWK01044708.1:1-796/1-796 GIWK01044708.1:1-757/1-757 GIWK01044708.1:1-715/1-715 GIWK01044708.1:843-1412/1-570 GIWK01044708.1:696-1412/1-717 GIWK01044708.1:1-631/1-631 GIWK01044708.1:1-589/1-589 GIWK01044708.1:1-673/1-673 GIWK01044708.1:1-460/1-460 GIWK01044708.1:927-1409/1-483 GIWK01044708.1:1-379/1-379 GIWK01044708.1:1-411/1-411 GIWK01044708.1:1-295/1-295 GIWK01044708.1:1-250/1-250 GIWK01044708.1:1-211/1-211 GIWK01011613.1:191-2048/1-1858 GIWK01011613.1:1549-2051/1-503 GIWK01011613.1:1566-2057/1-492 GIWK01011613.1:1549-2057/1-509 GIWK01011613.1:1566-2057/1-492 GIWK01011613.1:1546-2057/1-512 GIWK01011613.1:1566-2057/1-492 GIWK01011613.1:1566-2057/1-492 GIWK01011613.1:1566-2057/1-492 GIWK01011613.1:1566-2057/1-492 GIWK01011613.1:1546-2040/1-495 GIWK01011613.1:1584-2057/1-474 GIWK01011613.1:1546-2057/1-512 GIWK01011613.1:1566-2026/1-461 GIWK01011613.1:1584-1816/1-233 GIWK01011613.1:1546-1771/1-226 GIWK01011613.1:1549-1732/1-184 GIWK01011613.1:163-198/1-36 GIWK01015682.1:1-1374/1-1374 GIWK01015682.1:1-1356/1-1356 GIWK01015682.1:1-1375/1-1375 GIWK01015682.1:1-1374/1-1374 GIWK01015682.1:1-1374/1-1374 GIWK01015682.1:2-1375/1-1374 GIWK01015682.1:1-1375/1-1375 GIWK01015682.1:9-1356/1-1348 GIWK01015682.1:3-1356/1-1354 GIWK01015682.1:1-1342/1-1342 GIWK01015682.1:1-1300/1-1300 GIWK01015682.1:28-1375/1-1348 GIWK01015682.1:1-1258/1-1258 GIWK01015682.1:66-1375/1-1310 GIWK01015682.1:129-1356/1-1228 GIWK01015682.1:1-1146/1-1146 GIWK01015682.1:1-1132/1-1132 GIWK01015682.1:1-1216/1-1216 GIWK01015682.1:154-1375/1-1222 GIWK01015682.1:213-1375/1-1163 GIWK01015682.1:1-1048/1-1048 GIWK01015682.1:1-1080/1-1080 GIWK01015682.1:1-964/1-964 GIWK01015682.1:238-1356/1-1119 GIWK01015682.1:297-1375/1-1079 GIWK01015682.1:1-1006/1-1006 GIWK01015682.1:339-1375/1-1037 GIWK01015682.1:1-835/1-835 GIWK01015682.1:441-1374/1-934 GIWK01015682.1:1-880/1-880 GIWK01015682.1:525-1375/1-851 GIWK01015682.1:1-754/1-754 GIWK01015682.1:528-1374/1-847 GIWK01015682.1:1-796/1-796 GIWK01015682.1:672-1375/1-704 GIWK01015682.1:714-1374/1-661 GIWK01015682.1:768-1375/1-608 GIWK01015682.1:1-628/1-628 GIWK01015682.1:1-460/1-460 GIWK01015682.1:823-1374/1-552 GIWK01015682.1:865-1374/1-510 GIWK01015682.1:1-421/1-421 GIWK01015682.1:1-379/1-379 GIWK01015682.1:1-250/1-250 GIWK01015682.1:1068-1375/1-308 GIWK01015682.1:1-169/1-169 GIWK01015682.1:1-127/1-127 GIWK01015682.1:1218-1374/1-157 GIWK01068433.1:597-1926/1-1330 GIWK01068433.1:745-1905/1-1161 GIWK01068433.1:774-1923/1-1150 GIWK01068433.1:762-1923/1-1162 GIWK01068433.1:745-1926/1-1182 GIWK01068433.1:762-1926/1-1165 GIWK01068433.1:762-1926/1-1165 GIWK01068433.1:745-1926/1-1182 GIWK01068433.1:762-1905/1-1144 GIWK01068433.1:742-1926/1-1185 GIWK01068433.1:762-1905/1-1144 GIWK01068433.1:762-1923/1-1162 GIWK01068433.1:762-1923/1-1162 GIWK01068433.1:829-1926/1-1098 GIWK01068433.1:762-1891/1-1130 GIWK01068433.1:745-1827/1-1083 GIWK01068433.1:888-1926/1-1039 GIWK01068433.1:913-1926/1-1014 GIWK01068433.1:742-1723/1-982 GIWK01068433.1:745-1684/1-940 GIWK01068433.1:984-1923/1-940 GIWK01068433.1:762-1642/1-881 GIWK01068433.1:1077-1923/1-847 GIWK01068433.1:762-1600/1-839 GIWK01068433.1:1140-1905/1-766 GIWK01068433.1:2-586/1-585 GIWK01068433.1:742-1558/1-817 GIWK01068433.1:1182-1926/1-745 GIWK01068433.1:1224-1926/1-703 GIWK01068433.1:742-1390/1-649 GIWK01068433.1:780-1429/1-650 GIWK01068433.1:1291-1926/1-636 GIWK01068433.1:1329-1926/1-598 GIWK01068433.1:762-1348/1-587 GIWK01068433.1:780-1306/1-527 GIWK01068433.1:742-1264/1-523 GIWK01068433.1:762-1222/1-461 GIWK01068433.1:1476-1926/1-451 GIWK01068433.1:780-1093/1-314 GIWK01068433.1:780-1012/1-233 GIWK01068433.1:742-967/1-226 GIWK01068433.1:745-928/1-184 GIWK01068433.1:1788-1923/1-136 GIWK01064514.1:1-1388/1-1388 GIWK01064514.1:1-1388/1-1388 GIWK01064514.1:22-1363/1-1342 GIWK01064514.1:1-1363/1-1363 GIWK01064514.1:1-1366/1-1366 GIWK01064514.1:1-1388/1-1388 GIWK01064514.1:10-1363/1-1354 GIWK01064514.1:1-1388/1-1388 GIWK01064514.1:1-1385/1-1385 GIWK01064514.1:1-1349/1-1349 GIWK01064514.1:1-1262/1-1262 GIWK01064514.1:1-1385/1-1385 GIWK01064514.1:52-1388/1-1337 GIWK01064514.1:77-1388/1-1312 GIWK01064514.1:119-1363/1-1245 GIWK01064514.1:1-1181/1-1181 GIWK01064514.1:1-1223/1-1223 GIWK01064514.1:157-1388/1-1232 GIWK01064514.1:220-1388/1-1169 GIWK01064514.1:1-1052/1-1052 GIWK01064514.1:1-1139/1-1139 GIWK01064514.1:1-971/1-971 GIWK01064514.1:262-1363/1-1102 GIWK01064514.1:322-1388/1-1067 GIWK01064514.1:406-1388/1-983 GIWK01064514.1:1-1013/1-1013 GIWK01064514.1:430-1386/1-957 GIWK01064514.1:472-1388/1-917 GIWK01064514.1:1-929/1-929 GIWK01064514.1:1-887/1-887 GIWK01064514.1:497-1388/1-892 GIWK01064514.1:1-845/1-845 GIWK01064514.1:658-1388/1-731 GIWK01064514.1:661-1388/1-728 GIWK01064514.1:1-761/1-761 GIWK01064514.1:739-1388/1-650 GIWK01064514.1:763-1388/1-626 GIWK01064514.1:1-554/1-554 GIWK01064514.1:830-1388/1-559 GIWK01064514.1:868-1371/1-504 GIWK01064514.1:1-400/1-400 GIWK01064514.1:1015-1388/1-374 GIWK01064514.1:1-302/1-302 GIWK01064514.1:1-334/1-334 GIWK01064514.1:1-218/1-218 GIWK01064514.1:1-134/1-134 GIWK01064060.1:7-1462/1-1456 GIWK01064060.1:2-1467/1-1466 GIWK01064060.1:4-1459/1-1456 GIWK01064060.1:2-1464/1-1463 GIWK01064060.1:1-1445/1-1445 GIWK01064060.1:5-1464/1-1460 GIWK01064060.1:5-1462/1-1458 GIWK01064060.1:67-1466/1-1400 GIWK01064060.1:31-1459/1-1429 GIWK01064060.1:5-1467/1-1463 GIWK01064060.1:2-1381/1-1380 GIWK01064060.1:2-1364/1-1363 GIWK01064060.1:169-1466/1-1298 GIWK01064060.1:109-1459/1-1351 GIWK01064060.1:1-1277/1-1277 GIWK01064060.1:193-1467/1-1275 GIWK01064060.1:235-1459/1-1225 GIWK01064060.1:5-1196/1-1192 GIWK01064060.1:260-1466/1-1207 GIWK01064060.1:5-1154/1-1150 GIWK01064060.1:1-1238/1-1238 GIWK01064060.1:302-1466/1-1165 GIWK01064060.1:5-1112/1-1108 GIWK01064060.1:5-1067/1-1063 GIWK01064060.1:2-986/1-985 GIWK01064060.1:403-1466/1-1064 GIWK01064060.1:466-1464/1-999 GIWK01064060.1:463-1467/1-1005 GIWK01064060.1:1-1028/1-1028 GIWK01064060.1:5-944/1-940 GIWK01064060.1:544-1466/1-923 GIWK01064060.1:5-860/1-856 GIWK01064060.1:652-1467/1-816 GIWK01064060.1:1-818/1-818 GIWK01064060.1:5-902/1-898 GIWK01064060.1:2-776/1-775 GIWK01064060.1:706-1459/1-754 GIWK01064060.1:736-1466/1-731 GIWK01064060.1:5-734/1-730 GIWK01064060.1:778-1467/1-690 GIWK01064060.1:845-1467/1-623 GIWK01064060.1:7-566/1-560 GIWK01064060.1:883-1467/1-585 GIWK01064060.1:2-401/1-400 GIWK01064060.1:1030-1466/1-437 GIWK01064060.1:2-317/1-316 GIWK01064060.1:7-272/1-266 GIWK01064060.1:2-181/1-180 GIWK01064060.1:5-149/1-145 GIWK01029243.1:576-1528/1-953 GIWK01029243.1:1-756/1-756 GIWK01066786.1:501-1455/1-955 GIWK01066786.1:1-628/1-628 GIWK01000457.1:4-1352/1-1349 GIWK01000457.1:28-1352/1-1325 GIWK01000457.1:4-1339/1-1336 GIWK01000457.1:1-1336/1-1336 GIWK01000457.1:4-1339/1-1336 GIWK01000457.1:1-1319/1-1319 GIWK01000457.1:5-1336/1-1332 GIWK01000457.1:47-1352/1-1306 GIWK01000457.1:4-1352/1-1349 GIWK01000457.1:4-1280/1-1277 GIWK01000457.1:22-1352/1-1331 GIWK01000457.1:4-1238/1-1235 GIWK01000457.1:127-1339/1-1213 GIWK01000457.1:89-1336/1-1248 GIWK01000457.1:190-1352/1-1163 GIWK01000457.1:376-1352/1-977 GIWK01000457.1:4-1154/1-1151 GIWK01000457.1:4-1109/1-1106 GIWK01000457.1:4-1070/1-1067 GIWK01000457.1:232-1336/1-1105 GIWK01000457.1:292-1352/1-1061 GIWK01000457.1:4-1028/1-1025 GIWK01000457.1:4-986/1-983 GIWK01000457.1:4-944/1-941 GIWK01000457.1:4-902/1-899 GIWK01000457.1:467-1352/1-886 GIWK01000457.1:442-1352/1-911 GIWK01000457.1:509-1352/1-844 GIWK01000457.1:568-1336/1-769 GIWK01000457.1:4-793/1-790 GIWK01000457.1:593-1352/1-760 GIWK01000457.1:4-724/1-721 GIWK01000457.1:694-1352/1-659 GIWK01000457.1:4-692/1-689 GIWK01000457.1:4-608/1-605 GIWK01000457.1:736-1352/1-617 GIWK01000457.1:757-1339/1-583 GIWK01000457.1:820-1352/1-533 GIWK01000457.1:4-647/1-644 GIWK01000457.1:4-524/1-521 GIWK01000457.1:904-1339/1-436 GIWK01000457.1:4-370/1-367 GIWK01000457.1:4-304/1-301 GIWK01000457.1:1-272/1-272 GIWK01000457.1:1072-1352/1-281 GIWK01000457.1:4-188/1-185 GIWK01000457.1:4-104/1-101 GIWK01052658.1:33-1024/1-992 GIWK01052658.1:1285-2252/1-968 GIWK01052658.1:819-1859/1-1041 GIWK01063328.1:26-1464/1-1439 GIWK01063328.1:285-1445/1-1161 GIWK01063328.1:314-1463/1-1150 GIWK01063328.1:302-1463/1-1162 GIWK01063328.1:285-1464/1-1180 GIWK01063328.1:302-1464/1-1163 GIWK01063328.1:302-1464/1-1163 GIWK01063328.1:285-1464/1-1180 GIWK01063328.1:302-1445/1-1144 GIWK01063328.1:302-1445/1-1144 GIWK01063328.1:282-1464/1-1183 GIWK01063328.1:302-1463/1-1162 GIWK01063328.1:302-1463/1-1162 GIWK01063328.1:369-1464/1-1096 GIWK01063328.1:302-1431/1-1130 GIWK01063328.1:285-1367/1-1083 GIWK01063328.1:428-1464/1-1037 GIWK01063328.1:453-1464/1-1012 GIWK01063328.1:282-1263/1-982 GIWK01063328.1:285-1224/1-940 GIWK01063328.1:524-1463/1-940 GIWK01063328.1:302-1182/1-881 GIWK01063328.1:302-1140/1-839 GIWK01063328.1:617-1463/1-847 GIWK01063328.1:680-1445/1-766 GIWK01063328.1:282-1098/1-817 GIWK01063328.1:722-1464/1-743 GIWK01063328.1:302-1053/1-752 GIWK01063328.1:764-1464/1-701 GIWK01063328.1:282-930/1-649 GIWK01063328.1:320-972/1-653 GIWK01063328.1:831-1464/1-634 GIWK01063328.1:869-1464/1-596 GIWK01063328.1:302-888/1-587 GIWK01063328.1:320-846/1-527 GIWK01063328.1:282-804/1-523 GIWK01063328.1:302-762/1-461 GIWK01063328.1:1016-1464/1-449 GIWK01063328.1:320-552/1-233 GIWK01063328.1:282-507/1-226 GIWK01063328.1:285-468/1-184 GIWK01063328.1:1328-1463/1-136 GIWK01015959.1:9-1424/1-1416 GIWK01015959.1:11-1435/1-1425 GIWK01015959.1:77-1435/1-1359 GIWK01015959.1:9-1424/1-1416 GIWK01015959.1:50-1435/1-1386 GIWK01015959.1:9-1410/1-1402 GIWK01015959.1:9-1424/1-1416 GIWK01015959.1:9-1424/1-1416 GIWK01015959.1:53-1424/1-1372 GIWK01015959.1:47-1424/1-1378 GIWK01015959.1:9-1365/1-1357 GIWK01015959.1:9-1284/1-1276 GIWK01015959.1:236-1424/1-1189 GIWK01015959.1:11-1326/1-1316 GIWK01015959.1:152-1424/1-1273 GIWK01015959.1:194-1435/1-1242 GIWK01015959.1:9-1200/1-1192 GIWK01015959.1:9-1155/1-1147 GIWK01015959.1:320-1435/1-1116 GIWK01015959.1:290-1424/1-1135 GIWK01015959.1:9-1074/1-1066 GIWK01015959.1:9-1032/1-1024 GIWK01015959.1:345-1424/1-1080 GIWK01015959.1:383-1423/1-1041 GIWK01015959.1:9-948/1-940 GIWK01015959.1:9-864/1-856 GIWK01015959.1:488-1424/1-937 GIWK01015959.1:572-1424/1-853 GIWK01015959.1:9-990/1-982 GIWK01015959.1:548-1424/1-877 GIWK01015959.1:9-906/1-898 GIWK01015959.1:632-1424/1-793 GIWK01015959.1:9-755/1-747 GIWK01015959.1:723-1424/1-702 GIWK01015959.1:782-1424/1-643 GIWK01015959.1:9-626/1-618 GIWK01015959.1:824-1424/1-601 GIWK01015959.1:9-728/1-720 GIWK01015959.1:866-1424/1-559 GIWK01015959.1:11-560/1-550 GIWK01015959.1:9-528/1-520 GIWK01015959.1:933-1424/1-492 GIWK01015959.1:9-399/1-391 GIWK01015959.1:1118-1424/1-307 GIWK01015959.1:9-318/1-310 GIWK01015959.1:9-150/1-142 GIWK01065725.1:140-1685/1-1546 GIWK01065725.1:56-1697/1-1642 GIWK01065725.1:2-1668/1-1667 GIWK01065725.1:1-1604/1-1604 GIWK01065725.1:80-1697/1-1618 GIWK01065725.1:8-1697/1-1690 GIWK01065725.1:164-1682/1-1519 GIWK01065725.1:1-1500/1-1500 GIWK01065725.1:231-1682/1-1452 GIWK01065725.1:347-1682/1-1336 GIWK01065725.1:2-1461/1-1460 GIWK01065725.1:273-1697/1-1425 GIWK01065725.1:1-1419/1-1419 GIWK01065725.1:1-1335/1-1335 GIWK01065725.1:371-1697/1-1327 GIWK01065725.1:413-1697/1-1285 GIWK01065725.1:455-1682/1-1228 GIWK01065725.1:1-1290/1-1290 GIWK01065725.1:1-1251/1-1251 GIWK01065725.1:1-1209/1-1209 GIWK01065725.1:509-1689/1-1181 GIWK01065725.1:539-1697/1-1159 GIWK01065725.1:1-1125/1-1125 GIWK01065725.1:1-1167/1-1167 GIWK01065725.1:606-1697/1-1092 GIWK01065725.1:1-1041/1-1041 GIWK01065725.1:707-1697/1-991 GIWK01065725.1:644-1697/1-1054 GIWK01065725.1:1-1083/1-1083 GIWK01065725.1:809-1697/1-889 GIWK01065725.1:749-1697/1-949 GIWK01065725.1:1-873/1-873 GIWK01065725.1:893-1697/1-805 GIWK01065725.1:1-887/1-887 GIWK01065725.1:917-1682/1-766 GIWK01065725.1:1-789/1-789 GIWK01065725.1:959-1697/1-739 GIWK01065725.1:984-1697/1-714 GIWK01065725.1:1-821/1-821 GIWK01065725.1:1-705/1-705 GIWK01065725.1:1068-1697/1-630 GIWK01065725.1:1106-1694/1-589 GIWK01065725.1:20-537/1-518 GIWK01065725.1:1-621/1-621 GIWK01065725.1:1253-1689/1-437 GIWK01065725.1:1-268/1-268 GIWK01065725.1:1-134/1-134 GIWK01065725.1:1565-1697/1-133 GIWK01067018.1:3-1421/1-1419 GIWK01067018.1:1-1421/1-1421 GIWK01067018.1:29-1423/1-1395 GIWK01067018.1:23-1425/1-1403 GIWK01067018.1:5-1418/1-1414 GIWK01067018.1:65-1418/1-1354 GIWK01067018.1:3-1423/1-1421 GIWK01067018.1:1-1427/1-1427 GIWK01067018.1:167-1427/1-1261 GIWK01067018.1:3-1404/1-1402 GIWK01067018.1:90-1425/1-1336 GIWK01067018.1:1-1340/1-1340 GIWK01067018.1:191-1418/1-1228 GIWK01067018.1:1-1281/1-1281 GIWK01067018.1:5-1323/1-1319 GIWK01067018.1:1-1236/1-1236 GIWK01067018.1:14-1197/1-1184 GIWK01067018.1:275-1425/1-1151 GIWK01067018.1:251-1425/1-1175 GIWK01067018.1:317-1418/1-1102 GIWK01067018.1:3-1155/1-1153 GIWK01067018.1:3-1113/1-1111 GIWK01067018.1:461-1423/1-963 GIWK01067018.1:401-1427/1-1027 GIWK01067018.1:1-987/1-987 GIWK01067018.1:485-1425/1-941 GIWK01067018.1:1-1026/1-1026 GIWK01067018.1:3-945/1-943 GIWK01067018.1:3-903/1-901 GIWK01067018.1:3-861/1-859 GIWK01067018.1:611-1427/1-817 GIWK01067018.1:3-777/1-775 GIWK01067018.1:636-1418/1-783 GIWK01067018.1:674-1425/1-752 GIWK01067018.1:1-819/1-819 GIWK01067018.1:737-1427/1-691 GIWK01067018.1:3-690/1-688 GIWK01067018.1:3-735/1-733 GIWK01067018.1:779-1427/1-649 GIWK01067018.1:804-1427/1-624 GIWK01067018.1:5-539/1-535 GIWK01067018.1:846-1427/1-582 GIWK01067018.1:3-473/1-471 GIWK01067018.1:1-399/1-399 GIWK01067018.1:989-1425/1-437 GIWK01067018.1:3-441/1-439 GIWK01067018.1:5-245/1-241 GIWK01067018.1:3-147/1-145 GHVE01065588.1:1-640/1-640 GHVE01065588.1:2530-3273/1-744 GHVE01065588.1:1257-2244/1-988 GHVE01065588.1:1635-2639/1-1005 GIWK01069519.1:628-1657/1-1030 GIWK01031270.1:9-1475/1-1467 GIWK01031270.1:1-1469/1-1469 GIWK01031270.1:1-1471/1-1471 GIWK01031270.1:1-1475/1-1475 GIWK01031270.1:1-1469/1-1469 GIWK01031270.1:9-1471/1-1463 GIWK01031270.1:9-1410/1-1402 GIWK01031270.1:35-1466/1-1432 GIWK01031270.1:9-1452/1-1444 GIWK01031270.1:1-1326/1-1326 GIWK01031270.1:12-1466/1-1455 GIWK01031270.1:1-1242/1-1242 GIWK01031270.1:50-1473/1-1424 GIWK01031270.1:113-1466/1-1354 GIWK01031270.1:9-1365/1-1357 GIWK01031270.1:138-1473/1-1336 GIWK01031270.1:197-1475/1-1279 GIWK01031270.1:1-1284/1-1284 GIWK01031270.1:1-1158/1-1158 GIWK01031270.1:222-1466/1-1245 GIWK01031270.1:323-1473/1-1151 GIWK01031270.1:1-1200/1-1200 GIWK01031270.1:1-1116/1-1116 GIWK01031270.1:348-1466/1-1119 GIWK01031270.1:425-1473/1-1049 GIWK01031270.1:1-1004/1-1004 GIWK01031270.1:1-1074/1-1074 GIWK01031270.1:1-938/1-938 GIWK01031270.1:509-1471/1-963 GIWK01031270.1:512-1473/1-962 GIWK01031270.1:9-990/1-982 GIWK01031270.1:1-906/1-906 GIWK01031270.1:1-819/1-819 GIWK01031270.1:590-1475/1-886 GIWK01031270.1:656-1475/1-820 GIWK01031270.1:614-1475/1-862 GIWK01031270.1:1-864/1-864 GIWK01031270.1:1-738/1-738 GIWK01031270.1:698-1466/1-769 GIWK01031270.1:9-780/1-772 GIWK01031270.1:782-1475/1-694 GIWK01031270.1:752-1473/1-722 GIWK01031270.1:807-1475/1-669 GIWK01031270.1:1-612/1-612 GIWK01031270.1:926-1475/1-550 GIWK01031270.1:950-1473/1-524 GIWK01031270.1:1010-1473/1-464 GIWK01031270.1:1-444/1-444 GIWK01031270.1:1076-1471/1-396 GIWK01031270.1:9-405/1-397 GIWK01031270.1:1-363/1-363 GIWK01031270.1:9-234/1-226 GIWK01031270.1:1-153/1-153 GIWK01031270.1:1-111/1-111 GIWK01031270.1:1328-1475/1-148 GIWK01014775.1:9-1487/1-1479 GIWK01014775.1:9-1487/1-1479 GIWK01014775.1:5-1487/1-1483 GIWK01014775.1:89-1487/1-1399 GIWK01014775.1:9-1487/1-1479 GIWK01014775.1:11-1487/1-1477 GIWK01014775.1:5-1487/1-1483 GIWK01014775.1:6-1445/1-1440 GIWK01014775.1:35-1487/1-1453 GIWK01014775.1:155-1487/1-1333 GIWK01014775.1:5-1403/1-1399 GIWK01014775.1:5-1361/1-1357 GIWK01014775.1:11-1319/1-1309 GIWK01014775.1:180-1487/1-1308 GIWK01014775.1:222-1487/1-1266 GIWK01014775.1:6-1210/1-1205 GIWK01014775.1:9-1190/1-1182 GIWK01014775.1:9-1141/1-1133 GIWK01014775.1:281-1487/1-1207 GIWK01014775.1:365-1487/1-1123 GIWK01014775.1:390-1487/1-1098 GIWK01014775.1:9-1109/1-1101 GIWK01014775.1:9-1064/1-1056 GIWK01014775.1:491-1487/1-997 GIWK01014775.1:6-1025/1-1020 GIWK01014775.1:533-1487/1-955 GIWK01014775.1:3-983/1-981 GIWK01014775.1:9-802/1-794 GIWK01014775.1:6-800/1-795 GIWK01014775.1:842-1487/1-646 GIWK01014775.1:9-941/1-933 GIWK01014775.1:9-738/1-730 GIWK01014775.1:5-783/1-779 GIWK01014775.1:801-1487/1-687 GIWK01014775.1:638-1487/1-850 GIWK01014775.1:6-699/1-694 GIWK01014775.1:1-657/1-657 GIWK01014775.1:6-615/1-610 GIWK01014775.1:926-1487/1-562 GIWK01014775.1:2-531/1-530 GIWK01014775.1:11-573/1-563 GIWK01014775.1:985-1487/1-503 GIWK01014775.1:1-402/1-402 GIWK01014775.1:1-321/1-321 GIWK01014775.1:6-363/1-358 GIWK01014775.1:1-237/1-237 GIWK01014775.1:6-185/1-180 GIWK01014775.1:1321-1487/1-167 GIWK01064888.1:396-1456/1-1061 GIWK01064888.1:1-680/1-680 GHVE01065598.1:1-744/1-744 GHVE01065598.1:2634-3377/1-744 GHVE01065598.1:1361-2348/1-988 GHVE01065598.1:1739-2743/1-1005 GIWK01019425.1:886-1992/1-1107 GIWK01019425.1:1-1221/1-1221 GIWK01006535.1:822-2127/1-1306 GIWK01006535.1:2-1342/1-1341 GHVE01065605.1:1-811/1-811 GHVE01065605.1:2701-3444/1-744 GHVE01065605.1:1428-2415/1-988 GHVE01065605.1:1806-2810/1-1005 GHVE01065596.1:1-923/1-923 GHVE01065596.1:2813-3556/1-744 GHVE01065596.1:1540-2527/1-988 GHVE01065596.1:1918-2922/1-1005 GHVE01065589.1:1-729/1-729 GHVE01065589.1:2619-3362/1-744 GHVE01065589.1:1346-2333/1-988 GHVE01065589.1:1724-2728/1-1005 GIWK01000094.1:43-1376/1-1334 GIWK01000094.1:1-596/1-596 GIWK01061984.1:7-1625/1-1619 GIWK01061984.1:1-1659/1-1659 GIWK01061984.1:1-1586/1-1586 GIWK01061984.1:22-1659/1-1638 GIWK01061984.1:121-1645/1-1525 GIWK01061984.1:118-1659/1-1542 GIWK01061984.1:1-1544/1-1544 GIWK01061984.1:205-1659/1-1455 GIWK01061984.1:202-1642/1-1441 GIWK01061984.1:1-1460/1-1460 GIWK01061984.1:283-1642/1-1360 GIWK01061984.1:1-1415/1-1415 GIWK01061984.1:1-1376/1-1376 GIWK01061984.1:2-1292/1-1291 GIWK01061984.1:349-1659/1-1311 GIWK01061984.1:647-1659/1-1013 GIWK01061984.1:391-1642/1-1252 GIWK01061984.1:475-1659/1-1185 GIWK01061984.1:445-1645/1-1201 GIWK01061984.1:1-1208/1-1208 GIWK01061984.1:645-1659/1-1015 GIWK01061984.1:1-1166/1-1166 GIWK01061984.1:517-1642/1-1126 GIWK01061984.1:1-1124/1-1124 GIWK01061984.1:731-1659/1-929 GIWK01061984.1:1-1082/1-1082 GIWK01061984.1:790-1659/1-870 GIWK01061984.1:1-946/1-946 GIWK01061984.1:815-1659/1-845 GIWK01061984.1:886-1642/1-757 GIWK01061984.1:1-869/1-869 GIWK01061984.1:1-830/1-830 GIWK01061984.1:1-914/1-914 GIWK01061984.1:979-1659/1-681 GIWK01061984.1:1042-1659/1-618 GIWK01061984.1:1-788/1-788 GIWK01061984.1:1-655/1-655 GIWK01061984.1:1084-1645/1-562 GIWK01061984.1:1-579/1-579 GIWK01061984.1:1126-1659/1-534 GIWK01061984.1:1-547/1-547 GIWK01061984.1:1-473/1-473 GIWK01061984.1:1378-1659/1-282 GIWK01061984.1:1-305/1-305 GIWK01061984.1:1-137/1-137 GIWK01061984.1:1-42/1-42 GIWK01032289.1:320-1615/1-1296 GIWK01032289.1:1-832/1-832 GIWK01032646.1:936-2223/1-1288 GIWK01032646.1:1-1444/1-1444 GIWK01067527.1:652-1939/1-1288 GIWK01067527.1:1-1164/1-1164 GIWK01026002.1:556-1857/1-1302 GIWK01026002.1:6-1070/1-1065 GIWK01029994.1:876-2146/1-1271 GIWK01012354.1:1073-2370/1-1298 GIWK01012354.1:112-1587/1-1476 GIWK01042089.1:33-1330/1-1298 GIWK01042089.1:1637-2785/1-1149 GIWK01042089.1:822-2289/1-1468 GIWK01042089.1:2949-3093/1-145 GIWK01043700.1:525-1818/1-1294 GIWK01043700.1:1-1039/1-1039 GIWK01000702.1:428-1719/1-1292 GIWK01000702.1:8-940/1-933 GIWK01017897.1:1299-2594/1-1296 GIWK01017897.1:338-1809/1-1472 GIWK01017897.1:1-992/1-992 GIWK01039335.1:2025-3316/1-1292 GIWK01039335.1:1108-2539/1-1432 GIWK01039335.1:658-1801/1-1144 GIWK01039335.1:1-660/1-660 GIWK01039335.1:1-660/1-660 GIWK01039335.1:1-661/1-661 GIWK01039335.1:19-660/1-642 GIWK01039335.1:1-660/1-660 GIWK01039335.1:1-660/1-660 GIWK01039335.1:1-660/1-660 GIWK01039335.1:1-660/1-660 GIWK01039335.1:1-637/1-637 GIWK01039335.1:1-660/1-660 GIWK01039335.1:1-659/1-659 GIWK01039335.1:1-660/1-660 GIWK01039335.1:1-660/1-660 GIWK01039335.1:1-660/1-660 GIWK01039335.1:1-660/1-660 GIWK01039335.1:37-660/1-624 GIWK01039335.1:7-660/1-654 GIWK01039335.1:1-620/1-620 GIWK01039335.1:79-660/1-582 GIWK01039335.1:1-581/1-581 GIWK01039335.1:121-660/1-540 GIWK01039335.1:19-539/1-521 GIWK01039335.1:1-287/1-287 GIWK01039335.1:19-326/1-308 GIWK01039335.1:373-661/1-289 GIWK01039335.1:19-245/1-227 GIWK01039335.1:2-203/1-202 GIWK01039335.1:1-161/1-161 GIWK01039335.1:2-119/1-118 GIWK01025856.1:779-2027/1-1249 GIWK01025856.1:389-1246/1-858 GIWK01025856.1:1-225/1-225 GIWK01063059.1:110-1400/1-1291 GIWK01063059.1:1-622/1-622 GHVE01065593.1:5-1053/1-1049 GHVE01065593.1:2943-3686/1-744 GHVE01065593.1:1670-2657/1-988 GHVE01065593.1:2048-3052/1-1005 GIWK01026969.1:987-2333/1-1347 GIWK01026969.1:398-1550/1-1153 GIWK01026969.1:1-854/1-854 GIWK01056241.1:607-1954/1-1348 GIWK01056241.1:1-1172/1-1172 GIWK01036431.1:31-1380/1-1350 GIWK01036431.1:818-2236/1-1419 GIWK01036431.1:1651-2783/1-1133 GIWK01036431.1:2942-3149/1-208 GIWK01000847.1:212-1622/1-1411 GIWK01008296.1:1454-2795/1-1342 GIWK01008296.1:447-1333/1-887 GIWK01008296.1:840-2016/1-1177 GIWK01008296.1:1-289/1-289 GIWK01049141.1:1975-3314/1-1340 GIWK01049141.1:1121-2539/1-1419 GIWK01049141.1:574-1704/1-1131 GIWK01049141.1:4-549/1-546 GIWK01049141.1:4-549/1-546 GIWK01049141.1:4-538/1-535 GIWK01049141.1:4-549/1-546 GIWK01049141.1:1-546/1-546 GIWK01049141.1:4-546/1-543 GIWK01049141.1:4-543/1-540 GIWK01049141.1:4-543/1-540 GIWK01049141.1:4-504/1-501 GIWK01049141.1:4-538/1-535 GIWK01049141.1:1-543/1-543 GIWK01049141.1:4-543/1-540 GIWK01049141.1:4-543/1-540 GIWK01049141.1:4-541/1-538 GIWK01049141.1:5-543/1-539 GIWK01049141.1:28-543/1-516 GIWK01049141.1:4-370/1-367 GIWK01049141.1:4-304/1-301 GIWK01049141.1:1-272/1-272 GIWK01049141.1:4-188/1-185 GIWK01049141.1:292-543/1-252 GIWK01049141.1:4-104/1-101 GIWK01000327.1:398-1744/1-1347 GIWK01000327.1:1-962/1-962 GHVE01065606.1:2-1043/1-1042 GHVE01065606.1:2933-3676/1-744 GHVE01065606.1:1660-2647/1-988 GHVE01065606.1:2038-3042/1-1005 GIWK01065045.1:661-2013/1-1353 GIWK01065045.1:1-1226/1-1226 GIWK01050953.1:1048-2394/1-1347 GIWK01050953.1:192-1612/1-1421 GIWK01000290.1:277-1623/1-1347 GIWK01000290.1:1-842/1-842 GHVE01065599.1:149-1037/1-889 GHVE01065599.1:2927-3670/1-744 GHVE01065599.1:1654-2641/1-988 GHVE01065599.1:2032-3036/1-1005 GIWK01001255.1:71-1410/1-1340 GIWK01001255.1:6-635/1-630 GIWK01033542.1:806-2151/1-1346 GIWK01033542.1:2-1368/1-1367 GIWK01046963.1:2547-3892/1-1346 GIWK01046963.1:1-986/1-986 GIWK01046963.1:1691-3111/1-1421 GIWK01046963.1:1145-2276/1-1132 GIWK01046963.1:1-703/1-703 GIWK01046963.1:1-703/1-703 GIWK01046963.1:1-703/1-703 GIWK01046963.1:13-703/1-691 GIWK01046963.1:1-703/1-703 GIWK01046963.1:1-703/1-703 GIWK01046963.1:1-701/1-701 GIWK01046963.1:1-703/1-703 GIWK01046963.1:1-703/1-703 GIWK01046963.1:1-703/1-703 GIWK01046963.1:1-703/1-703 GIWK01046963.1:1-703/1-703 GIWK01046963.1:1-703/1-703 GIWK01046963.1:3-703/1-701 GIWK01046963.1:1-703/1-703 GIWK01046963.1:1-599/1-599 GIWK01046963.1:33-703/1-671 GIWK01046963.1:87-703/1-617 GIWK01046963.1:1-465/1-465 GIWK01046963.1:184-703/1-520 GIWK01046963.1:1-367/1-367 GIWK01046963.1:13-399/1-387 GIWK01046963.1:1-283/1-283 GIWK01046963.1:387-693/1-307 GIWK01046963.1:1-199/1-199 GIWK01046963.1:1-115/1-115 GIWK01039616.1:2107-3457/1-1351 GIWK01039616.1:1251-2672/1-1422 GIWK01039616.1:704-1836/1-1133 GIWK01039616.1:1-545/1-545 GIWK01039616.1:2-262/1-261 GIWK01039616.1:2-262/1-261 GIWK01039616.1:3-262/1-260 GIWK01039616.1:36-252/1-217 GIWK01039616.1:36-262/1-227 GIWK01039616.1:2-262/1-261 GIWK01039616.1:2-262/1-261 GIWK01039616.1:1-262/1-262 GIWK01039616.1:2-158/1-157 GHVE01065586.1:66-1052/1-987 GHVE01065586.1:2942-3937/1-996 GHVE01065586.1:1669-2656/1-988 GHVE01065586.1:2047-3051/1-1005 GIWK01032908.1:777-1688/1-912 GIWK01032908.1:1-622/1-622 GIWK01032908.1:1-346/1-346 GIWK01032908.1:2-346/1-345 GIWK01032908.1:1-346/1-346 GIWK01032908.1:1-346/1-346 GIWK01032908.1:1-346/1-346 GIWK01032908.1:1-346/1-346 GIWK01032908.1:1-346/1-346 GIWK01032908.1:1-346/1-346 GIWK01032908.1:12-336/1-325 GIWK01032908.1:1-262/1-262 GIWK01032908.1:2-108/1-107 GIWK01032908.1:2-94/1-93 GIWK01004636.1:773-2116/1-1344 GIWK01004636.1:135-1338/1-1204 GIWK01004636.1:1-648/1-648 GIWK01005258.1:1007-2053/1-1047 GIWK01005258.1:46-1519/1-1474 GHVE01065604.1:16-1054/1-1039 GHVE01065604.1:2944-3687/1-744 GHVE01065604.1:1671-2658/1-988 GHVE01065604.1:2049-3053/1-1005 GIWK01039343.1:1-1476/1-1476 GIWK01039343.1:1747-3088/1-1342 GIWK01039343.1:891-2309/1-1419 GIWK01039343.1:1-47/1-47 GHVE01065583.1:39-854/1-816 GHVE01065583.1:2744-3718/1-975 GHVE01065583.1:1471-2458/1-988 GHVE01065583.1:1849-2853/1-1005 GIWK01047599.1:1-1091/1-1091 GIWK01047599.1:518-1373/1-856 GKQY01000390.1:164-1646/1-1483 GIWK01061608.1:286-1353/1-1068 GIWK01061608.1:1-850/1-850 GIWK01061608.1:1376-1502/1-127 GIWK01053399.1:865-2711/1-1847 GIWK01053399.1:2146-3236/1-1091 GIWK01053399.1:1-706/1-706 GIWK01053399.1:1-423/1-423 GIWK01053399.1:1-423/1-423 GIWK01053399.1:1-423/1-423 GIWK01053399.1:1-423/1-423 GIWK01053399.1:1-423/1-423 GIWK01053399.1:1-421/1-421 GIWK01053399.1:1-423/1-423 GIWK01053399.1:11-423/1-413 GIWK01053399.1:1-423/1-423 GIWK01053399.1:1-319/1-319 GIWK01053399.1:107-413/1-307 GIWK01053399.1:1-185/1-185 GIWK01053399.1:1-119/1-119 GIWK01053399.1:1-87/1-87 GHVE01065620.1:406-1878/1-1473 GHVE01065581.1:664-2136/1-1473 GHVE01065581.1:5338-6898/1-1561 GHVE01065581.1:2341-3448/1-1108 GHVE01065581.1:4065-5052/1-988 GHVE01065581.1:4443-5447/1-1005 GHVE01065580.1:619-2091/1-1473 GHVE01065580.1:4975-6535/1-1561 GHVE01065580.1:2298-3085/1-788 GHVE01065580.1:3702-4689/1-988 GHVE01065580.1:4080-5084/1-1005 GHVE01065579.1:524-1996/1-1473 GHVE01065579.1:4577-6214/1-1638 GHVE01065579.1:3304-4291/1-988 GHVE01065579.1:3682-4686/1-1005 GHVE01065579.1:2220-2687/1-468 GIWK01022639.1:215-1365/1-1151 GIWK01022639.1:713-2186/1-1474 GIWK01022639.1:1672-2464/1-793 GKQY01000394.1:164-1868/1-1705 GIWK01037455.1:1260-2677/1-1418 GIWK01037455.1:2114-3221/1-1108 GHVE01065578.1:3053-4690/1-1638 GHVE01065578.1:99-1163/1-1065 GHVE01065578.1:1780-2767/1-988 GHVE01065578.1:2158-3162/1-1005 GIWK01046137.1:1566-2985/1-1420 GIWK01046137.1:19-861/1-843 GIWK01046137.1:2420-3515/1-1096 GIWK01046137.1:1020-2149/1-1130 GIWK01046137.1:1-578/1-578 GIWK01046137.1:1-578/1-578 GIWK01046137.1:1-578/1-578 GIWK01046137.1:1-578/1-578 GIWK01046137.1:1-578/1-578 GIWK01046137.1:1-578/1-578 GIWK01046137.1:1-578/1-578 GIWK01046137.1:1-578/1-578 GIWK01046137.1:1-578/1-578 GIWK01046137.1:61-578/1-518 GIWK01046137.1:1-578/1-578 GIWK01046137.1:55-578/1-524 GIWK01046137.1:1-578/1-578 GIWK01046137.1:1-474/1-474 GIWK01046137.1:58-578/1-521 GIWK01046137.1:1-340/1-340 GIWK01046137.1:262-568/1-307 GIWK01046137.1:1-158/1-158 GIWK01009859.1:83-1500/1-1418 GIWK01009859.1:937-2027/1-1091 GIWK01028724.1:525-1957/1-1433 GIWK01028724.1:1-1087/1-1087 GIWK01027668.1:1-1312/1-1312 GKQY01000391.1:164-3431/1-3268 GKQY01000393.1:164-3431/1-3268 GKQY01000389.1:164-3448/1-3285 GKQY01000392.1:164-2083/1-1920
