## Supplementary figures and images for "Tracing the vertebrate selenoproteome evolution reveals expansions in ray-finned fishes and convergent depletions in tetrapods"

### FIGURE_S2_SEPHS_SYNTENY_BIRDS.pdf

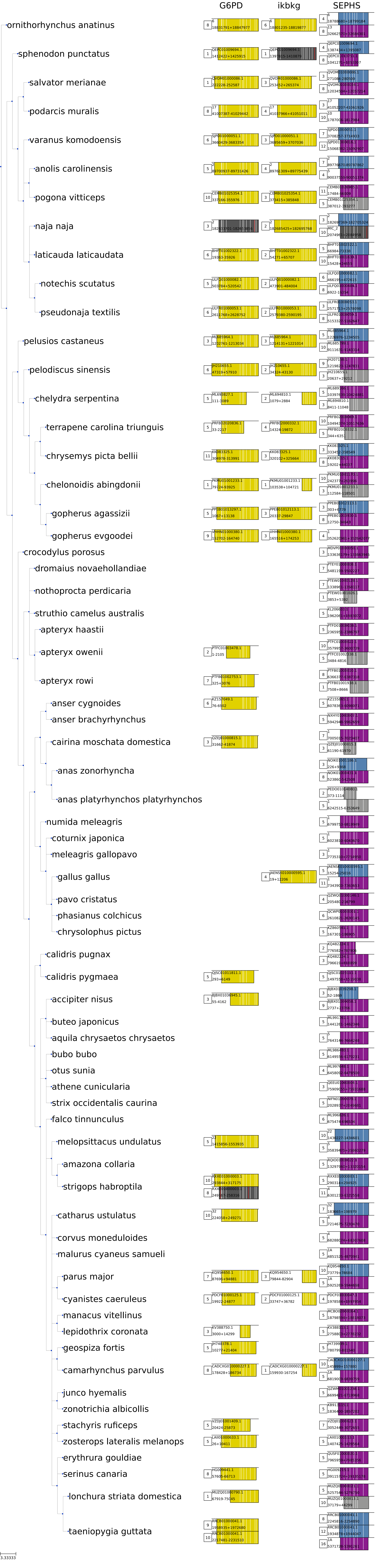

### FIGURE_S4_DIO.pdf

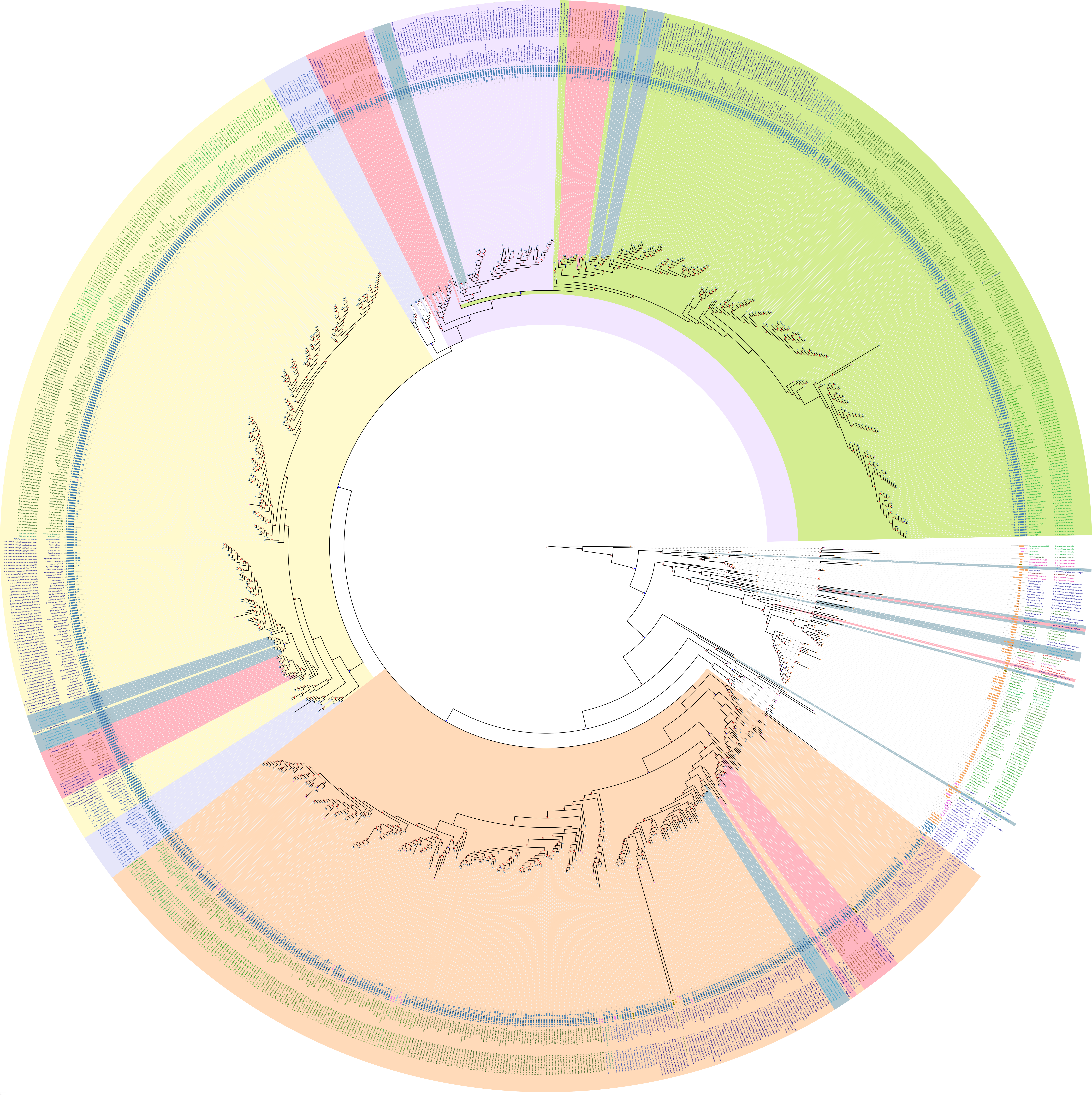

### FIGURE_S5_DIO_DRAWER.pdf

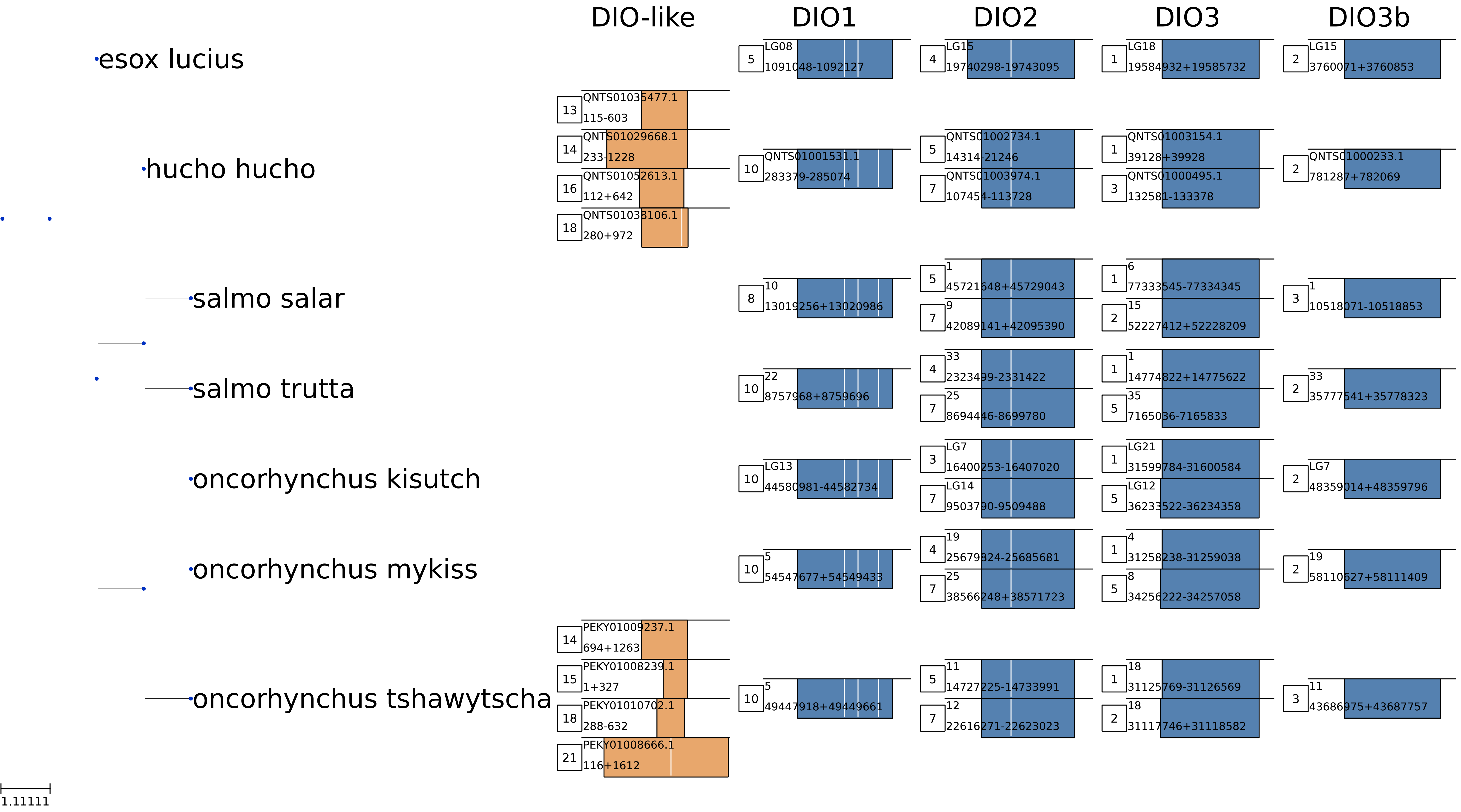

### FIGURE_S6_TXNRD.pdf

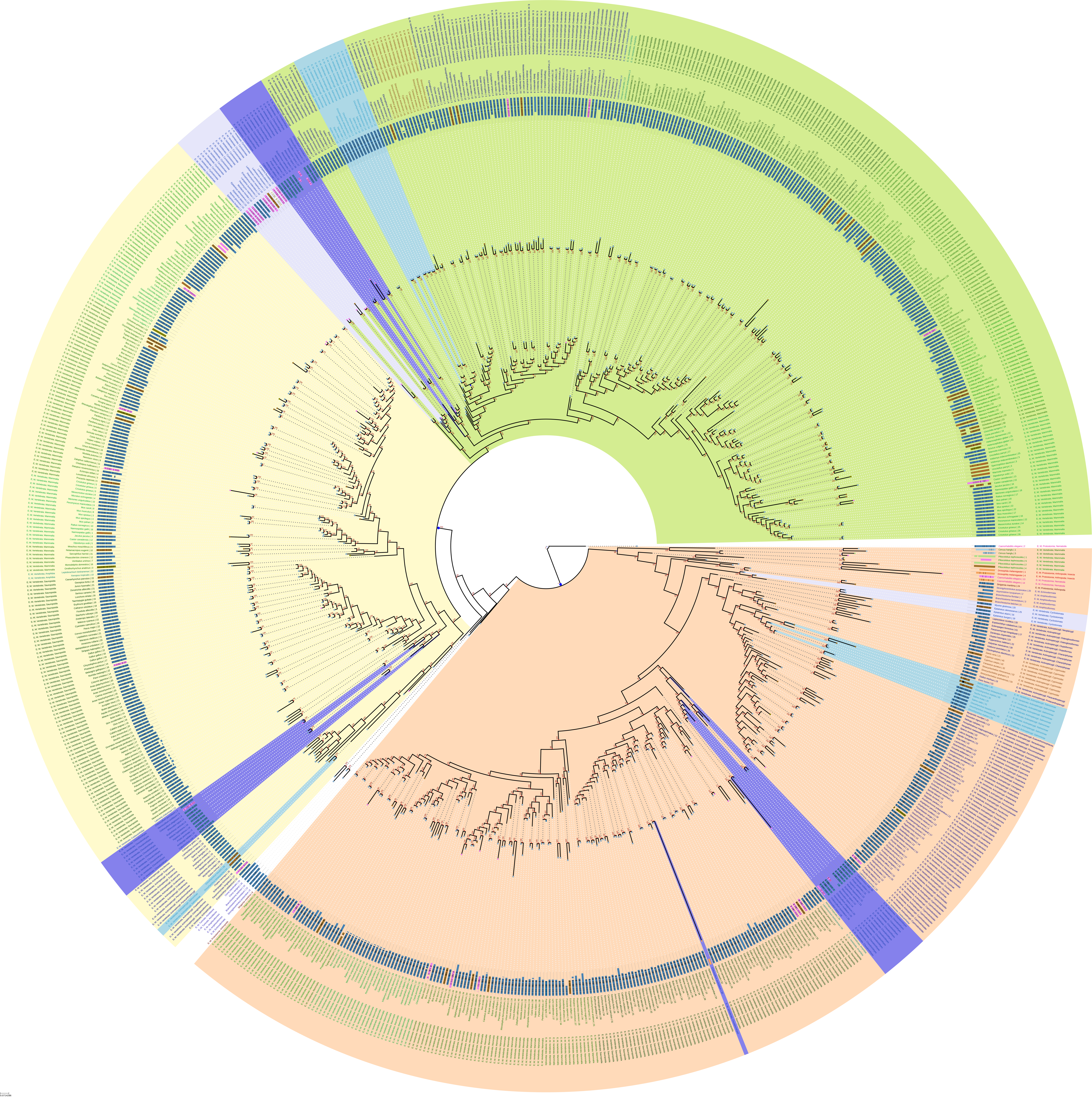

### FIGURE_S7_TXNRD1_SYNTENY.pdf

#### Synteny analysis of TXNRD1

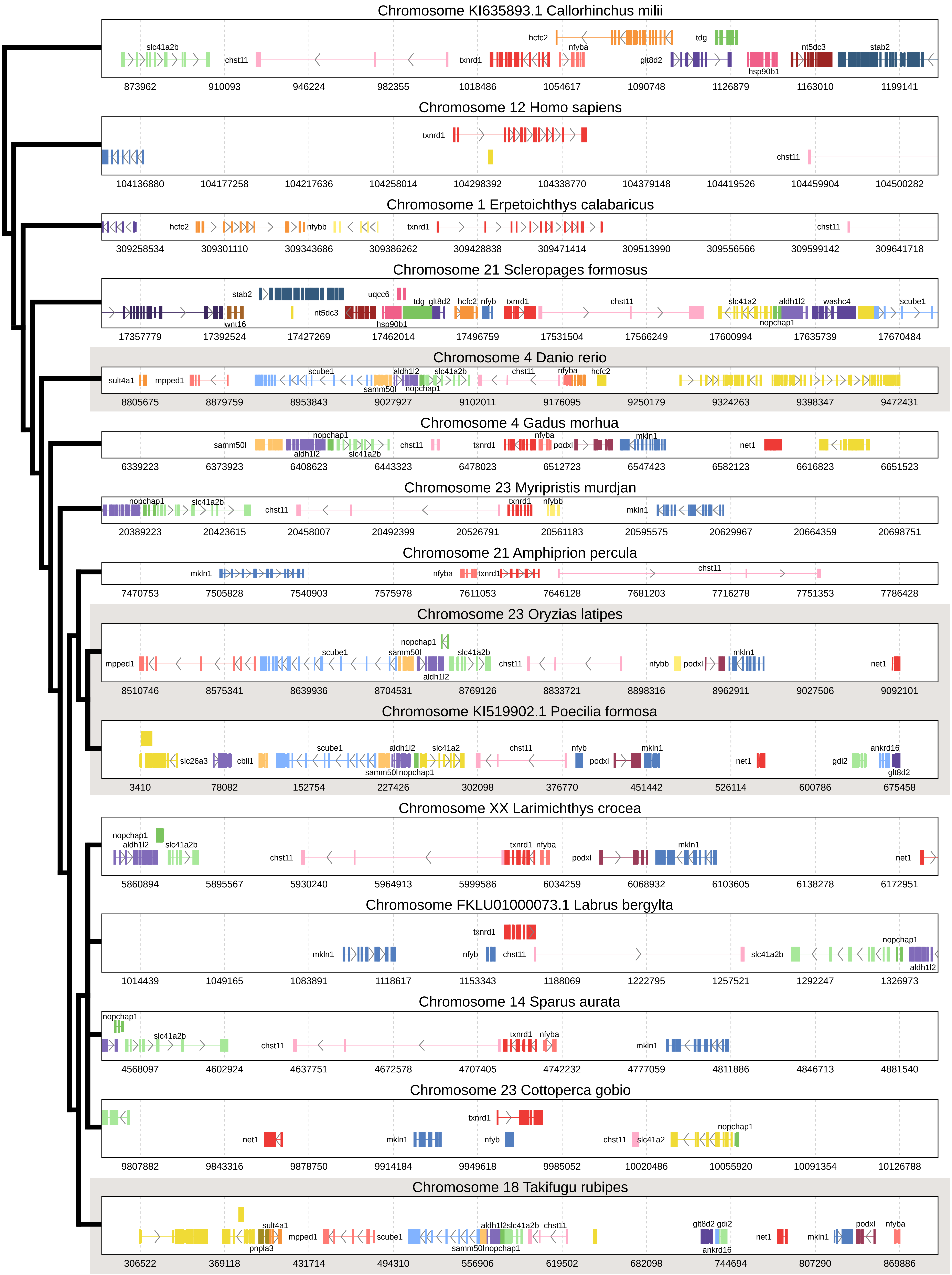

### FIGURE_S8_TXNRD3_SYNTENY.pdf

#### Synteny analysis of TXNRD3

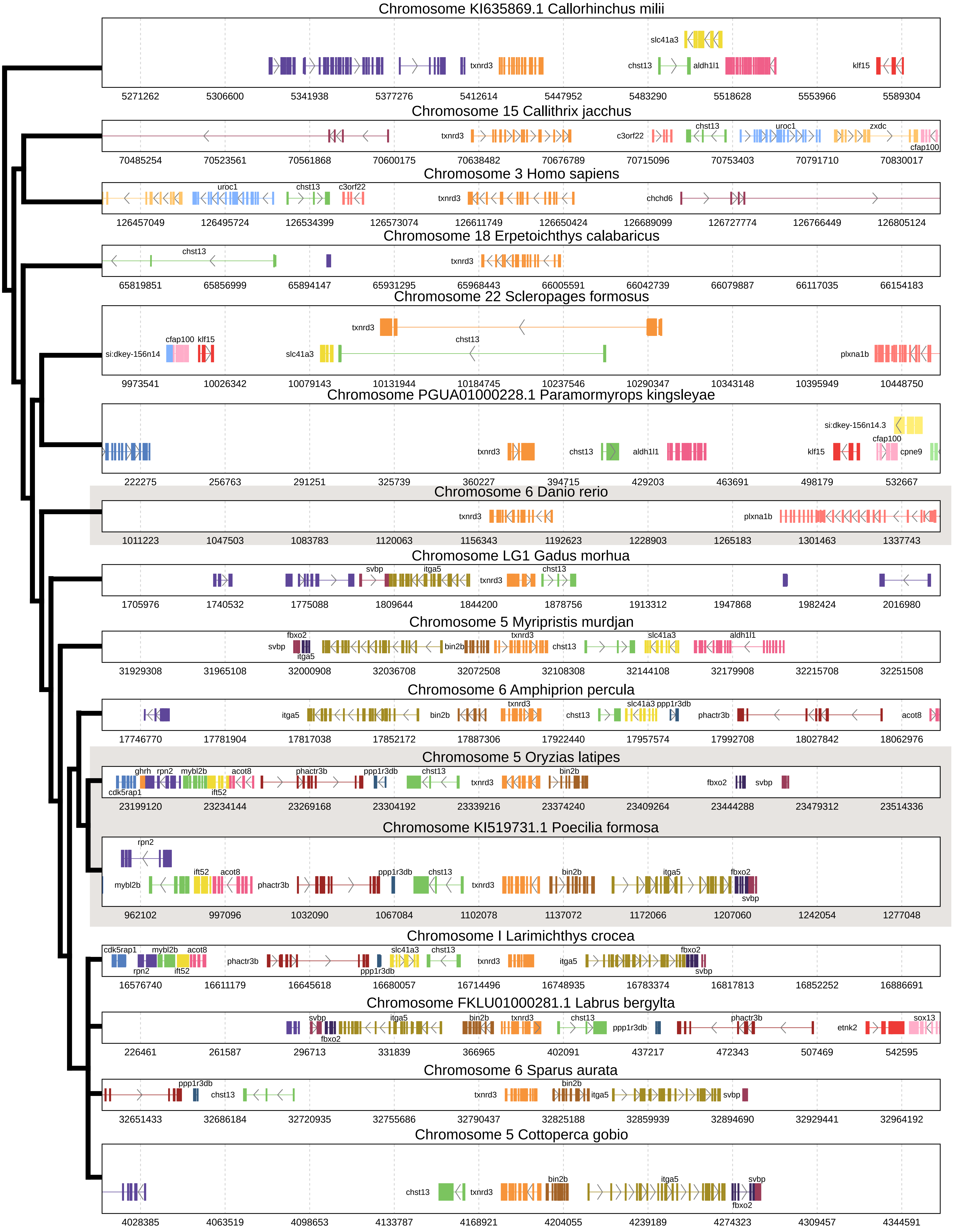

### FIGURE_S9_GRX.pdf

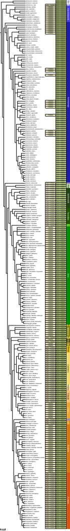

### FIGURE_S10_SELENOJ.pdf

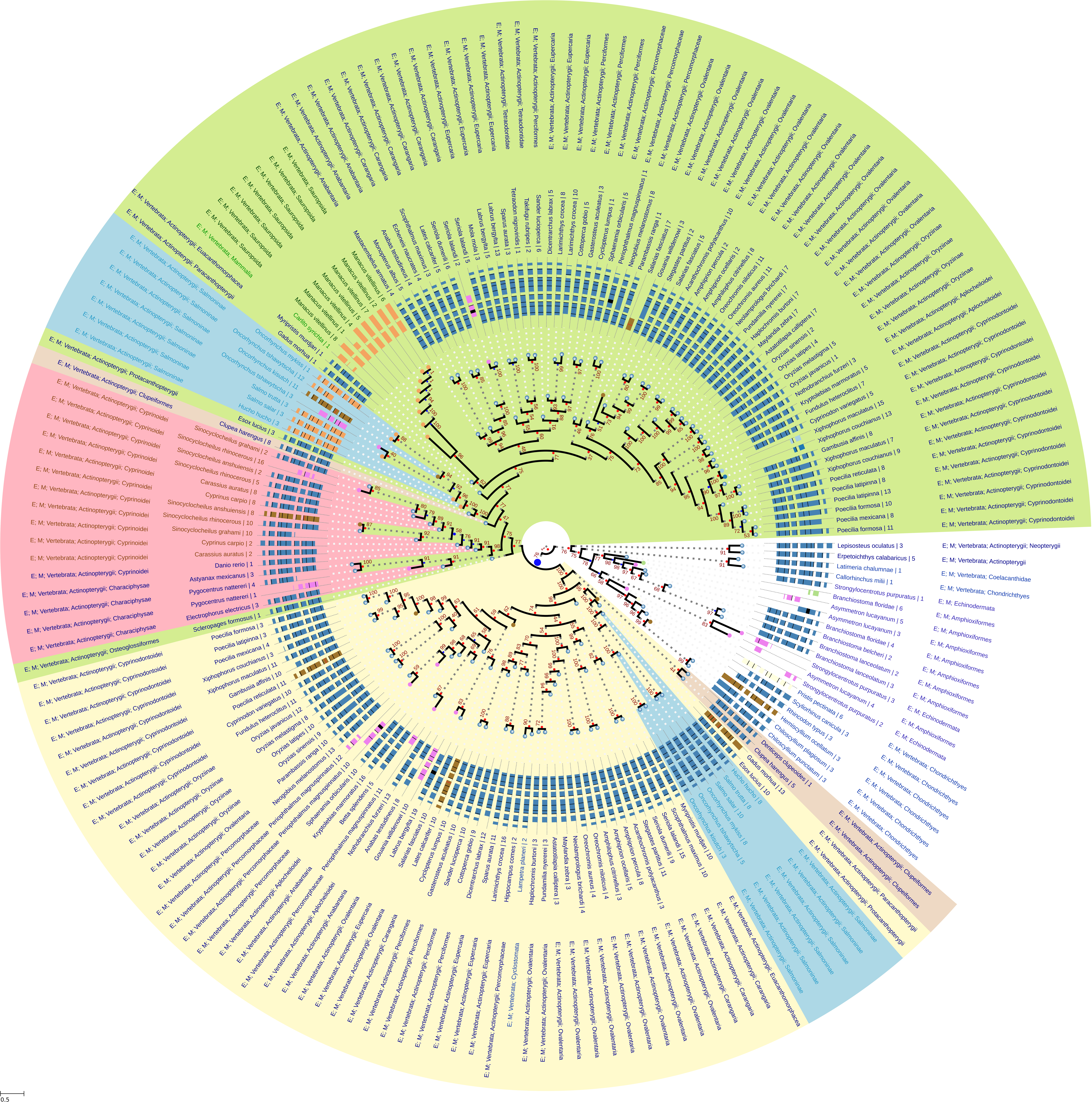

### FIGURE_S11_SELENOJ_DRAWER.pdf

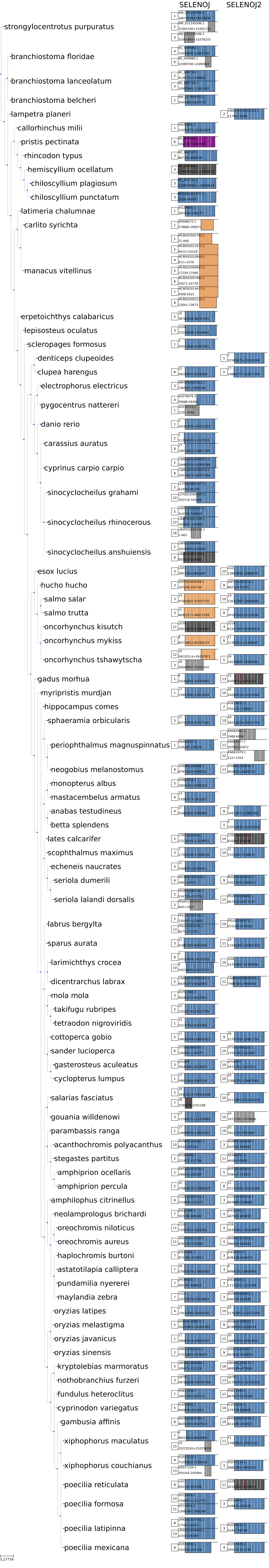

### FIGURE_S12_SELENOO.pdf

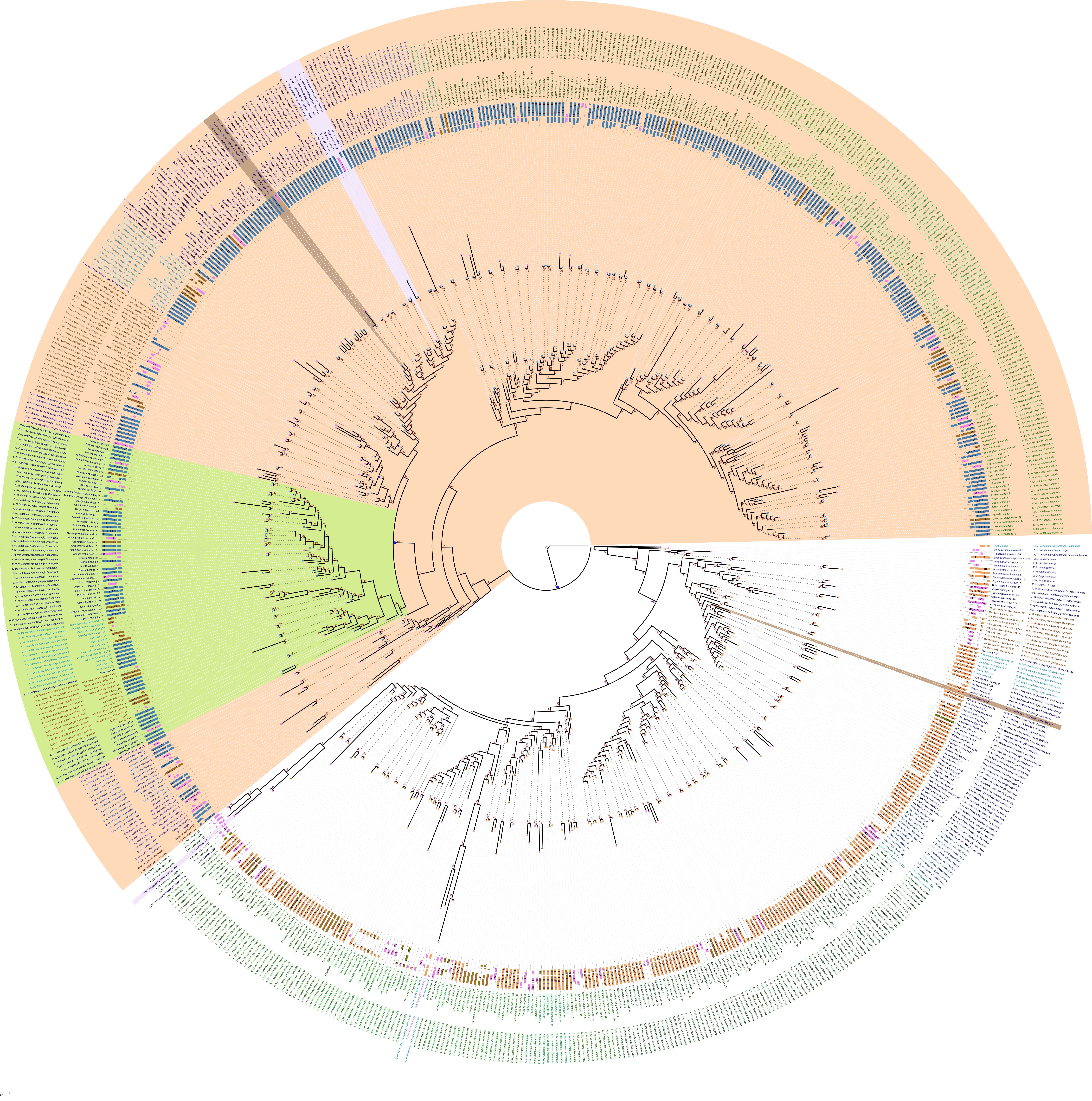

### FIGURE_S13_SELENOO_DRAWER.pdf

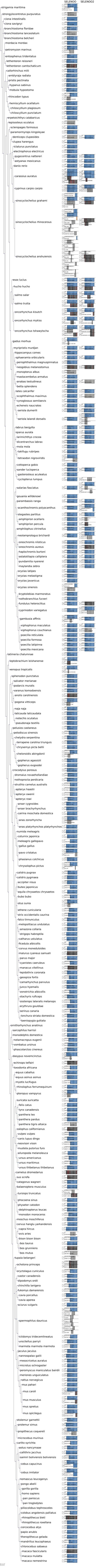

### FIGURE_S14_MSRB.pdf

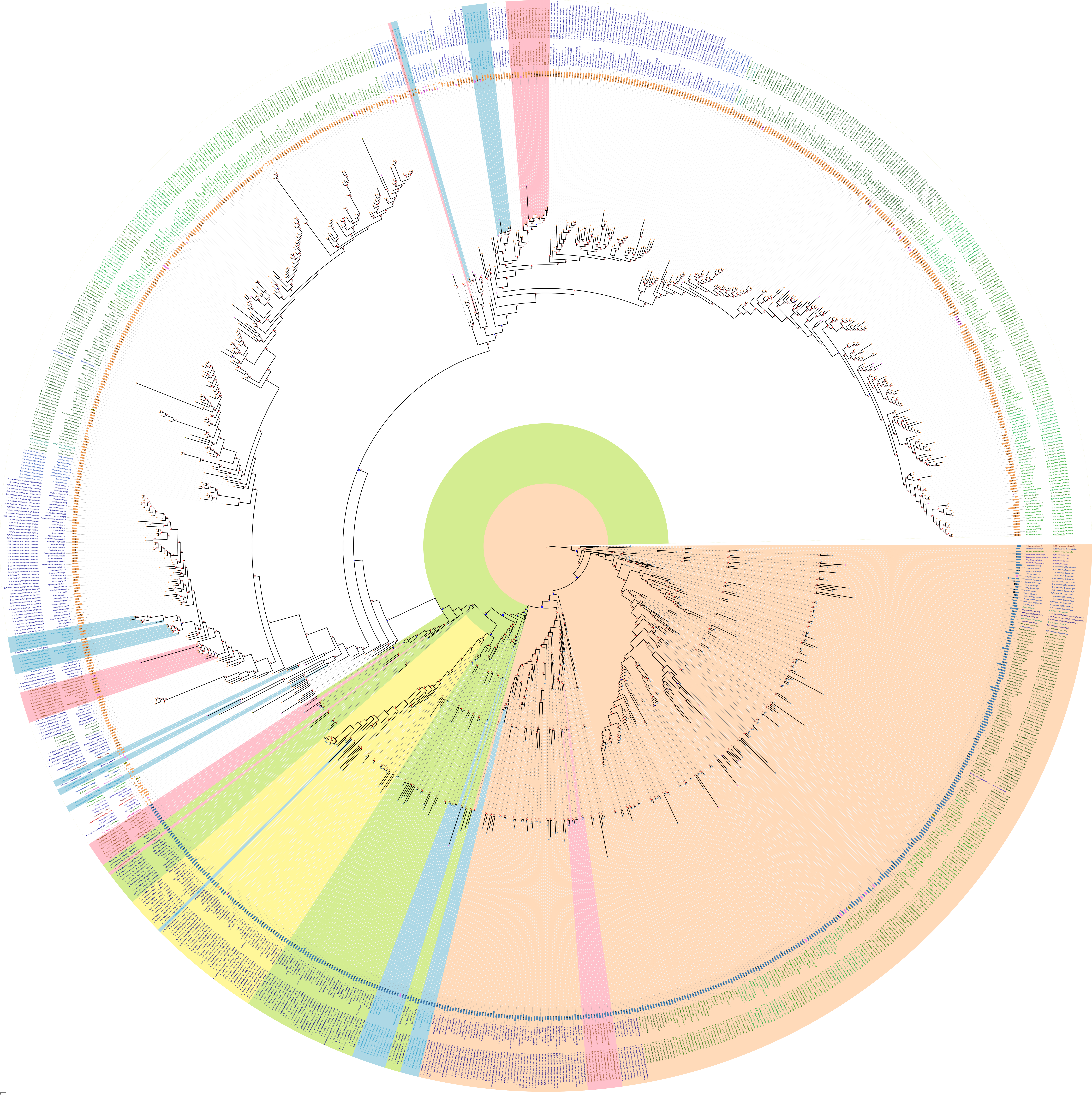

### FIGURE_S15_MSRB_DRAWER.pdf

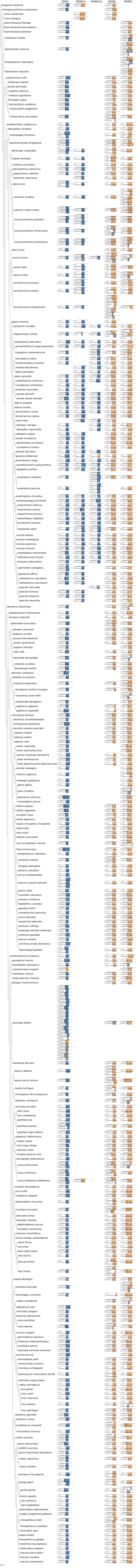

### FIGURE_S16_SELENOT.pdf

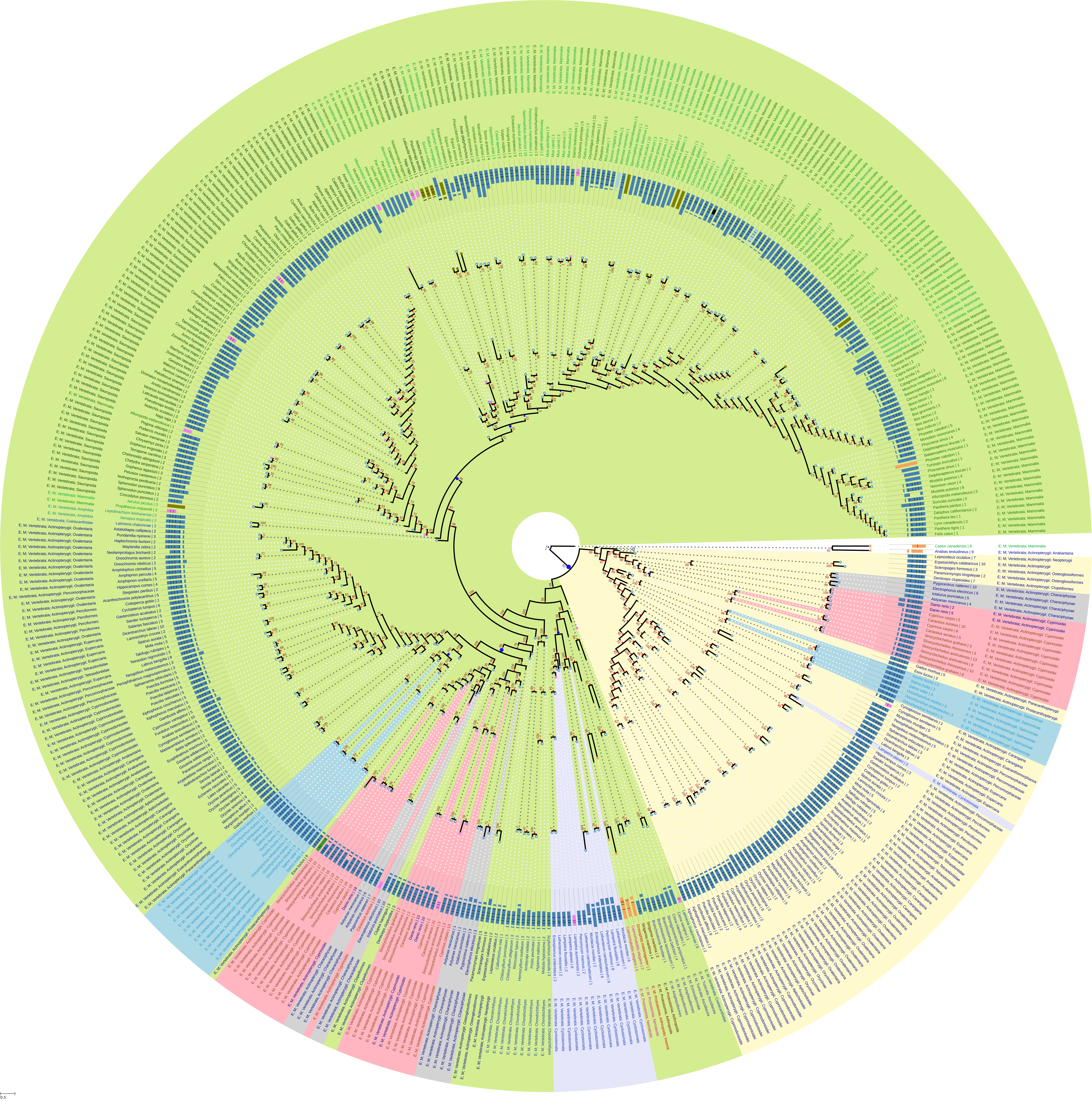

### FIGURE_S17_SELENOU.pdf

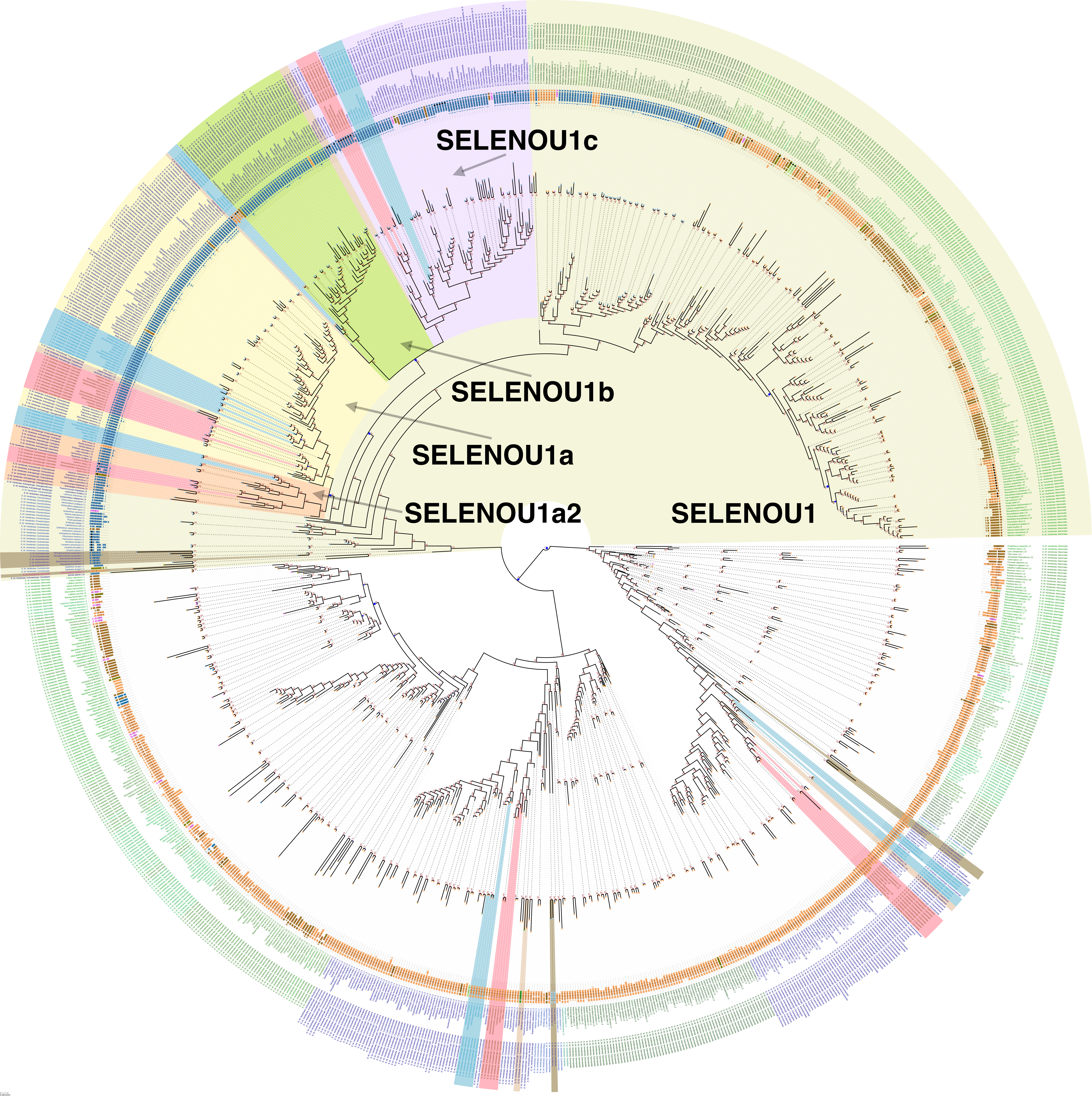

### FIGURE_S18_SELENOW.pdf

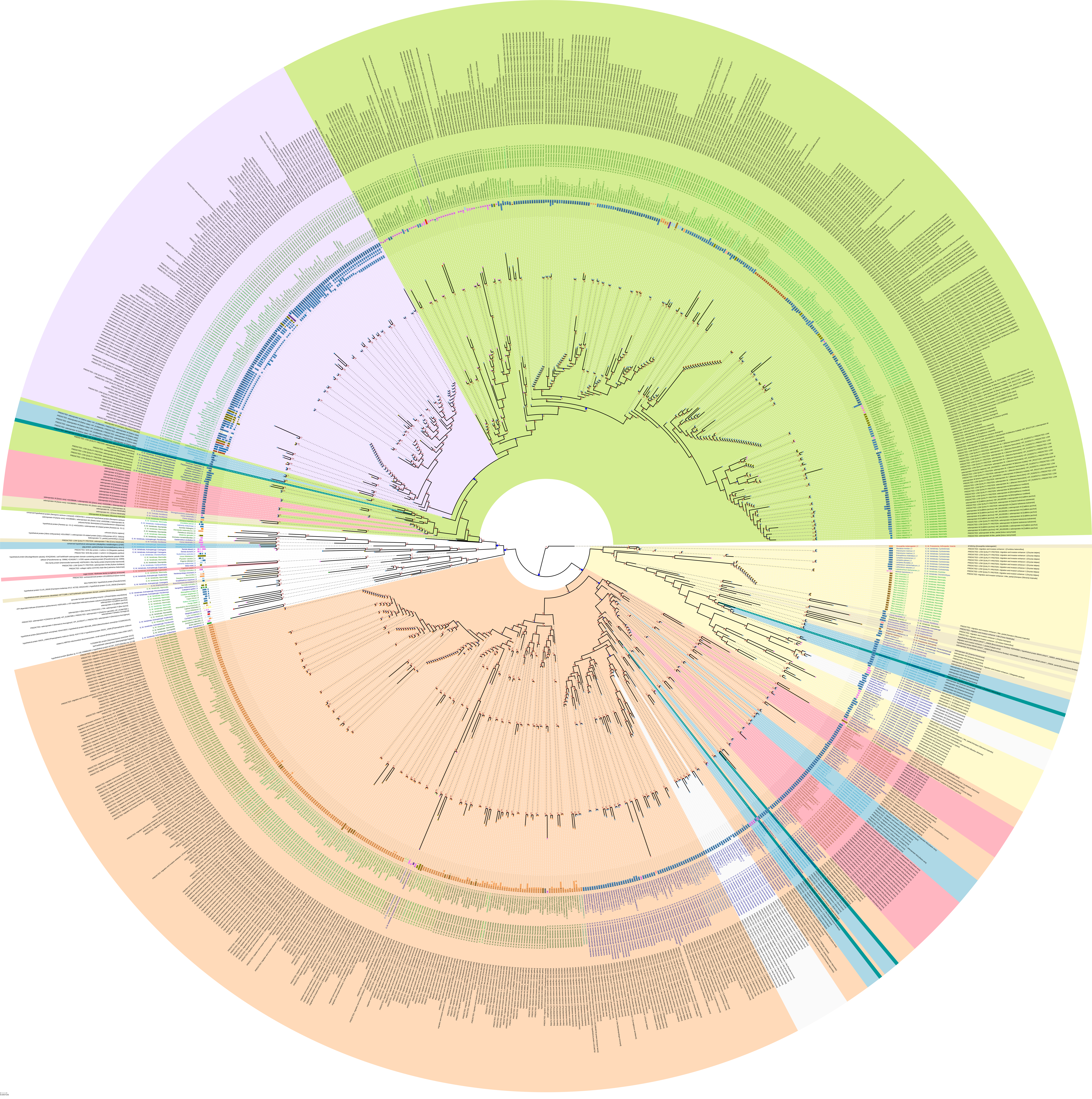

### FIGURE_S19_SELENOW1_SYNTENY.pdf

#### Synteny analysis of SELENOW1

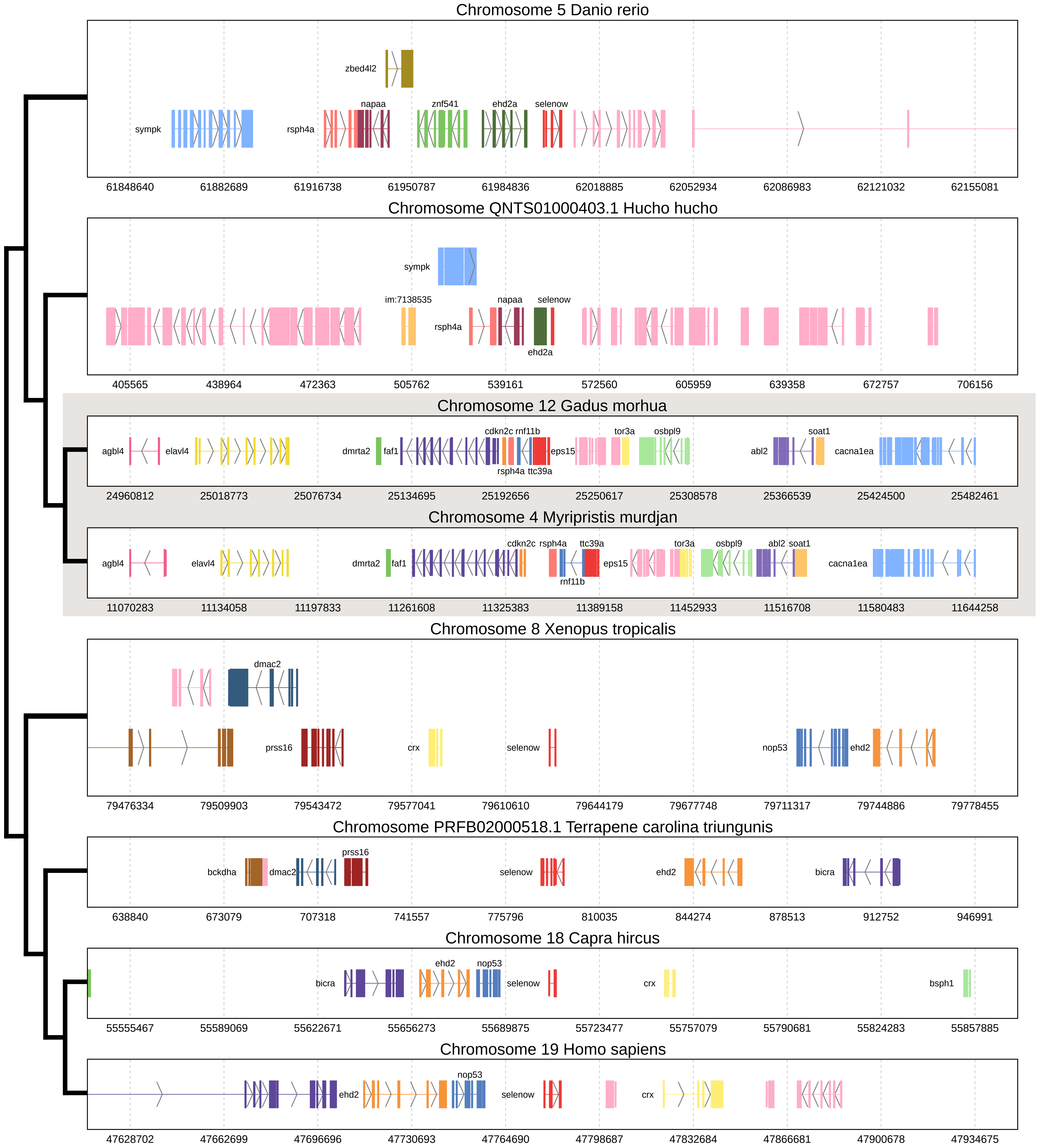

### FIGURE_S20_SELENOW2_SYNTENY.pdf

#### Synteny analysis of SELENOW2

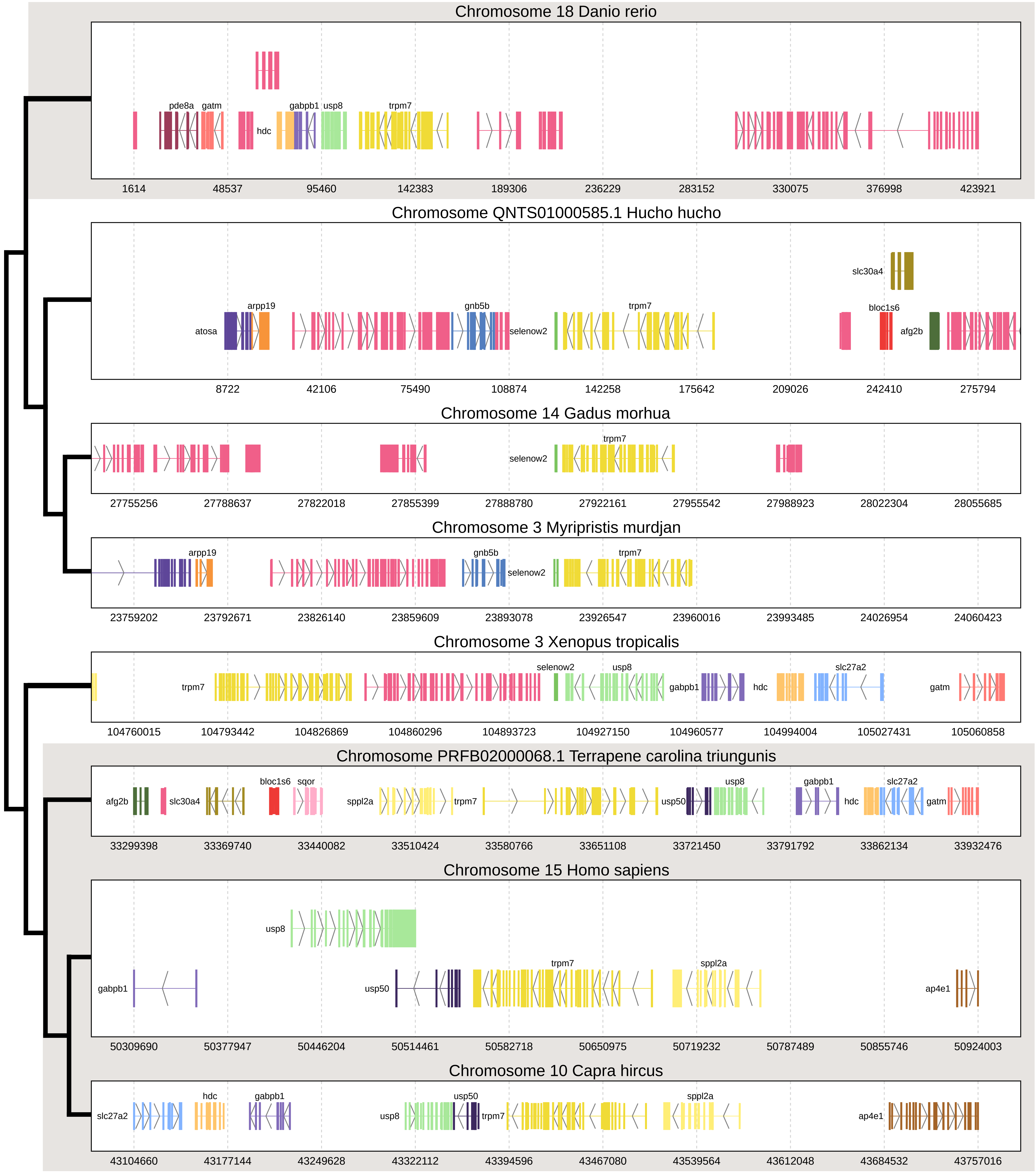

### FIGURE_S21_MIEN1_SYNTENY.pdf

#### Synteny analysis of MIEN1

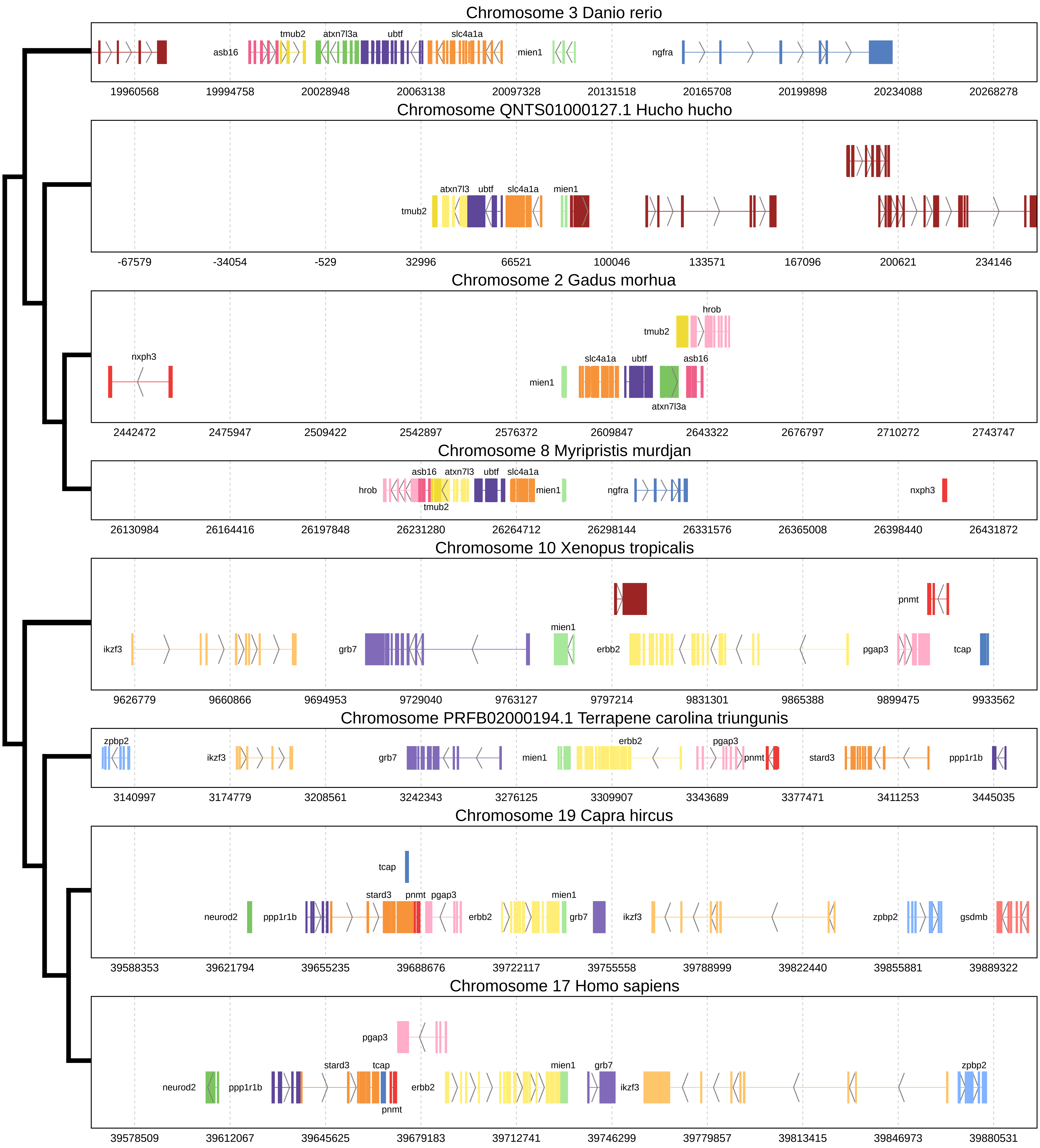

### FIGURE_S22_SELENOP.pdf

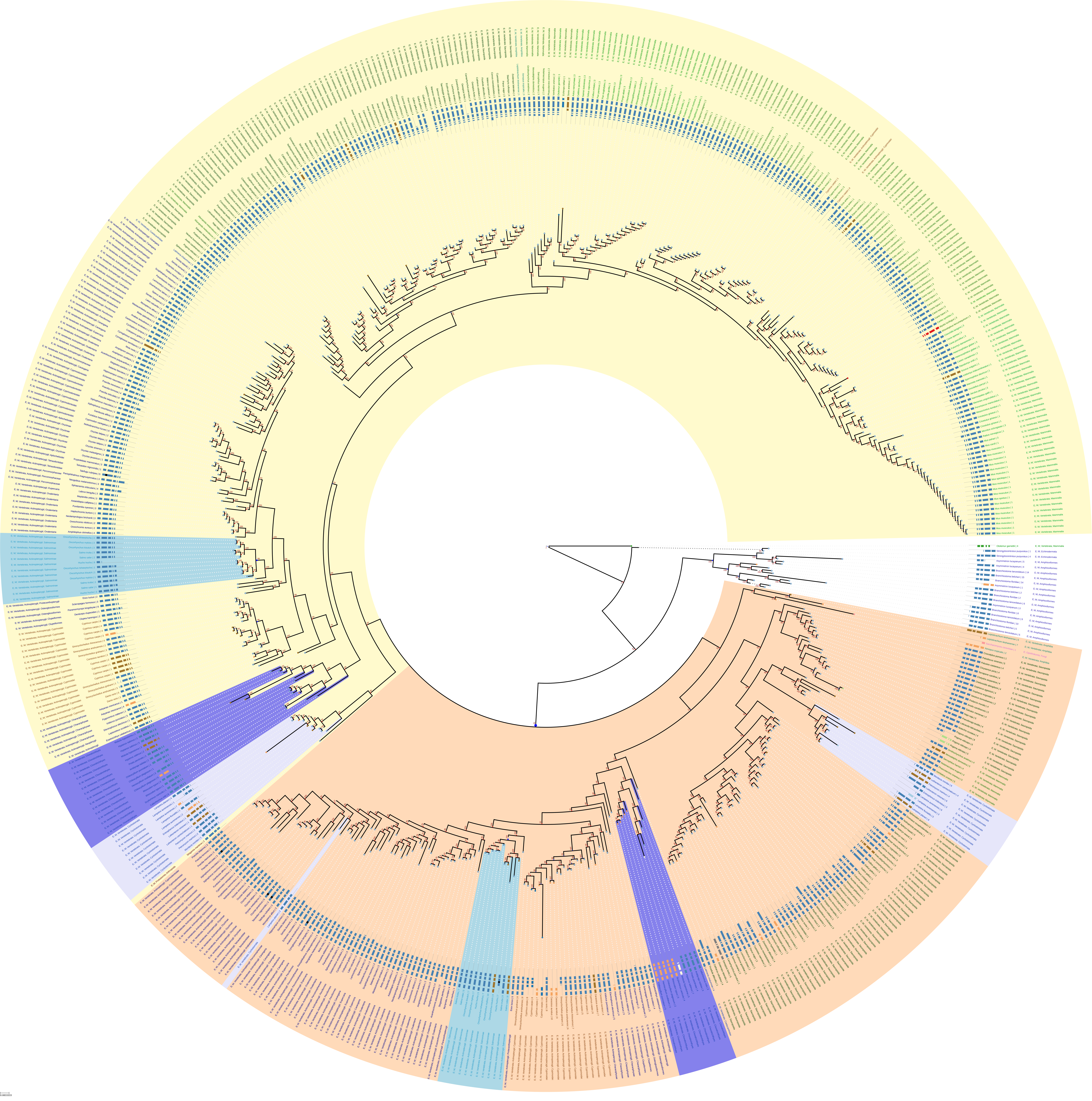

### FIGURE_S23_SELENOE.pdf

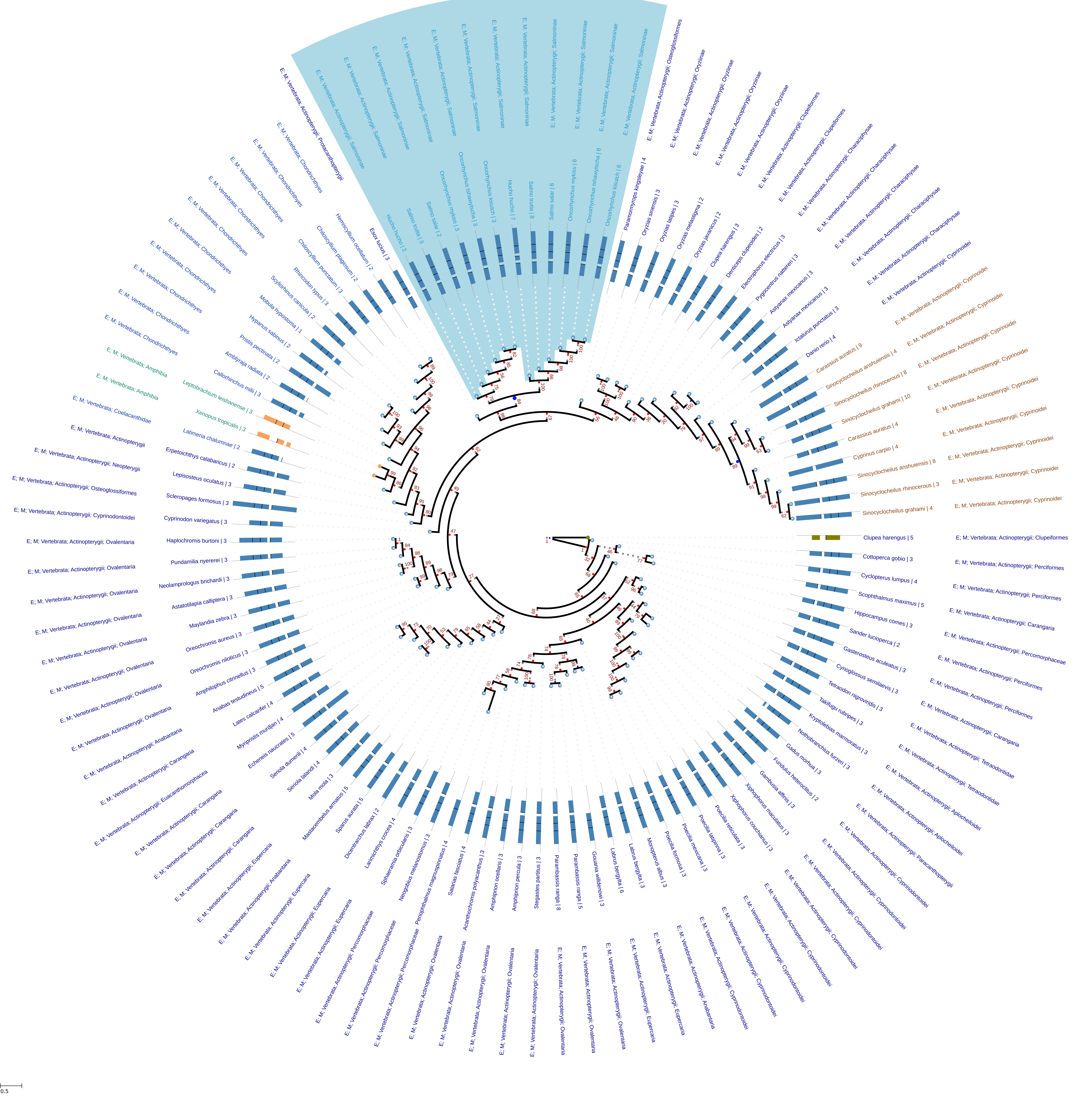

### FIGURE_S24_SELENOF.pdf

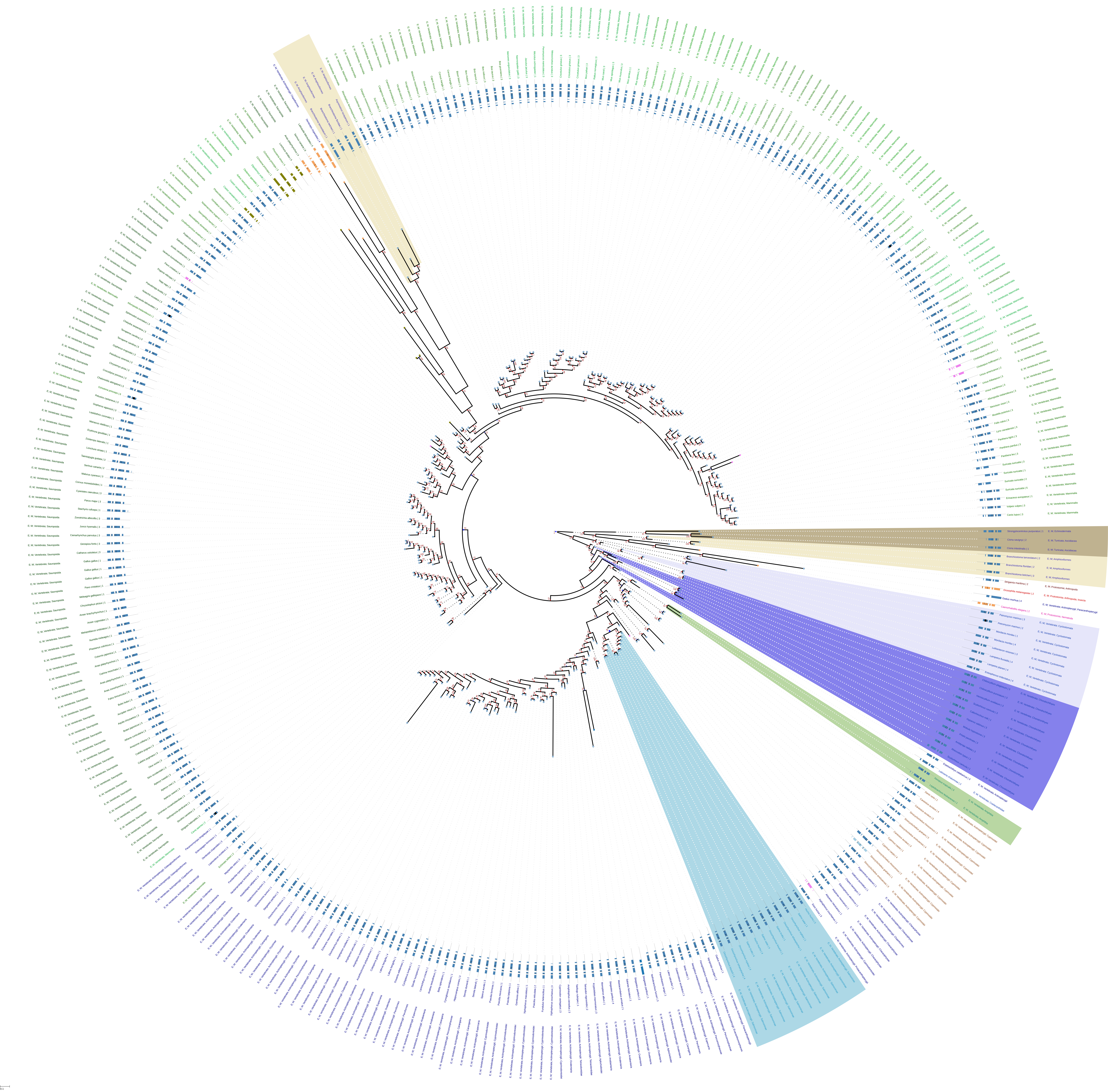

### FIGURE_S25_SELENOH.pdf

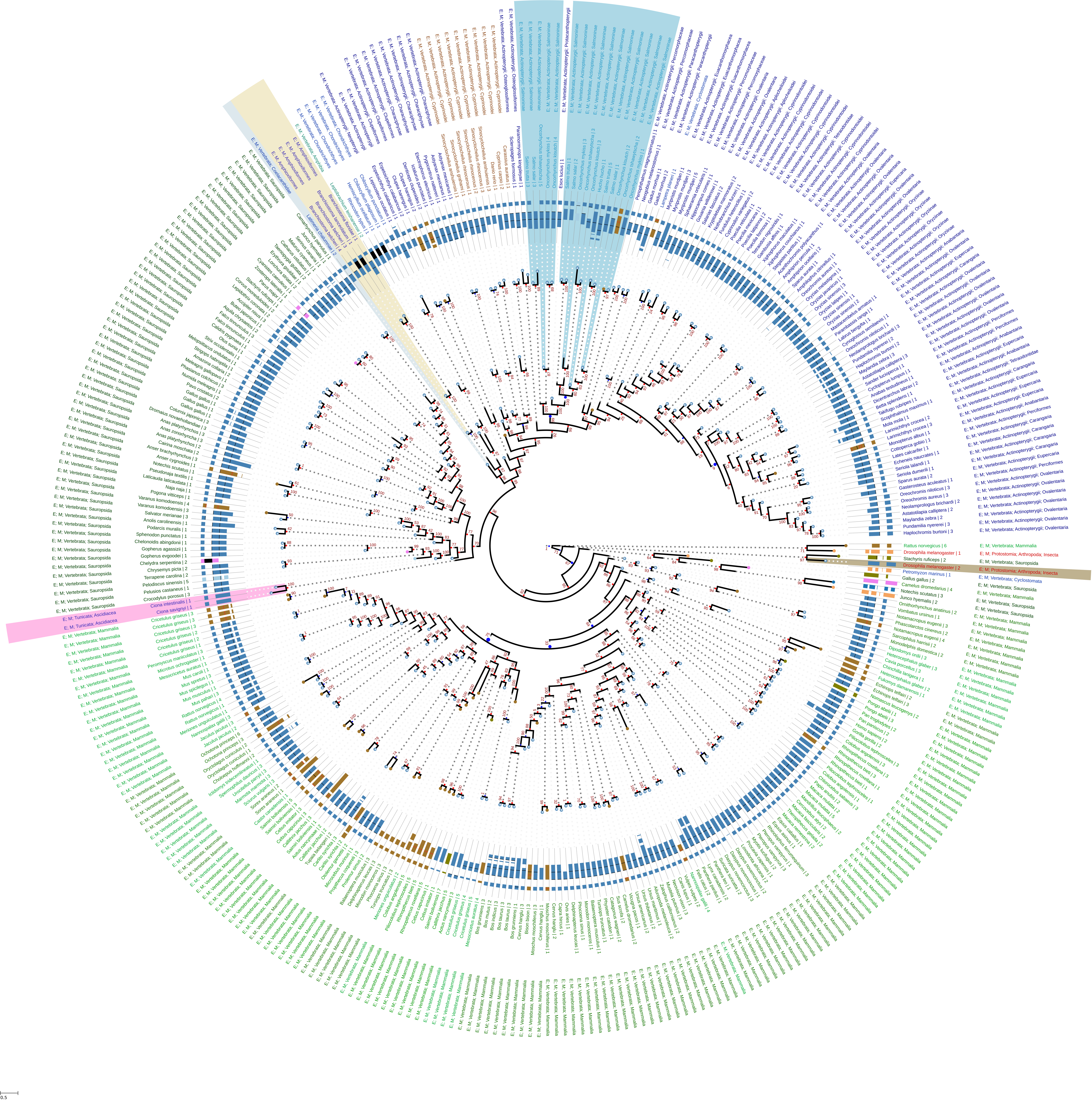

### FIGURE_S26_SEPHS2.pdf

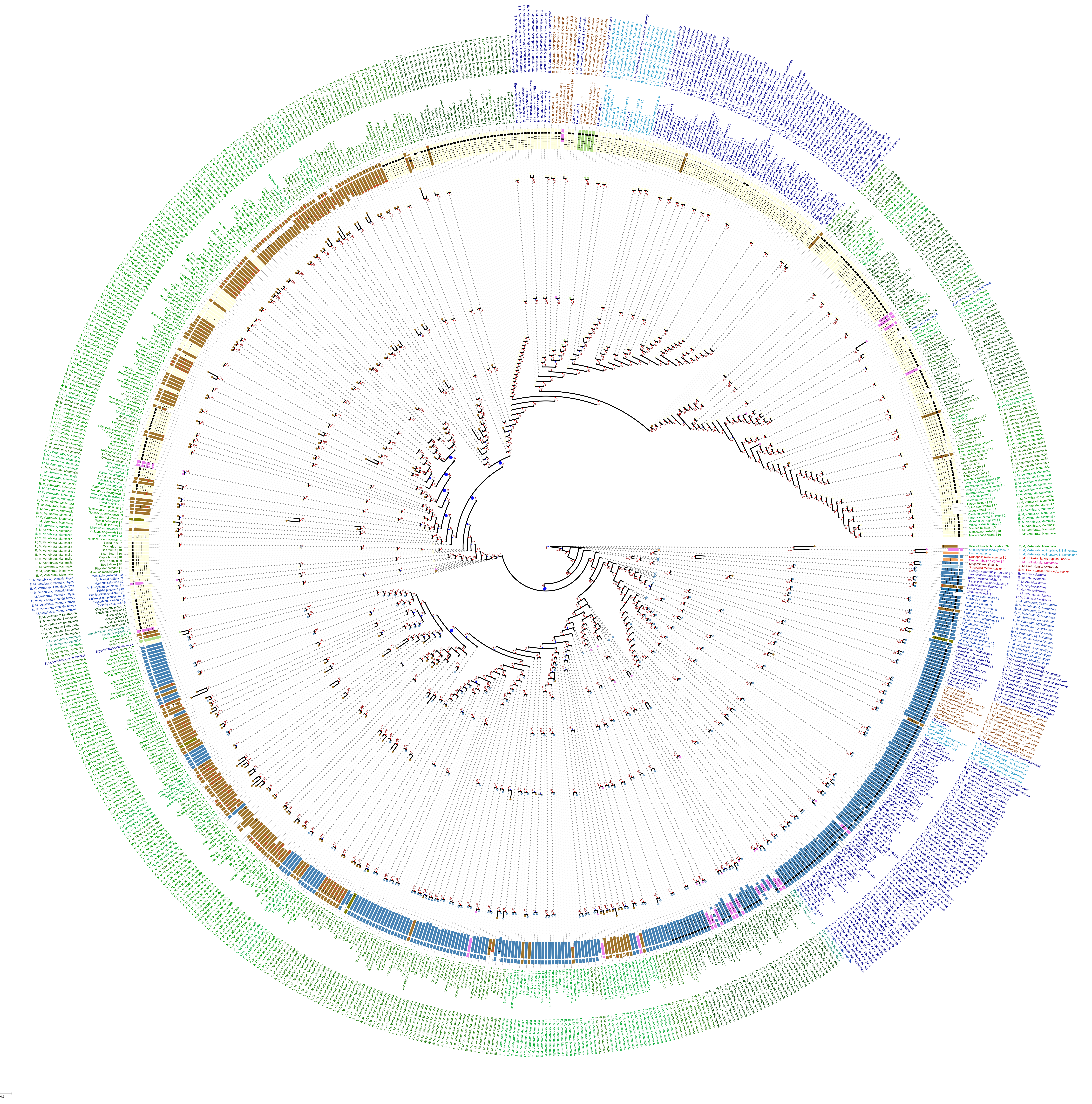
